# Screening for Polysaccharide Utilization Loci Targeting Marine Polysaccharides

**DOI:** 10.64898/2026.05.19.726164

**Authors:** William Helbert, Anthoula. Mettou, Laurent Poulet, Mélanie Loiodice, Sophie Drouillard, Marie Couturier, Aurélie Rousset, Ronan Pierre, Ahmed Khamassi, Nicola Curci, Véronique Roig-Zamboni, Gerlind Sulzenbacher, Renaud Vincentelli, Elodie Drula, Marie-Line Garron, Vincent Lombard, Youssef Bouargalne, Nushin Aghajari, Nicolas Terrapon

**Affiliations:** CERMAV, CNRS and Grenoble Alpes Université, BP53, 38000 Grenoble Cedex 9, France; Centre d’Etudes et de Valorisation des Algues, Presqu’île de Pen Lan, BP3, 22610 Pleubian, France; Centre national de la recherche scientifique (CNRS), Aix-Marseille Univ, UMR 7257 Architecture et fonction des macromolécules Biologiques (AFMB), Marseille, 13288, France; Institut national de recherche pour l’agriculture, l’alimentation et l’environnement (INRAE), USC 1408 AFMB, 13288 Marseille, France; INRAE, Aix Marseille Univ, UMR 1163 Biodiversité et Biotechnologie Fongiques (BBF), 13288 Marseille, France; Molecular Microbiology and Structural Biochemistry, UMR 5086, CNRS-Université de Lyon, F69367 Lyon, France

## Abstract

Polysaccharide utilization loci (PULs) have been a goldmine for the characterization of novel carbohydrate active enzymes (CAZymes) and the understanding of their synergistic degradation of complex polysaccharides. We collected PUL predictions containing CAZymes from glycoside hydrolase families GH29, GH50 and GH117, expected to participate in marine polysaccharide breakdown. We explored the evolutionary diversity in these families in terms of sequences and PUL composition, based on sulfatases and CAZymes. From 41 selected PULs, more than 400 putative enzymes were produced, purified and screened on a large collection of carbohydrates. We attributed a function to more than 130 enzymes from five sulfatase subfamilies, 29 known CAZymes families and discovered an activity for 4 families previously of unknown function, including an α-L-galactosidase structurally and functionally characterized with mutants. Finally, our detailed analysis of the enzymatic synergies in five PULs, two targeting marine polysaccharides and three targeting eukaryotic polysaccharides, by marine and human gut organisms, highlight the efficiency of our exploratory strategy.

## Introduction

Marine polysaccharides represent the most abundant and the most diverse marine biomass^1,2^. These later are frequently decorated with sulfate esters, which are widely assimilated by marine organisms^3,4^ due to the high concentration of inorganic sulfate ions in seawater (25 to 28 mM) compared to its concentration in freshwater or soil (10–50 μM)^5^. The diversity of marine sulfated polysaccharides reflects the diversity of evolutionary lineages found in marine environments^6,7^. Notably, characteristic sulfated polysaccharides are extracted from the cell wall of red (e.g., agars, carrageenans)^8^, brown (e.g., fucans, fucoidan)^9^ or green algae (e.g. ulvans, cladophorans)^10^. Many other sulfated polysaccharides whose structures are not fully elucidated are biosynthesised by microalgae, diatoms, bacteria and animals^11,12^.

Degradation of marine sulfated polysaccharides requires the concerted action of enzyme cocktails comprising glycoside hydrolases (GH) and polysaccharide lyases (PL), both able to cleave glycosidic bonds, and sulfatases that catalyse the removal of sulfate ester groups. Bacteria of the phylum Bacteroidota are well-known polysaccharide degraders^13,14^. They have evolved sophisticated systems for the efficient breakdown of glycans in which the enzymes involved in the same polysaccharide degradation pathway are encoded by colocalized genes in so-called *polysaccharide utilization loci* (PUL)^15^. The typical PUL is organized around a tandem of genes homologous to *susC* (oligosaccharide transmembrane transporter) and *susD* (carbohydrate receptor) along with genes encoding the appropriate enzymes as well as regulators^13,14^. PULs may additionally encode proteins of yet unknown function, which might be novel enzymes participating in glycan uptake, awaiting biochemical characterization.

The study of PULs in marine bacteria helped to elucidate all the degradation pathways of several algal polysaccharides, such as carrageenan^16,17^ agars^18,19,20^, and ulvan^21,22^. However, the ability to digest algal polysaccharides is not restricted to marine organisms but was also reported in species of the human gut with reports of the degradation of agars^18,23,24^, carrageenan^19^, alginates^25^ and chaetomorphan^26^. This could be explained by the lateral transfer of genetic material of bacteria living at the surface of the algae used in diet^23^, shedding light on the important relationships between humans and ocean.

The localization of PULs involved in the degradation of a given polysaccharide could be highlighted by transcriptomic experiments^27,28^. However, such identification requiring cultivable strains and important sequencing costs, PULDB (www.cazy.org/PULDB/) was developed to propose PUL predictions based on genomic data only^29^, taking advantage of the high-quality annotations of the carbohydrate-active enzymes database (CAZy: www.cazy.org/)^30^. The selection of a particular PUL for biochemical studies then relies on its hypothetical substrate, among known polysaccharide structures, a hypothesis based on the known specificity of the GHs, PLs and sulfatase families composing each PUL. Improving the knowledge on enzyme specificities is thus essential and requires the biochemical exploration of families. These last years, medium throughput screening using collections of oligo- and polysaccharides initiated a more systematic exploration of sequence diversity within CAZy families^25,31,32^. In the same vein, screening activities of PUL proteins distantly related to CAZymes and of unknown function, led to the discovery of new CAZy families and specificities^26,33,34^.

In this context, we proposed a strategy combining bioinformatics exploration of CAZy and PULDB data, with functional screening assays to attribute an activity to enzymes co-located in PULs and, *in extenso*, a substrate to these PULs. Our strategy was based on the rational selection of PULs potentially active on marine polysaccharides, as composed of sulfatases and some CAZy families known to be active on marine polysaccharides: GH29, GH50 and GH117. These families mostly contain α-L-fucosidases (essential for fucan/fucoidan breakdown), β-agarases and α-L-anhydrogalactosidases (anhydro-L-galactose being only reported in agars so far), respectively. The genes encoding more than 400 selected targets, belonging to 41 PULs, were synthesized and their recombinant proteins were screened on a wide diversity of substrates. The results led to an increase of biochemically characterized enzymes highlighting polyspecificity in GH29, GH50 and GH117 families. It also led to the discovery of new families and new degradation pathways for marine polysaccharides by marine, but also, gut bacteria.

## Results

### 1. Selection and production of PUL enzymes

The strategy for the rational selection of enzymes to characterize started with the identification of PULs putatively involved in the degradation of marine polysaccharides. Among the ∼31 thousands predicted PULs across ∼1000 genomes in PULDB (June 2018), less than 6% of them combine GH, PL and sulfatases, still representing more than 1800 PULs potentially involved in the degradation of sulfated polysaccharides^35^. We hence focused on CAZy families known to be frequently found in PULs and to encompass marine polysaccharide degrading enzymes. Our subset of relevant families included: GH50, that contains characterized endo-β-agarases; GH117, with reported α-L-anhydrogalactosidases that cleave residues specifically found in agars; and GH29, known for α-L-fucosidases essential for the breakdown of fucans and fucoidans found in the cell wall of brown algae. All PULs containing at least one member of these three families were extracted from the PULDB database. The amino-acid catalytic domain sequences were used to calculate phylogenetic trees of the GH29, GH50 or GH117 PUL members. Leaves were labelled with the CAZymes and sulfatases composing the cognate PUL (Fig. S1-3). While based on amino-acid sequences only, these trees also group PULs with similar compositions.

Based on GH29, GH50 and GH117 phylogenies, we selected 36 PULs that covered a large diversity in terms of sequences and PUL compositions, hypothesized to reflect variability in substrate composition, structure or decoration. We completed our selection with five PULs containing both sulfatase subfamilies of interest and members of the GH127 family that includes α-D-anhydrogalactosidases found in carrageenan catabolism. The final selection totalizes 41 PULs (Table S1) and is not restricted to marine bacteria (23 PULs) but also includes several species from gut microbiota (14 PULs) which could target sulfated polysaccharides originating from either algae, other organisms in the diet, or the host itself^19,23,25^. In these PULs, we selected ∼230 genes encoding GHs, ∼20 PLs, ∼120 sulfatases and ∼120 proteins with unknown function (Table S1).

These genes were synthesised and cloned in expression vectors that encoded His-tag for purification. Expression assays were conducted at the microplate scale and protein production for screening assays were up-scaled in 50mL culture medium. According to our expression conditions, the yield of soluble proteins was 62.5%. The yield of soluble enzymes selected from the gut microbiota was 87.3% (n=100) while only 51.7% (n=144) for proteins of marine strains (Fig. 1A). CAZymes were better expressed soluble (68%, n=183) than the sulfatases (43%, n=22); expression of proteins of unknown function were obtained with an intermediate yield (58%, n=56) (Fig. 1B). The high expression yield of enzymes selected from gut bacteria could be explained by the shared ecosystem with the *E. coli* expression system. In contrast, marine enzymes represent a challenge for soluble expression, as previously observed^36^. Activity screening of CAZymes, sulfatases and proteins of unknown function were conducted at 96-well microplate scale using a library of substrates including pNP-glycosides, oligo- and polysaccharides following the same strategy as previously implemented^26^ and detected enzyme activities were reported (Table 1, 2 and S2).

**Figure 1:**
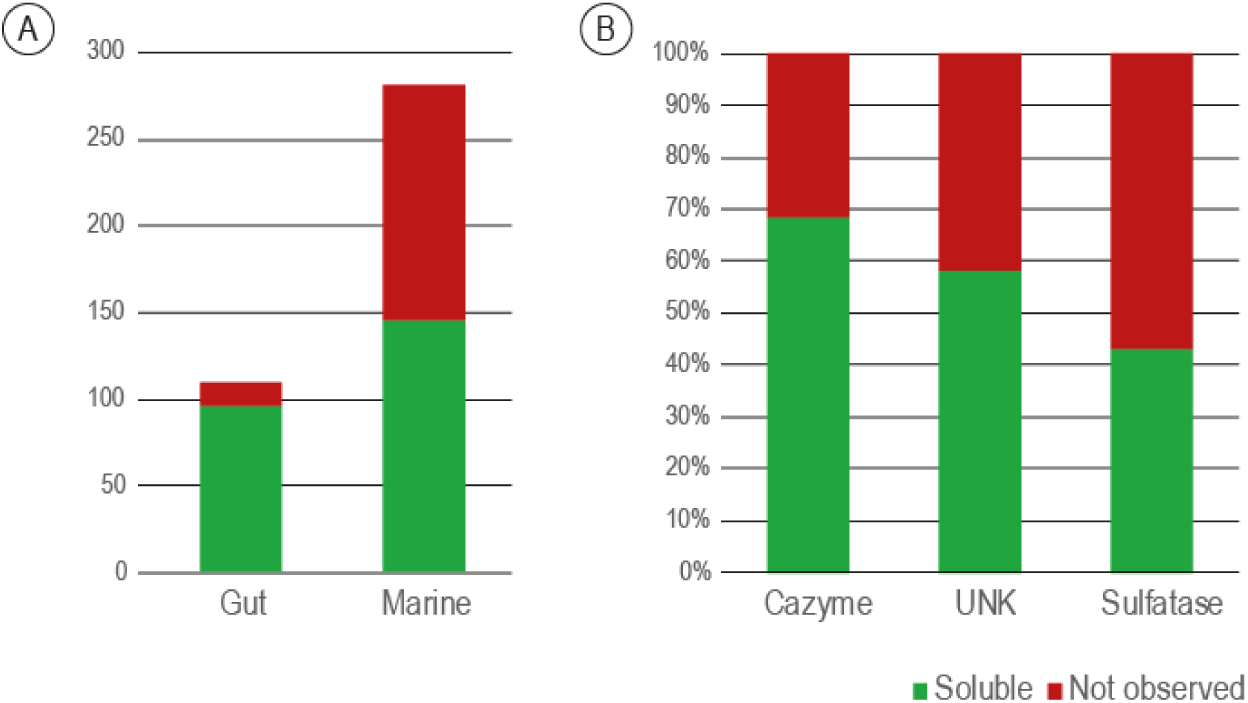
Protein expression results. **A**) Comparison of the protein expression between protein selected in gut microbiota and in marine organisms. **B**) Comparison between enzymes as a function of their classification: CAZymes (glycoside hydrolases and polysaccharides lyases), formylglycine dependant sulfatase and unknown protein. “Soluble” refers to overexpressed proteins purified by nickel-affinity chromatography and detected using gel electrophoresis; “not observed,” to proteins that did not bind to any affinity column (e.g., inclusion bodies, misfolded proteins).

**Table 1:**
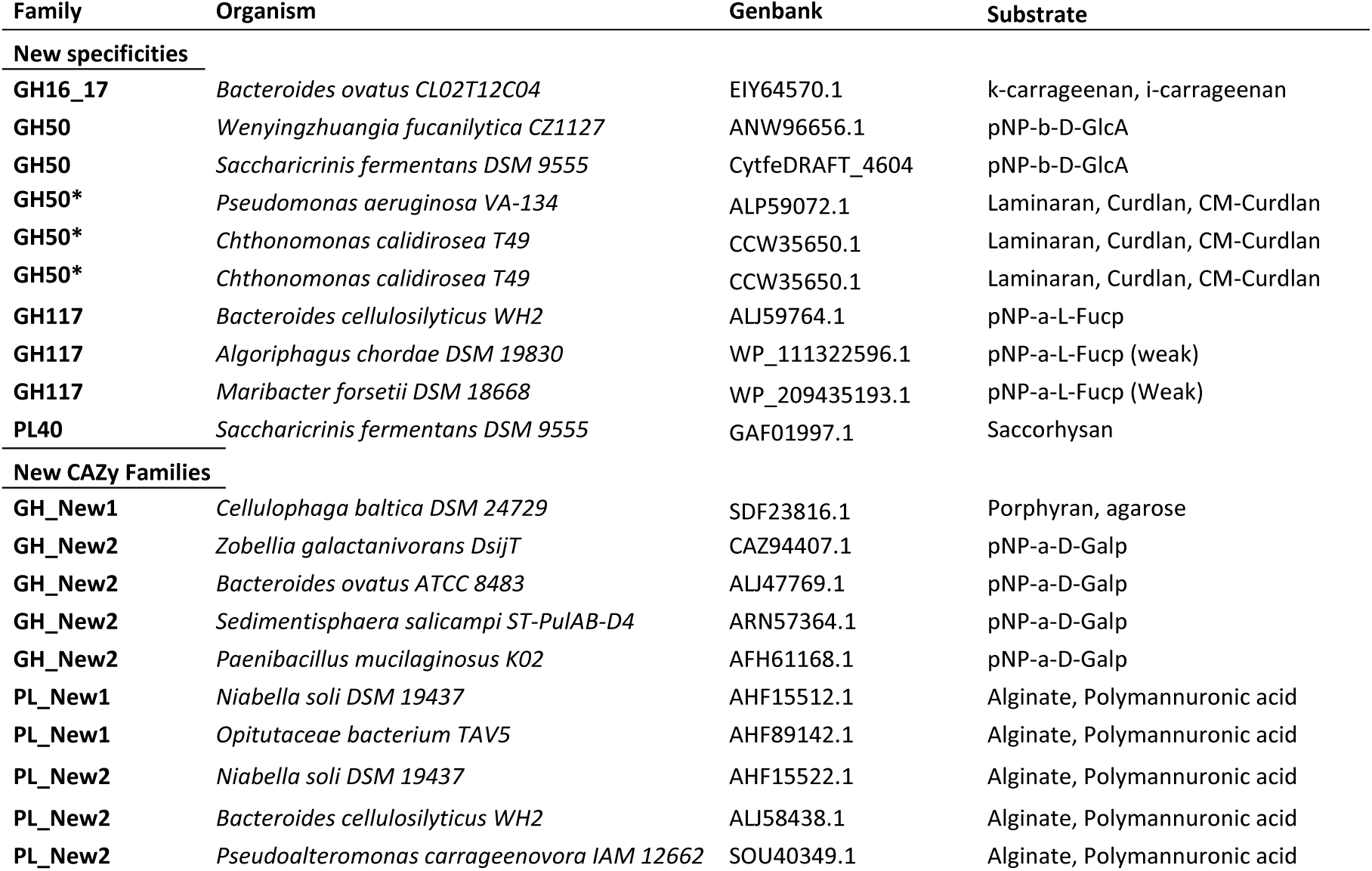
List of new enzyme specificities discovered in established CAZy families and of new CAZy families.

**Table 2:** Function assignment to 5 sulfatase subfamilies.

| Family | Organism | Genbank | Activity | Comments |
| --- | --- | --- | --- | --- |
| **S1_9** | *Bacteroides stercoris* CC31F | EPH20505.1 | Δ4,5-hexuronate-2-O-sulfate 2-O-sulfohydrolase | |
| **S1_9** | *Bacteroides cellulosilyticus* WH2 | ALJ59766.1 | Aryl-sulfatase |  |
| **S1_16** | *Bacteroides ovatus* CL02T12C04 | EIY64579.1 | Exo-4S-k-carrageenan sulfatase | **New specificity** |
| **S1_30** | *Bacteroides ovatus* CL02T12C04 | EIY64557.1 | Exo-4S-i-carrageenan sulfatase | **First member** |
| **S1_30** | *Bacteroides ovatus* CL02T12C04 | EIY64575.1 | Exo-4S-i-carrageenan sulfatase | **First member** |
| **S1_46** | *Bacteroides dorei* DSM 17855 | EEB26054.1 | Aryl-sulfatase |  |
| **S1_81** | *Bacteroides ovatus* CL02T12C04 | EIY64580.1 | Exo-2S-a-carrageenan sulfatase |  |

### 2. Exploration of GH29, GH50 and GH117

The selection of enzymes from families GH29, GH50 and GH117 offered an opportunity to explore the functional diversity in these families (Fig. S4-S6).

The GH50 family contained almost exclusively β-agarases but one β-1,3-glucanase^37^ and one β-galactosidase^38^ were also characterized. We selected eleven PUL members and obtained nine soluble proteins (82%). We were only able to detect an activity for three members: an exo-β-neo-agarobiosidase (BTO15_18125) and two β-D-glucuronidases (CytfeDRAFT_4604 and AXE80_10390), confirming polyspecificity in this family (Fig. S4). To better cover the sequence diversity and to confirm the polyspecificity, we selected and screened 13 additional GH50 enzymes, beyond the Bacteroidota phylum (Table S3). This led to the confirmation of rare activities in the GH50 family, with two endo-β-1,3-D-glucosidases (ATC05_19450 and CCALI_01838, Fig. S7) active on curdlan and laminaran, the storage polysaccharide of brown algae, as recently observed^39^, and a β-galactosidase (NIES2109_37100).

The GH117 family contains almost exclusively α-L-anhydrogalactosidases and one β-D-galactofuranosidase^26^. We detected four α-L-anhydrogalactosidase activities (Fig. S8B, S8C), three belonging to the clade that encompasses all previously characterized GH117 α-L-anhydrogalactosidases (BTO17_12280, BTO17_12335, and Ga0070226_10985), while one appears in a much more distant clade (BTO17_12250; Fig. S5). Noteworthy, three of these enzymes appears within the same genomic region of *Polaribacter reichenbachii KCTC 23969* (Fig. S8A). Interestingly, we also detected three α-L-fucosidase activities, mainly weak, for enzymes from gut and marine bacteria (BcellWH2_02525, P177DRAFT_02789, LV85DRAFT_03918) appearing in a common and previously uncharacterized clade. The two marine enzymes each appears in a PUL with a member of GH95 family, with mostly α-L-fucosidase reported activities. This suggests the polyspecificity in a previously thought highly specific family, other clades remaining to be characterized, such as the group including Ga0062136_116105, Ga0062136_116109, AXE80_07375 and Ga0104390_100627 (from both gut and marine bacteria as well; Fig. S5).

The GH29 family is very large and mostly includes demonstrated α-L-fucosidases. Our screening experiments confirmed that the main product of these enzymes is α-L-fucose, an activity we obtained for 17 members over 31 soluble targets (74%). This is in line with previous functional exploration of the GH29 family which highlighted that the diversity inside the GH29 family was correlated to the diversity of fucosylated substrates and of α-L-fucose linkages^40,41^. However, in a narrow region of the phylogenetic tree grouping both gut and marine bacteria (Fig. S6), we discovered that two members were able to release α-L-galactose from the non-reducing end of desulfated oligo-porphyrans (Ga0070226_10977 and Ga0070226_10982). A similar observation was firstly reported by Robb and co-workers^20^. The poly-specificity in GH29, as in GH117, could be a consequence of the structural similarity of α-L-galactose (or its 6-deoxy form for GH117) with α-L-fucose. To deepen our studies of the GH29 family, seven additional members were selected based on their important sequence diversity despite being all colocalized within only two PULs (Fig. S1, Table S3). All were soluble and active on pNP-α-L-fucose.

### 3. New functions and new CAZy families

Screening experiments revealed enzyme activities for 75 CAZymes spread in 30 CAZy families beyond the aforementioned 39 members of GH29, GH50 and GH117. For most of them, the observed activities were already known in the family and some were also confirmed by other groups during the course of our work (Table S2). However, as for the GH29, GH50 and GH117 families, we also detected novel specificities in some other families.

An enzyme from the GH16_17 subfamily (*B. ovatus*, GH16_17A, HMPREF1069_02099) had a κ-carrageenase activity as expected, but also showed a strong ι-carrageenase activity which was never observed in the GH16 family so far (Fig. 2B, see section 6.2). This completes the list of sulfated galactanases (e.g., porphyranases, α- and κ-carrageenanases) characterized in the GH16 family, illustrating again the evolution and plasticity of the active site of these enzymes for the degradation of agars and carrageenans.

**Figure 2:**
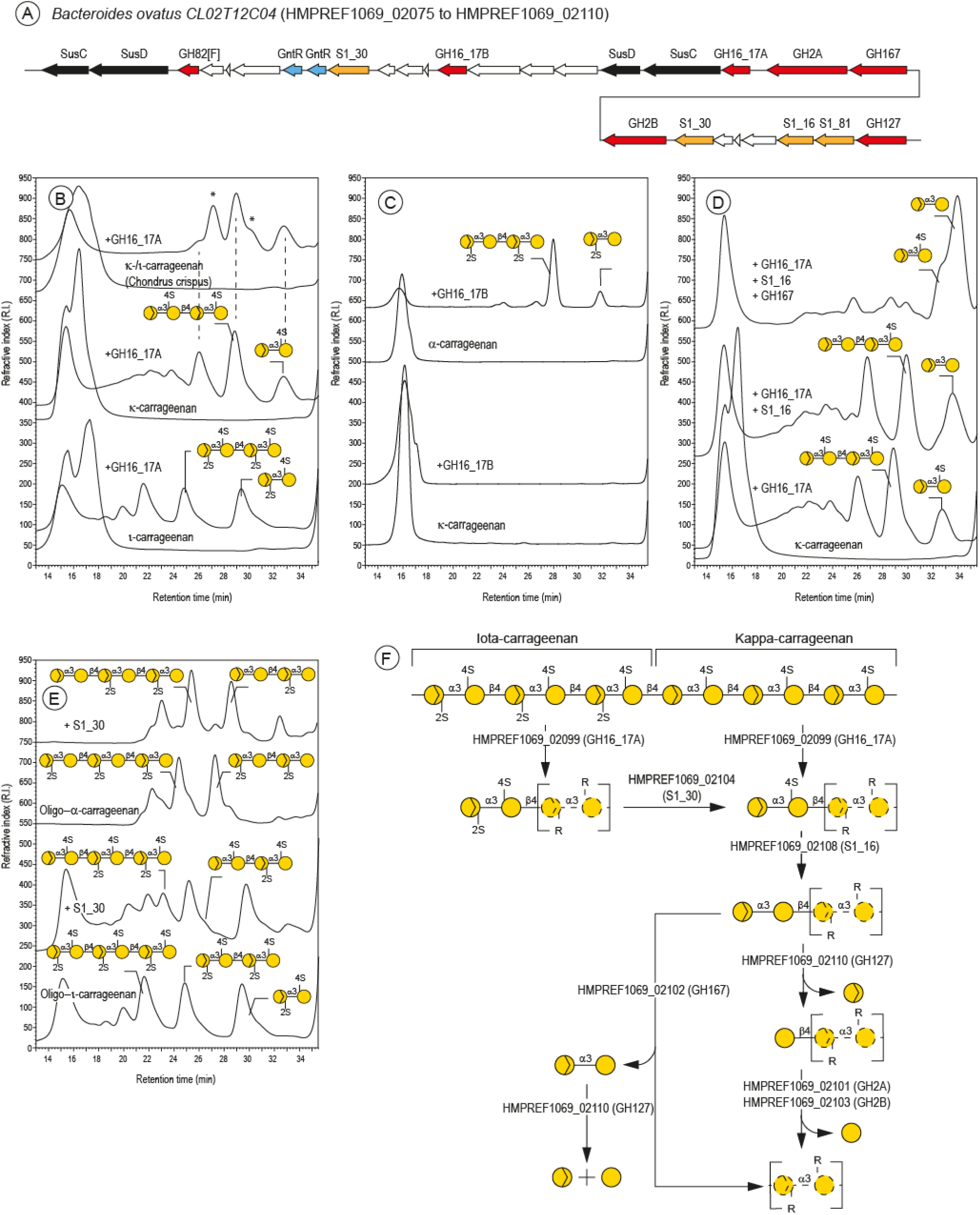
PUL Carrageenan of *Bacteroides ovatus* CL02T12C04. **A**) Organization of the PUL. **B**) Degradation profiles of ι-and κ-carrageenan digested with the GH16_17A κ-/ι-carrageenase (HMPREF1069_02099). Degradation of the hybrid κ-/ι-carrageenan extracted from the red algae *Chondrus crispus* revealed the preference of the carrageenase for the κ-carrageenan fraction. * indicates hybrid oligo- κ-/ι-carrageenans. **C**) Degradation profile of the α-carrageenan degraded by the GH16_17B α-carrageenase (HMPREF1069_02109). **D**) Series of chromatograms obtained after the successive action of enzymes involved in the degradation pathway of the κ-carrageenan. The S1_16 sulfatase (HMPREF1069_020108) removes the sulfate at position 4 of the non-reducing galactose residue. **E**) Incubation of the oligo-ι- and oligo-α-carrageenans with the S1_30 sulfatase (HMPREF1069_02086) demonstrated the desulfation at the position 2 of the galactose residue at the non-reducing end similarly to sulfatase of the S1_81 sub-family. **F**) Deduced degradation pathway of hybrid ι-/κ-carrageenan.

We observed the partial degradation of saccorhizan, a sulfated (xylo)fucoglucuronomannan extracted from the brown algae *Saccorhiza polyschides*, by a polysaccharide lyase of family PL40 (CytfeDRAFT_1478). The PL40 family previously contained a unique characterized enzyme, an ulvan lyase lacking investigations concerning the specificity towards glucuronic or iduronic acids, or both^22^. Ulvan and saccorhizan share D-glucuronic acid residues, which are β-linked to rhamnose and mannose residues, respectively (Fig. S9). We also detected activity on saccorhizan for an enzyme (P177DRAFT_02788) from the PL43 family, which was founded concomitantly with our work based on the recognition and cleavage of D-glucuronic acid in similar sulfated fucoglucuronomannans from another brown algae *Kjellmaniella crassifolia*^42^.

Activity screening of proteins of unknown function unveiled new glycoside hydrolases and polysaccharides lyases, leading to the creation of four novel CAZy families (Table 1; Supplementary 1), while two others were discovered concomitantly, GH167^43^ and PL43^42^. Notably, we characterized by chromatography and NMR, the founding member of the GH_New1 family as a new β-agarase (Ga0070226_10976) located in the porphyran PUL of *C. baltica* (Fig. 2; section 6.1). This family shows remote similarities to GH140 family, and predicted structures of GH_New1 well aligned with GH140 structures and to a structure (PDB entry 4N0R) not associated with any publication and likely part of a distinct but distantly related family. Two additional members of GH_New1 and three homologs of 4N0R (Table S3) were selected and were produced soluble. While none have shown a detectable activity, we identified the putative catalytic amino-acids which are conserved across in these three distantly related families as in the whole GH-A clan (Additional information File 1).

We created two novel PL families, PL_New1 and PL_New2, after the characterization of two alginate lyases (Niaso_1999 and Niaso_2009) in a PUL of *Niabella soli* DSM 19437 (Fig. S10, Table 1). Interestingly, during family creation we noticed bi-modular proteins with a fusion of both PL_New1 and PL_New2 modules (Additional information File 1). Because we did not detect activity for the enzyme Niaso_2005 grouped in PL_New1 family, we investigated five additional PL_New1 members (Table S2) and observed an alginate lyase activity for one of the four soluble enzymes (OPIT5_01565). In this PUL, we characterized several other enzymes involved in alginate breakdown, as well as unrelated activities, suggesting it could target interlinked and complex algal cell-wall polysaccharides.

We identified a new glycoside hydrolase family, GH_New2, built from an α-D-galactosidase (zobellia_334) remotely related to GH36 and GH27 families that both mainly encompass α-D-galactosidases. Four additional members were investigated and three showed the same activity (Bovatus_03161, STSP1_01769 and B2K_10610). We hereafter present the in-depth structural and functional characterization performed on the latter.

### 4. Molecular determinants of the B2K_10610 α-galactosidase founder of GH_New2 family

Enzymology parameters of the B2K_10610 α-galactosidase were determined with *p*NP-α-Gal. The enzyme displayed optimal activity at pH 6.0 at 50 °C, with a K_M_ of 7.56 mM and a catalytic efficiency of 2.37 mM.s⁻¹ (Additional information file 2). We solved the apo crystal structures of B2K_10610 at 2.05 Å resolution, and in complex with a galactose in the active site at 1.50 Å (Figure 4). The enzyme B2K_10610 comprises three folded domains: an N-terminal β-sandwich domain, a (β/α)_8_ central domain (TIM barrel), and a C-terminal β-sandwich domain (Figure 4B). The N-terminal β-sandwich is a large domain, comprising 281 amino acids that form 18 β-strands. The TIM barrel domain is connected to the N-terminal domain by an extended α-helix. The C-terminal domain forms a small β-sandwich composed of 91 amino acids arranged into eight antiparallel β-strands. This overall structure should be conserved within the GH_New2 family, based on the high sequence similarity among its members. This structure revealed a catalytic “pocket” with the galactose bound in the -1 subsite, confirming the exo-acting mode of action of this enzyme. This high-resolution structure enabled the identification of putative catalytic residues, which were subsequently validated by single-point mutagenesis (Additional information File 2). Comparative structural analysis with the closely related GH27 and GH36 families highlighted the specificities of this family regarding the overall structure as well as the catalytic site. Collectively, these findings support the integration of GH_New2 within the GH-D clan. In order to validate the α-galactosidase activity of B2K_10610 on natural substrates, the enzyme was incubated with various oligosaccharide harbouring α-galactopyranose (Gal*p*): 6-α-D-galactopyranosyl-D-glucopyranose (melibiose) and 3-α-D-galactopyranosyl-D-galactoyranose (Gal-α1.3-Gal). Both disaccharides were hydrolysed by the enzyme (Figure 4D, Additional information File 2), confirming its ability to accommodate a terminal galactose in its active site.

### 5. Characterization of new formyl-glycine dependent sulfatases

Carbohydrate sulfatases belong to the formyl-glycine dependent sulfatases family S1 in the SulfAtlas classification^44^ and require a post-translational maturation of the catalytic cysteine or serine to formyl-glycine to obtain active enzymes. Although sulfatases can be well overexpressed in *E. coli*, their maturation is not systematic. Further, carbohydrate sulfatases are often exo-acting enzymes, active on mono- or oligosaccharides, while only few examples of marine endo-acting sulfatase have been observed^45,46,47^. In this context, the selected sulfatases were tested on the oligosaccharide series obtained with the CAZymes co-localized in the same PUL and, when unsuccessful, on unspecific pNP-sulfate.

We determined the substrate specificities for 7 sulfatases (Table 2), with four being colocalized in a PUL dedicated to carrageenan breakdown (Fig. 3; discussed in section 6.2). Interestingly, all the sulfatases for which activity was determined were encoded by species of the *Bacteroidales* order isolated from gut microbiota (16 on 17 selected *Bacteroidales* targets were solubles). Conversely, no activity was observed for the 12 marine sulfatases expressed as soluble protein of 34 selected targets. This suggests that the sulfatase production and maturation by *E. coli* is more efficient, so far, for enzymes originating from the same environment, the human gut, than from the marine environment.

**Figure 3:**
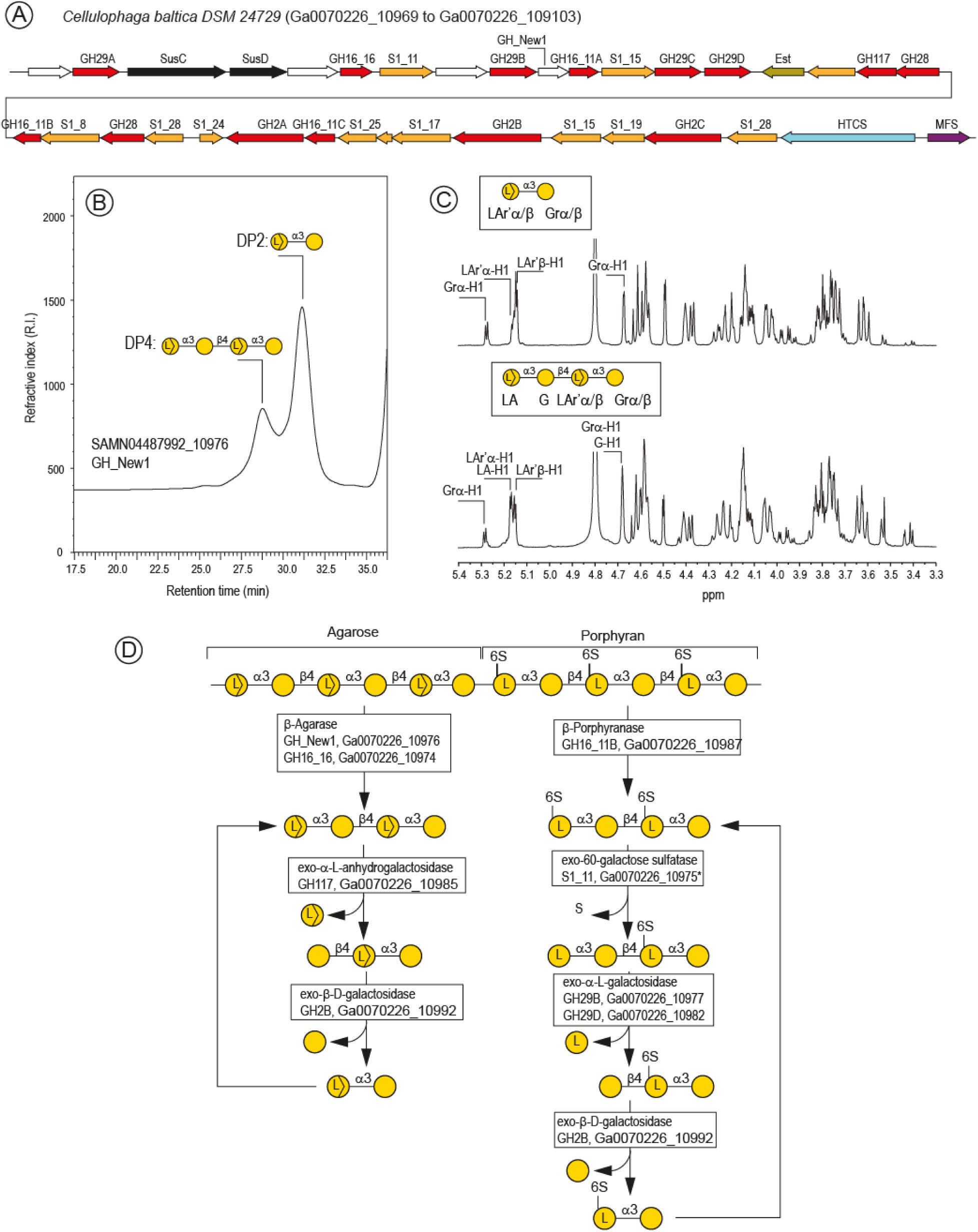
PUL dedicated to the degradation of agars (e.g.; agarose and porphyran). **A**) Organization of the *Cellulophaga baltica* DSM 24729 agars PUL. **B**) Size exclusion chromatogram recorded on the end-products obtained after incubation of agarose with the new GH_New1 agarase (SAMN04487992_10976). **C**) The ^1^H NMR recorded shows that the end-product belong to the neo-oligo-agarose series and demonstrated the cleavage of the β-links. **D**) The degradation pathways of the agarose and the porphyran fractions of the pophyran determined with soluble and active enzymes of the PUL.

We confirmed the △4,5-hexuronate-2-O-sulfatase activity of two S1_9 sulfatase (HMPREF1181_01721 and BcellWH2_02527, Figure S11) in agreement with the studies on this subfamily^45,48,49^.

During the course of our work, the first members of the subfamilies S1_46 and S1_16 were concomitantly reported^50,51^. The first characterized S1_46 member (BT1918) showed an N-acetyl-D-glucosamine-3-sulfate activity, a rare motif found in heparin and heparan sulfate^50,51^. While our collection does not include such a rare substrate, we obtained an activity on *p*NP-sulfate for a S1_46 sulfatase (BACDOR_01585). Interestingly, no gene related to polysaccharide utilization can be identified in the neighbourhood of BT1918, while BACDOR_01585 belongs to a CAZyme cluster including a GH20 characterized in this work as a β-D-N-Acetyl-6-sulfo-hexosaminidase, as well as several GH29 likely targeting α-L-fucoses and a GH117. This raises questions about the specificity of S1_46 to 3O-sulfation of GlcNAc or about the co-occurrence of heparans with fucosylated GAGs.

The first characterized S1_16 was an exo-acting 4O-galactose/-galactosamine sulfatase specific to mucins^50,51^. We also observed specificity towards the position 4O-sulfation of galactose located at the non-reducing end of the oligo-κ-carrageenan for a S1_16 sulfatase (HMPREF1069_02108). The encoding-gene is colocalized with three sulfatases that were characterized as well (see section 6.2). We report the first characterizations within subfamily S1_30 with two sulfatases (HMPREF1069_02086 and HMPREF1069_02104) active on the 2O sulfate group of the anhydrogalactose located at the non-reducing end of the oligo-ι- and oligo-α-carrageenans. Classified in subfamily S1_81, the last sulfatase (HMPREF1069_02109) presents a weak activity towards 2O sulfation of anhydrogalactose of oligo-α-carrageenan as previously observed^52^.

### 6. Structural studies of the S1_30 ι-carrageenan-sulfatase from *Bacteroides xylanisolvens*

In order to contribute to the understanding of the determinants governing substrate recognition in subfamily S1_30, for which no structural data have been reported so far, we determined the crystal structures of an inactive mutant of the human gut bacterium *Bacteriodes xylanisolvens* to 1.41 Å and 1.8 Å resolution, respectively, with and without ligand bound (Additional information File 3). The mutant enzyme, hereafter referred to as *Bx*S1_30_S85A, as previously reported S1 crystal structures^53^, exhibits a two-domain architecture: an N-terminal alkaline phosphatase α/β/α-fold domain (residues 21-397), and a smaller C-terminal domain (residues 398-482). A notable finding is that while most S1 sulfatases bind a calcium ion at the sulfate binding (S) site, Bx_S1_30 lacked this cation in both structures; instead, a water molecule occupied the position and performed hydrogen bonding with the active site residues. This suggests relatively low or context-dependent calcium coordination. The crystal structure of the inactive mutant co-crystallized in the presence of hexasaccharides produced by enzymatic degradation of ι-/ν-carrageenan^54^, revealed an electron density in the active site corresponding to a hexa-saccharide (Figure 5; Additional information File 3, Fig 3B and 3C). Surprisingly, detailed inspection revealed that a neo-ι-ν-ι-carrahexaose (ινι-NC6) was bound in the active site (Figure 5), and not a ι-NC6 as expected (see Supporting Information for details). The high-resolution structure of the enzyme complex allowed the identification catalytic residues, with the alanine residue substituting the catalytic serine (S85A) being positioned ∼3.15 Å below the targeted sulfate ester, as well as other residues involved in substrate binding (Additional information File 3: Fig. 2 and Table 1). His225, being positioned ∼2.9 Å from the scissile bond in an orientation which is in agreement with the protonation of the ester oxygen, is a good candidate as the catalytic acid. Substrate interactions are primarily concentrated in subsite 0 that holds a G4S moiety, and subsite +1 in which binding of the D2S6S unit takes place (Figure 5 and Additional information File 3: Fig. 2B, 2C, 2D, 4 and 5). In subsite 0, all amino acid residues interacting with the G4S are conserved to the *Pseudoalteromonas sp.* PS47 endo-4S-ι-carrageenan sulfatase (*Ps*S1_19A)^55^ with the exception of His317 which holds an Asn (292) at this position and His164 for which no enzyme residue is present in the PsS1_19 counterpart. Comparative studies with *Ps*S1_19A and with *Pf*S1_81 (formerly *Pf*S1_NC)^56^, both displaying activity on ι-NC’s, demonstrate that *Bx*S1_30 possesses unique loop regions essential for the recognition of the D2S6S moiety in subsite +1 (Additional information File 3: Fig. 5). One of these two loops furthermore interact with the DA2S at the non-reducing end subsite -1. The C-terminal domain was found to have no direct interactions with the substrate, placing it approximately 6 Å away from the non-reducing end. Surface representations of *Bx*S1_30, *Ps*S1_19A, and *Pf*S1_81_complexes, highlight broader active site clefts of the former two, contrasting with the narrow pocket of the exo-active S1_81 (Additional information File 3: Fig 6). This feature shows that BxS1_30 possesses endo activity. Overall, these studies and the comparative structural analysis with sub-families S1_19 and S1_81, for which crystal structure complexes with ι-NCs have been reported previously, have allowed identifying differences in subsite environments contributing to the understanding of the substrate specificity differences. In particular, specific features of this family, especially with regards to the recognition of ν-NC could be highlighted.

**Figure 4:**
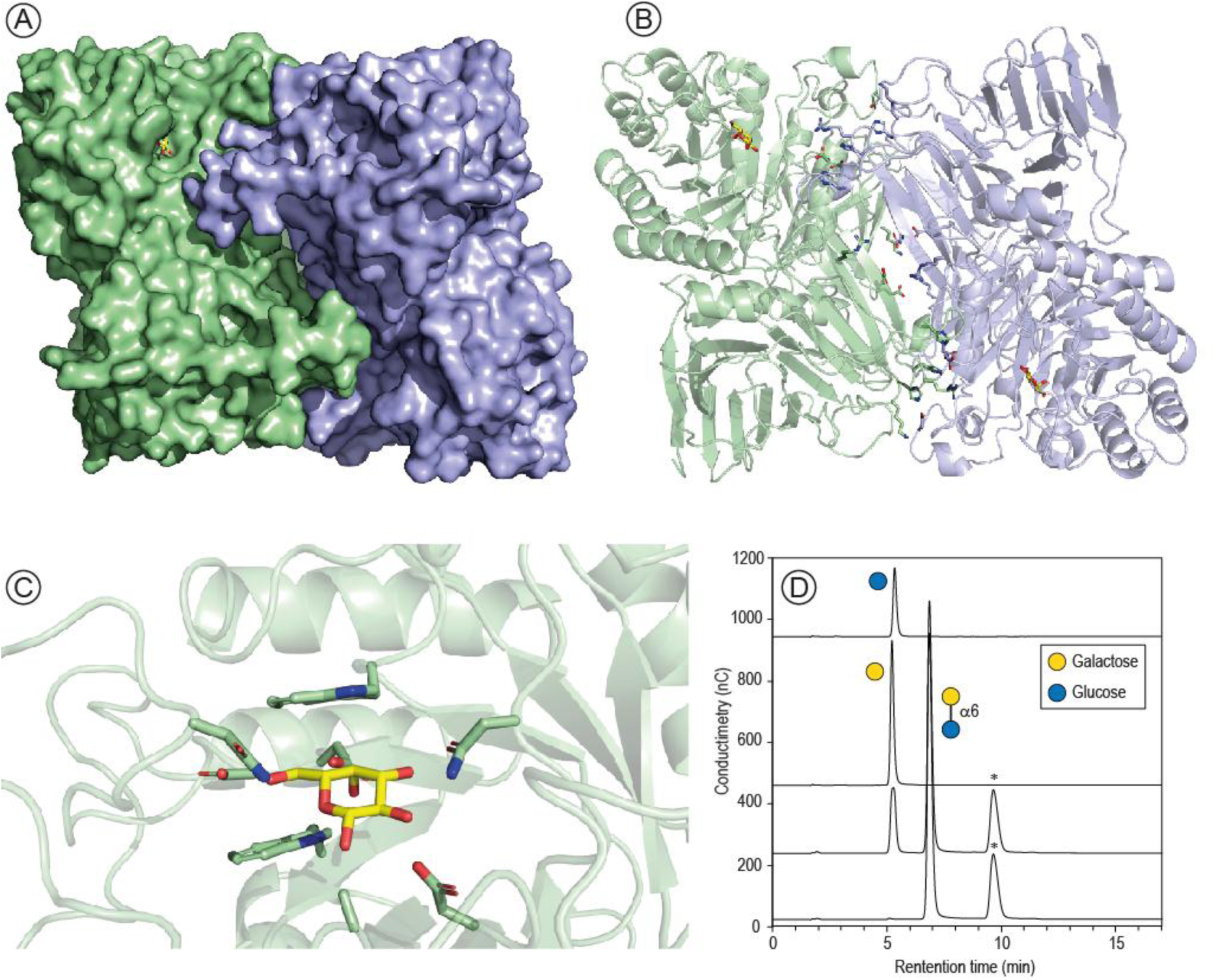
Crystal of the α-galactosidase B2K_10610. **A** and **B**) Overall structure of the enzyme dimer. **B**) Ribbon diagram of the B2K_10610 dimer. Amino acids involved in intermolecular salt bridges shown as sticks. **C**) Active site organization. Amino acids involved in galactose binding are shown in green sticks; the pNP-α-galactopyranoside molecule, bound in the active site, is shown in yellow sticks. **D**) Anion exchange chromatography analysis of the melobiose incubated with the B2K_10610 α-galactosidase. Galactose and glucose served as monosaccharide standards. * Peak 4: buffer

**Figure 5:**
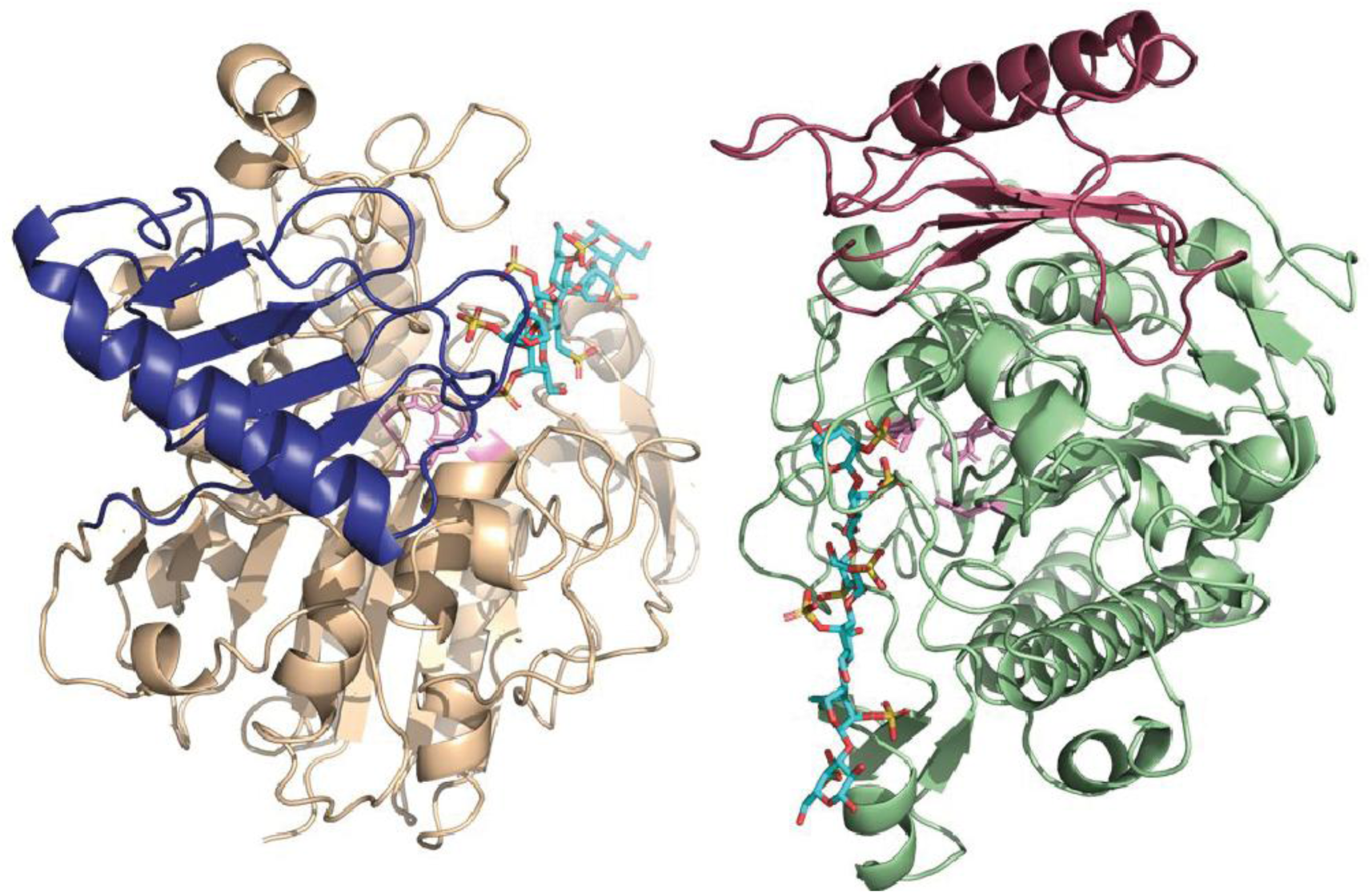
Crystal structure of the *Bacteroides xylanisolvens* ι-carrageenan-sulfatase inactive mutant, *Bx*S1_30_S85A in complex with neo-ι-ν-ι-carrahexaose (ινι-NC6). Ribbon diagram of the overall structure of the enzyme dimer with the N-terminal domains coloured in wheat (monomer A) and green (monomer C), and those of the C-terminal domains coloured in dark-blue (monomer A) and ruby (monomer C), respectively. Amino acids corresponding to the catalytic residues are shown as light-pink sticks. Neo-ι-ν-ι-carrahexaoses are shown in blue sticks.

### 7. PUL synergies in sulfated polysaccharide degradation

Based on our enzyme characterization, we predicted the target substrate of 16 of the selected PULs (Table S4). For several of them, we identified all the catabolic steps leading to the saccharification of the concerned polysaccharides (see below). As expected, we found several PULs targeting sulfated marine polysaccharides extracted from algae (e.g., agars, carrageean, sacchorizan) but also several PULs involved in the degradation of glycosaminoglycan found in marine or land organisms (e.g., chondroitin sulfate, dermatan sulfate, heparan sulfate and heparosan).

#### 7.1. Carrageenan PUL of the human gut *Bacteroides ovatus* CL02T12C04

The α-, ι- and κ-carrageenans are made of the same disaccharide repetition backbone (α-1,3-D-anhydrogalactose linked to β-1,4-D-galactose) but differ by the position of the ester sulfate: position 4 of galactoses (κ-carrageenan), position 2 of anhydogalactoses (α-carrageenan) and both (ι-carrageenan), hereafter denoted 4S_Gal_ and 2S_AnGal_. We identified the ability of a human gut species to synergistically breakdown these polysaccharides (selected PUL 28; Fig. 2A) with two initial endo-acting GH16_17 followed by the coordinated depolymerization by sulfatases and exo-acting GH2, GH127 and GH167.

The prime degradation by endo-acting enzymes allows to cleave β-1,4 linkages and release oligosaccharides despite the sulfate groups. Such enzymes can be found in the GH16_17 subfamily, with 14 κ-carrageenases from marine origin reported so far. The degradation profile of the first GH16_17A in this PUL (HMPREF1069_02099) revealed both κ-carrageenase and ι-carrageenase activities (Fig. 2B; endo-producing di- and tetra-saccharides). The incubation of enzymes with hybrid κ-/ι-carrageenan (extracted from *Chondrus crispus*) lead to the production of neo-oligo-κ-carrageenan and hybrid neo-oligo-κ-/ι-carrageenan series, but no neo-oligo-ι-carrageenan series, suggesting that the κ-carrageenan fraction is the preferential substrate of this enzyme. In contrast, the GH16_17B PUL member (HMPREF1069_02093) was inactive on κ- and ι-carrageenan, but successfully degraded α-carrageenans (Fig. 2C). These two GH16_17, first characterized members from human gut species, can thus accommodate 2S-anhydrogalactose in their active site, an original specificity not reported yet in marine GH16_17, despite the ι-carrageenan binding affinity of the CBM92 in bimodular Cgk16A^58^.

The released neo-oligo-carrageenans display sulfations at their non-reducing ends, preventing cleavage by exo-acting CAZymes. Screening assays revealed four sulfatases in this PUL (see section 5 and Table S2) with activities compatible with the sequential removal of their sulfate groups. We further confirmed by chromatography sulfatase activities of a S1_16 subfamily member (HMPREF1069_02108) targeting 4S_Gal_ in neo-oligo-κ-carrageenans (Fig. 2D) and a S1_30 member (HMPREF1069_02086) targeting 2S_AnGal_ in neo-oligo-α/ι-carrageenans (Fig. 2E).

The complete depolymerization is accomplished by a GH167 (HMPREF1069_02102), producing neo-carrabioses confirmed by chromatography (Fig. 2E), and followed by exo-acting GH2 (GH2A: HMPREF1069_02101; GH2B: HMPREF1069_02103) and GH127 (HMPREF1069_02110) releasing β-1,4-D-galactose and α-1,3-D-anhydrogalactose, respectively (Table S2). The final overall scheme (Fig. 2F) is similar to that previously described in marine organisms^16,52^ while involving different enzyme (sub)families.

#### 7.2. Porphyran utilization by the marine *Cellulophaga baltica* DSM 24729

Porphyran, the matrix cell wall polysaccharide of the red algae *Porphyra sp*., is composed of two disaccharides repetition moieties: agarobiose (neutral) and porphyranobiose (sulfated)^8^. Complete degradation of porphyran orchestrated by the proteins encoded in PUL (selected PUL 40; Fig. 3A) requires, at first, the initial endo-cleavage by β-porphyranases or β-agarases to produce oligosaccharides. These activities, known in subfamilies GH16_11 and GH16_16, were confirmed here (Ga0070226_10987 and Ga0070226_10974, respectively). We also detected β-agarase activity with the founding member of the GH_New1 family (Ga0070226_10976) (Fig. 3B-3C). The depolymerisation of oligo-porphyrans involves a succession of exo-acting enzymes, as previously described^18,20,57^. First a sulfatase is required to remove the 6O-sulfation at the non-reducing end. Over the 12 sulfatases of this PUL, none of the three enzymes expressed as soluble (from subfamilies S1_15, S1_19 and S1_25) were active on oligo-porphyrans nor any other sulfated substrates in our collection. Therefore, we used *B. plebieus* S1_11 (bacple_01701) to prepare desulfated oligo-porphyrans. These could be further degraded by the successive action of demonstrated α-L-galactosidases in families GH29 (GH29B: Ga0070226_10977; GH29D, Ga0070226_10982; Fig. S6) and GH117 (Ga0070226_10985, Fig. S5) followed by β-galactosidase in family GH2 (GH2B: Ga0070226_10992). Aside from the predictive depolymerization scheme (Fig. 3D), the involvement of additional enzymes not produced as soluble according to our experimental conditions: GH29 (GH29A: Ga0070226_10970 and GH29C: Ga0070226_10981), GH2 (GH2C: Ga0070226_10997) and two members of the GH28 family, known to encompass only α-galacturonidase activities so far, probably participate in the porphyran metabolism.

#### 7.3. PULs dedicated to the degradation of glycosaminoglycans

Three PULs in human gut species containing GH29 or GH117 were found to digest glycoaminoglycans: heparan sulfate/heparosan and chondroitin sulfate. Heparan sulfate and heparosan share the same major disaccharide repetition unit, made of β-1,4-D-glucuronic acid linked α-1,4-N-acetyl-D-glucosamine. Heparosan sulfate displays additional sulfations at position 2 of the glucuronic acid and position 6 of the galactosamine residues. In *Bacteroides stercoris* CC31F, a PUL (selected PUL 31; Fig. S11A) revealed six genes encoding for heparan sulfate/heparosan lyases according to screening assays (Table S2). Size-exclusion chromatography monitoring of heparan sulfate incubated with two PL15_2 (BsPL15_2A: HMPREF1181_01722, BsPL15_2: HMPREF1181_01727) showed the unique release of unsaturated disaccharides (Fig. S11B, S11C), similarly to reducing-end exolytic heparinase from the PL15_2 subfamily^59^. The disaccharide end-products were completely degraded by the combined action of a △4,5-hexuronate-2-O-sulfatase from subfamily S1_9 (HMPREF1181_01721, Fig. S11D, S11F) followed by an unsaturated glucuronyl hydrolase from family GH88 (BsGH88: HMPREF1181_01743, Fig. S11C). The removal of the sulfate groups of the unsaturated residues was mandatory to allow the breakdown of this terminal residue by the GH88 enzyme. On heparosan, one PL15_2 (BsPL15B: HMPREF1181_01727) additionally released large amounts of tetrasaccharides while the other PL15_2A (BsPL15_2A: HMPREF1181_01722) and the PL33_1 (BsPL33_1: HMPREF1181_01744) show similar profiles (Fig. S11E) and only the GH88 (BsGH88: HMPREF1181_01743) is required for complete degradation. Heparosan was also catabolized by a PUL of *Bacteroides clarus* YIT 12056 (selected PUL 39, Fig. S11A) following a similar scheme to that deduced for *B. stercoris* CC31F (Fig. S11G), as supported by chromatography of the homologous PL15_2 (BcPL15A: HMPREF9445_03076; BcPL15B: HMPREF9445_03083) and PL33_1 (BcPL33_1: HMPREF9445_03068) (Fig. S11D) and the presence of two GH88 (BcGH88A: HMPREF9445_03069; BcGH88B: HMPREF9445_03080). In addition, these PULs contained α-L-fucosidase from GH29 and GH95 families, some characterized in this study (BsGH95: HMPREF1181_01727; BsGH29: HMPREF1181_01742 and BcGH29: HMPREF9445_03070), while α-L-fucose has never been reported in heparin-like polysaccharides to our knowledge. Such co-localization suggests that the substrate differs from the known human glycosaminoglycan used in this study, such as the fucosylated glycosaminoglycan found in the Japanese scallop, *Patinopecten yessoensisfrom*^60^.

Chondroitin sulfates (CS) are composed of a repeating disaccharide (β-1,3-D-glucuronic acid and β-1,4-N-acetyl-D-galactosamine, GalNAc) with different sulfation patterns in the different fractions: position 4 and 6 of GalNAc in CS-A and CS-C respectively, and at both position 2 of glucuronic acid and position 4 of GalNAc in CS-D. In *Bacteroides cellulosilyticus* WH2, a PUL (selected PUL 16; Fig. S12A) encodes three chondroitin lyases of families PL8_2 (BcellWH2_02521, BcellWH2_02528) and PL30 (BcellWH2_02524; previously characterized on hyaluronan)^26^. Chromatography analyses indicate they specifically release unsaturated disaccharides specific of the three fractions (Fig. S12B-C). The sulfate located at the position 2 of the unsaturated residues of CS-D units are removed by a family S1_9 sulfatase (BcellWH2_02527; Fig. S12D). We were not able to observe further degradation of the disaccharides using other proteins encoded in this PUL, notably proteins of unknown function that could compensate for the absence of a co-localized GH88. Therefore, we used an unsaturated glucuronyl hydrolase (GH88; NCTC13071_00479) of *Prevotella oris* which successfully removed the unsaturated residues in desulfated CS-B (Fig. S12D) and CS-C (Fig. S12E) units. This renders the sulfate groups on GalNAc accessible to the S1_15 sulfatase (BcellWH2_02518; Fig. S12E). Although this PUL encodes several enzymes involved in the degradation of chondroitin, it lacks an unsaturated glucuronyl hydrolase and a sulfatase acting on the position 4 of GalNAc. Interestingly, two genes encoding enzymes from GH88 and S1_27 families, whose members have been reported with the desired specificity, could be found about 10 genes upstream this PUL (BcellWH2_02505 and BcellWH2_02506).

## Discussion

In this work, we characterized enzymes from a set of selected PULs and detailed their concerted activity in the degradation of algal polysaccharides as well as of glycosaminoglycans occurring in land and marine organisms. The selection of the targets followed a new strategy.

Genes encoding members of three relevant families – GH29, GH50 and GH117 – located in PUL were extracted to compute phylogenetic trees labelled with PUL compositions guiding the exploration of diversity. We pointed out the poly-specificity of the three families used as starting point, in agreement with the diversity of the PULs listed in the respective families. Thanks to the large diversity of substrates used in screening assays, we were able to discover novel specificities (e.g., saccorhizan lyase) and to demonstrate the activity of more than 130 enzymes grouped in 29 CAZy families, in five sulfatase S1_subfamilies and in four newly established CAZy families (GH_New1, GH_New2, PL_New1, and PL_New2). However, substrate availability remains a major limitation to identify the natural substrate of PULs and cognate enzymes, for instance fucosylated heparan sulfate-like or branched agars-like.

We noticed that the protein expression of marine enzymes in *E. coli* was lower than enzymes of the gut microbiota. Also, the sulfatase maturation was better for enzymes of gut species. All these observations suggest that using *E. coli* is an excellent expression system for gut enzymes likely due to their shared environmental origin. With only about 50% of marine proteins well expressed in *E. coli*, the characterization of marine enzymes and the synergy of their PULs are progressing at a slower pace, calling for the development of marine bacterium expression platforms. In the frame of a medium throughput screening strategy carried out in this study, the expression conditions were not optimized systematically for each target while some enzymes could be obtained soluble using more elaborate experimental conditions. Nevertheless, two expression temperatures (18°C and 25°C) were tested but didn’t significantly improve the amounts of soluble proteins.

Another bottleneck for the characterization of marine PULs is the production of active sulfatases. Indeed, while the initial production of sulfated oligosaccharides by endo-acting enzymes is not usually impaired by the sulfate groups, they frequently prevent the complete depolymerization by exo-acting enzymes. Thereby, while we produced a series of marine oligosaccharides (e.g., porphyrans, fucans, carrageenans and saccorhizan), only few marine sulfatases were produced as soluble (7 of 34) and none were active, although they belong to subfamilies known to encompass carbohydrate sulfatases. This was notably a strong limitation in the case of saccorhizan (extracted from the cell wall of brown algae *Saccorhiza polyschides*; Fig. S9), the oligosaccharides generated by our novel saccorhizan lyase could not be desulfated by any tested sulfatase. This prevented us from confirming on a natural substrate the α-L-fucosidase and β-D-xylosidase activities obtained on *p*NP-substrates, while in agreement with the polysaccharide composition.

Beside marine organisms, we selected some PULs from gut bacteria given that previous studies have demonstrated the ability of our flora to degrade marine polysaccharides^18,23,25^. We discovered a carrageenan PUL in *B. ovatus* CL02T12C04 which differs from the characterized degradation systems found in marine organisms as it contains the first GH16 able to cleave ι-carrageenan and three carrageenan sulfatases including a new specificity for a S1_16 subfamily and the first characterized members of the S1_30 subfamily subfamily along with the first reported crystal structures from this latter subfamily. The discovery of this carrageenan PUL suggests that mining the enzymatic diversity should not be limited to marine bacteria as the animal gut also offers opportunities to discover new enzymes and degradation pathways for marine polysaccharides.

Because of the stereochemical complexity of carbohydrate and, more especially, polysaccharides, estimation of the diversity of marine polysaccharide biomass is facing to the nowadays technical difficulties to isolate and to sequence polysaccharides as it is envisioned, for example, in large sequencing programs of marine genomes and metagenomes. However, based on the genomic and metagenomic databases, it is possible to list the diversity of the polysaccharide degrading systems without knowing the diversity of the marine polysaccharides recycled in the ocean. The pioneering work of Lapébie and co-workers^35^, who calculated that the number of enzyme combinations found in PULs (several thousands) are far lower than the possible combination to build polysaccharides^61^, suggests that assessing of the diversity of recycled polysaccharides is challenging but attainable by combining innovative bioinformatic strategies and rational selection of enzyme targets for screening methods to improve. In this context, although the work presented herein highlighted some limitations concerning the medium throughput heterologous expression of marine enzymes and the need of an as large as possible set substrates, the discovery of new marine CAZy families and the attribution of new functions to enzymes paves the way to a comprehensive description of the diversity and recycling process of the marine polysaccharide biomass.

## Material and Methods

### Target selection

Families GH29, GH50 and GH117 protein sequences in a predicted Bacteroidota PUL were extracted from the PULDB database (June 2018)^29^. Multiple sequence alignments were generated using MAFFT (version 7)^62^ using L-INS-i parametrization for local alignment optimization, and manually analysed to discard fragmented sequences. Phylogenetic trees were reconstructed using the PhyML (version 3) software^63^ using default parameters. For each protein, the composition of their PUL was extracted from PULDB, transformed into a non-redundant list of CAZyme families and sulfatase subfamilies and indicated at leaf labels in the tree displayed in iTOL (version 3)^64^. For each of the 41 selected PULs (Fig. S1-S3), member proteins were investigated for their domain composition with Pfam^65^ version 32 and their predicted structure using Phyre2^66^, to manually identify proteins of unknown functions that could be novel CAZymes. These genes as well as relevant CAZymes and sulfatases were ordered as synthetic genes.

### Cloning and overexpression assays

The production of the 454 targeted genes in expression vector were outsourced to the Twist Bioscience company (https://www.twistbioscience.com). Genes were codon optimized, synthesized and cloned in pET29b (kanamycine resistant) using Nde1 and Xho1 restriction sites. The genes encoded His-tag at the N-terminal of the CAZymes and the unknown enzymes, but at C-terminal for the sulfatases. The expression vectors were transformed in *E. coli* BL21(DE3) pLysS strain (chloramphenicol resistant).

Transformed *E. coli* strains with the plasmid of interest were grown overnight at 37°C in 2mLLuria-Bertani broth completed with kanamycine and chloramphenicol antibiotics. 50mL NZY auto-induction LB media (NZYtech) was inoculated with 500µL of preculture and maintained at 37°C for 4 hours prior to cooling down the culture to 25°C for 10 hours. Bacterial cells were pelleted by centrifugation and resuspended in 4mL binding medium (10 mM imidazole, 300 mM NaCl, 50 mM Tris-HCl, pH 8) and 40µL of lysozyme (50 µg/mL). After a freeze-thaw cycle, 40µL of DNAse solution (0.67 mg/mL bovine DNAse I, 1.33M MgSO_4_) was added to decrease the viscosity of the samples (∼20 min, 10°C). The cell debris were removed using an Allegra 64R Benchtop centrifuge (Beckman Coulter) working at 20000g for 30min at 4°C. The supernatant was transferred with 400mL microbeads Protino Ni-NTA Agarose (Macherey Nagel) in disposable Reveleris™ Empty Solid Loader Tube/Frit (Buchi) mounted on a Visiprep™ SPE Vacuum Manifold (Supelco). After washing three times with 700µL of binding buffer, the proteins were then collected by adding twice 800µl of elution buffer (500mM imidazol, 300 mM NaCl, 50mM Tris-HCl, pH 8). The solubility and the molecular weight were assessed by gel electrophoresis.

### Screening experiments

Screening experiments were conducted according to the strategy previously reported^26,67^. The proteins were incubated with the polysaccharide substrates in 96-well microtiter filter plates (10 kDa, PES, Pall corporation) overnight at 25°C. After filtration on a multiscreen HTS vacuum manifold (MSVMHTS00, Millipore) connected to a high-output vacuum pressure pump (Millipore), the concentration of the reducing ends in the filtrate was measured by the ferricyanide method^68^.

Positive hits were validated by analysing the enzymatic degradation products by gel permeation chromatography using Superdex S200 10/300 and Superdex peptide 10/300 (GE Healthcare) columns mounted in series and connected to a high-performance liquid chromatography (HPLC) Ultimate 3000 system (Thermo Fisher). The injection volume was 20 µL and the elution was performed at 0.4 mL.min^-1^ in 0.1 M NaCl. Oligosaccharides were detected by differential refractometry (Iota 2 differential refractive index detector, Precision Instruments). Complex sulfated oligo-saccharide mixtures (e.g., glycoaminoglycan oligosaccharides) were analysed by high performance anion exchange chromatography (HPAEC). The sample were injected on a IonPac™ AS11 column (Dionex™) mounted on Dionex ICS 3000 chromatography system equipped with a conductimetry detector. Elution was performed at a flow rate of 1mL.min^-1^ with linear gradient of NaOH, starting from 3% to 100% 0.29M.

Enzyme activities were also confirmed with enzymes highly purified by affinity chromatography using a 1 mL HisTrap™ HP column (GE Healthcare) connected to a NGC chromatography system (Bio-Rad). The fractions containing the protein of interest were pooled and injected on a gel permeation ENRich650 column (Bio-Rad) and eluted in 20 mM Tris-HCl pH 7.5, 100 mM NaCl. The purity of the fractions was assessed using 10% SDS-PAGE analysis. To prevent possible errors, the plasmids harbouring the encoding genes of interest were sequenced again.

To minimize consumption of substrate during our screening procedure, the substrates were divided in three sub-collections composed respectively of substrates containing α- or β-anomeric linkages (α- and β-banks) and poly-uronic acid substrates (lyase bank). At first, the proteins were tested on the sub-collection based on relatedness to a known family since the orientation of the glycosidic bond is strictly conserved within a family^69,70^. When no activity was found, all the substrates available were then tested in a second round.

### Characterization of enzyme end-products

End-products obtained after of oligo- or polysaccharide substrate incubated with glycoside hydrolases, polysaccharide lyases and sulfatases were purified on a semi-preparative size-exclusion chromatography system which consisted of a Knauer pump (pump model 100), a HW40 Toyopearl column (120 x 16 mm; Tosoh Corporation), a refractive index detector (Iota 2, Precision Instruments) and a fraction collector (Foxy R1) mounted in series. Elution was conducted at a flow rate of 0.4 mL/min at room temperature using 100 mM (NH_4_)_2_CO_3_ as the eluent. The fractions containing pure oligosaccharides were collected and freeze-dried.

Carbon-13 and proton NMR spectra were recorded with a Bruker Avance 400 spectrometer operating at a frequency of 100.618 MHz for ^13^C and 400.13 MHz for ^1^H. Samples were dissolved in D_2_O at a temperature of 293 K for the oligosaccharides and 343 K or 353 K for the polysaccharides. Residual signal of the solvent was used as the internal standard: HOD at 4.85 ppm at 293 K and 4.35 ppm at 343 K. ^13^C spectra were recorded using 90° pulses, 20,000 Hz spectral width, 65,536 data points, 1.638 s acquisition time, 1 s relaxation delay, and between 8192 and 16,834 scans. Proton spectra were recorded with a 4006 Hz spectral width, 32,768 data points, 4.089 s acquisition times, 0.1 s relaxation delays, and 16 scans.

The ^1^H and ^13^C-NMR assignments were based on ^1^H-^1^H homonuclear and ^1^H-^13^C heteronuclear correlation experiments (correlation spectroscopy, COSY; heteronuclear multiple-bond correlation, HMBC; heteronuclear single quantum correlation, HSQC). They were performed with a 4006 Hz spectral width, 2048 data points, 0.255 s acquisition time, 1 s relaxation delay; 32 to 512 scans were accumulated.

