## Additional File 1 for "Screening for Polysaccharide Utilization Loci Targeting Marine Polysaccharides"

### Description of novel CAZyme families

#### GH_New1

This family is mainly found in Bacteroidota and to a lesser extent in Gammaproteobacteria and Planctomycetota. A single copy appears in these genomes, except a handful with two copies and *Polaribacter sp.* Q13 having three, each in a distinct genomic cluster. These proteins are only composed of the catalytic domain, a TIM barrel flanked at N- and C-terminal by β-sheets subdomains (Additional Figure 1A), conserved in all members. These domains are never associated, so far, to any known CAZy nor Interpro domain[1]. This family show remote homology to the GH140 family, which reported α-apiosidases (PDB: 5MSY and 8T9W) do not display an equivalent N-terminal module. Both families are part of the GH-A clan. We could thus identify the catalytic amino-acids, two glutamates (Glu275 and Glu353 in the characterized member; see Additional Figure 1B-C), fully conserved in the family and highly conserved within this clan. In this work, a β-porphyranase activity (EC 3.2.1.178) was demonstrated in the *Cellulophaga baltica* DSM 24729 member. In PUL context (Additional Figure 1D),

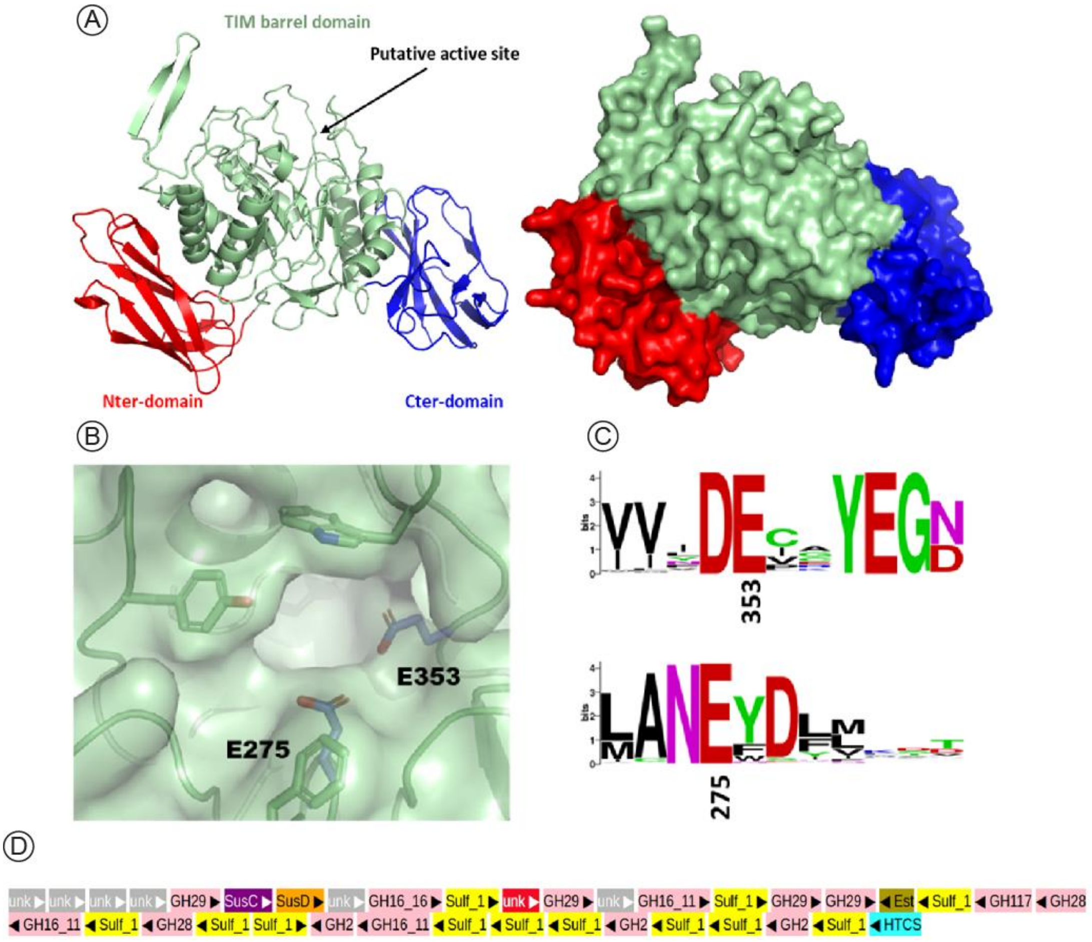


**Additional Figure 1: Analysis of the predicted structure and PUL organization of the characterized member Ga0070226_1097**. **A**) AlphaFold- structure of Ga0070226_10976, revealing a central TIM barrel domain linked to β-sheet-rich domains at both the N- and C-termini. **B**) Identification of two glutamic acids (Glu275 and Glu353) at the surface of the putative catalytic groove. **C**) Logo of sequence conservation around Glu275 and Glu353 among GH_New1 members. **D**) Ga0070226_10976 PUL in PULdb, highlighted in red.

its most frequent partners are GH16_11 and GH16_16, GH29, GH117 and sulfatases confirming porphyran/agar as a preferential substrate.

#### GH_New2

This family is found predominantly in Bacteroidota, and in many other bacterial phyla and a single archeal genome so far. The four characterized members cover a large sequence and taxonomic diversity, with 20% to 35% identity (Blastp) across the two Bacteroidota (a human gut and a marine species), the Bacillota (from soil) and the Planctomycetota (from a saline sediment) proteins. In the majority of genomes, it is found in a single copy (~90%), with some exceptions having an additional copy, such as half of *B. ovatus* strains, for which we characterized one of the copies in the model strain ATCC 8483 (Bovatus_03161). A detailed functional and structural characterization of the Bacillota representative (B2K_10610) is provided in Additional Information File 2. The GH_New2 proteins seem only constituted of this catalytic module, as no additional module could be detected from CAZy nor Interpro. The PUL contexts show a strong association with GH2 and GH3 families, likely driven by many human gut Bacteroides stains. However, given the substrate diversity in these families the prediction of a substrate is difficult (Additional Figure 2). In the marine *Z. galactanivorans*, the PUL possesses only GH3 but much more putative fucosidases (two active on pNP in this work) which, along with the presence of sulfatases, suggests a marine fucan substrate.

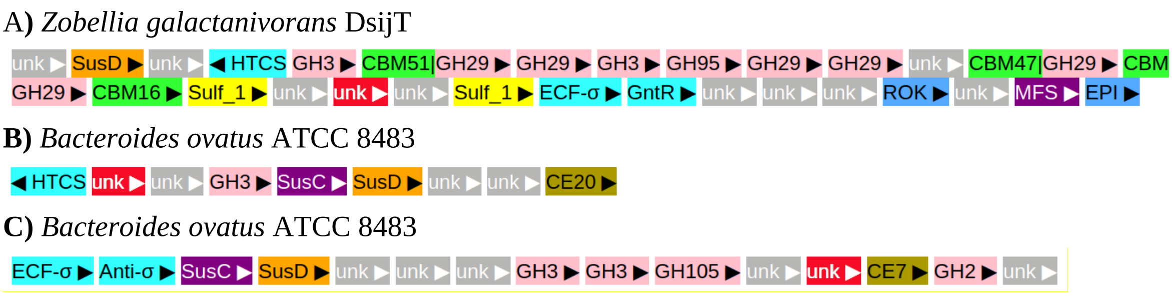


**Additional Figure 2: PUL organizations** for two characterized members (**A-B**) and another copy (**C**), non-characterized in the same genomes as (**B**), all in red. In *Z. galactanivorans* (A) ZOBELLIA_334 co-occurs with GH29, GH95 and sulfatases suggesting a sulfated fucan substrate. Among the rare organisms with two copies, the model *B. ovatus* ATCC 8483 displays: (B) Bovatus_03161 in a small PUL with only an extra GH3 and carbohydrate esterase, and (C) Bovatus_02938 in a larger PUL with both GH2 and GH3 (GH_New2's most frequent partners), a GH105 (despite the absence of PL in the neighborhood), and an esterase as well, for [illegible]

#### PL_New1

This family, found predominantly in Bacteroidota genomes, and spread across many bacterial groups (Proteobacteria, PVC, Terrabacteria), is also retrieved in many fungal genomes (Ascomycota, Basidiomycota and Mucoromycota). The three characterized members cover a large sequence and taxonomic diversity, with 26% to 35% identity (Blastp) between the two Bacteroidota (a human gut and a soil species) and the Verrucomicrobiota sequences. They display a β-helix fold, as other PLs, according to AlphaFold structures (Additional Figure 3). However, sequence and structural alignments, across family members or with related families (e.g., PL29) did not allow the identification of highly conserved surface amino-acids as putative catalytic residues. In complete genomes, one copy is usually

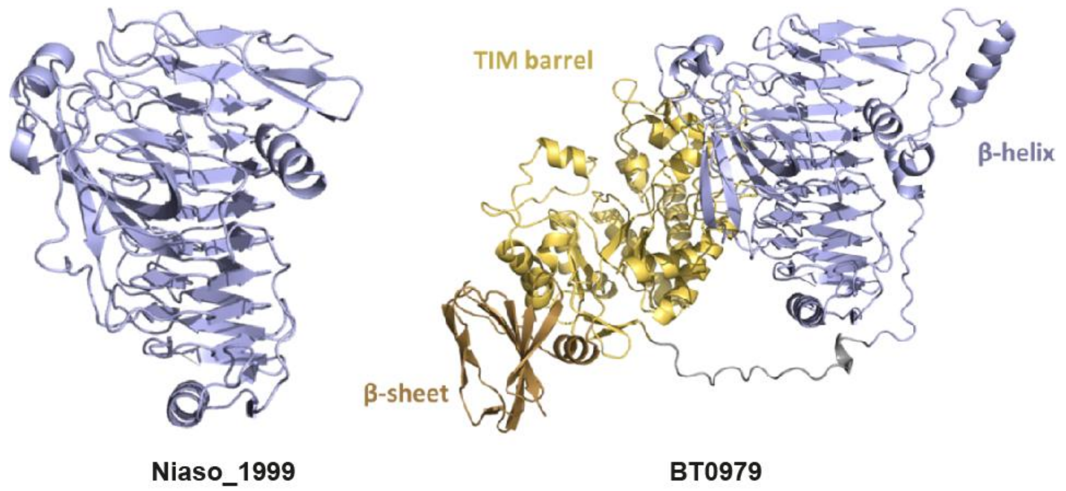


**Additional Figure 3: AlphaFold models** of the monomodular Niaso_1999 and multimodular BT0979 with the additional domains. The β-helix domain is shown in blue, while the C-terminal TIM barrel and β-sheet domains of BT0979 are represented in gold and maroon, respectively.

observed (>70%), or two (~14%), while some species developed more extensive repertoires, with more than 6 copies in *Opitutaceae bacterium* TAV5 (one characterized here), *Paenibacillus sp.* H1-7, *Basidiobolus meristosporus* CBS 931.73 and many fungal species among the Mortierellaceae family. A significant proportion of members (~35%) from bacteria but not restricted to a single phylum, display an additional C-terminal modules of unknown function, a TIM barrel linked to a β-sheet modules, leading to proteins twice longer (~1 000 aa-long / ~110 kDa), such as BT0979 (Genbank ID: AAO76086.1) functionally characterized in this work. Homology searches based on the primary sequence of the TIM barrel showed it only exists attached to PL_New1 and, while its fold is similar to GH-A clan, the similarity levels are too low to guess if it could be a CAZyme, another enzyme or else. The domain architectures of PL_New1-containing proteins mostly consist of the sole catalytic module (~53%, such as Niaso_1999 and OPIT5_01565 – Genbank IDs AHF15512.1 and AHF89142.1 –, characterized in this work; Additional Figure 3). Some homologs were found appended with CBMs (8% of members), notably CBM32 in distinct bacterial phyla (mostly in Paenibacillus) and rarer

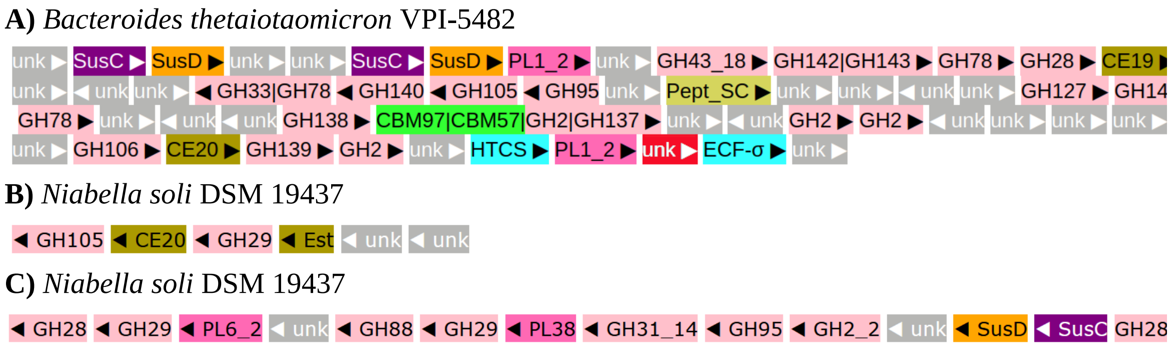


**Additional Figure 4: PUL organizations** in the two Bacteroidota characterized members. In *B. thetaiotaomicron*, BT0979 (in red) appears with CAZyme partners targeting RG-II (**A**) while in *N. soli*, Niaso_1999 (not shown) was not included in a PUL prediction as on the opposite strand but directly upstream the clusters of Niaso_2000-Niaso_2005 (**B**) and of Niaso_2006-Niaso_2019 (**C**) which could collaborate in the degradation of a complex substrate given the specificity of the neighboring CAZymes.

associations found with PL_New2 (~1%, mostly in Bacteroides) one shown to display alginate lyase activity in this work; and even some GHs (~1%), notably GH88 in Sphingobacterium. The family GH88 are known to act on the products of lyase activity, even if only reported on unsaturated glucuronyl residues so far. All three characterized PL_New1 members display the same activity, alginate lyase (no generic EC number exists) while we noticed remote similarity to the PL29 family, which has a single characterized member to date, active on several chondroitin types. In PUL contexts, we observe many members in rhamnogalacturonan-II (RG-II) PULs, including one characterized here, in the model RG-II PUL of *Bacteroides thetaiotaomicron*[2] (Additional Figure 4). Interestingly, another characterized member in *Niabella soli* belongs to a PUL which likely target another substrate as beyond several alginate lyases, reported activities of GHs suggest the presence of fucose, α-galactose, α-galacturonic acid and β-glucuronic acid residues. All these call for further investigation of the functional diversity of this family.

#### PL_New2

This family is predominantly found in Actinomycetota and Bacteroidota phyla, and to a lesser extent in some marine Gammaproteobacteria. The three characterized members cover a large sequence and taxonomic diversity, with 55% identity (Blastp) between the two Bacteroidota (a soil and a human gut species) sequences, having only 29-34% identity to the marine Gammaproteobacteria one. The predicted AlphaFold structure of Niaso_2009 adopts a β-jelly roll fold, characterized by an extended β-sandwich architecture (Additional Figure 5). This model reveals a putative catalytic groove consistent with an endo-acting mode of action of this enzyme. Additionally, positively charged residues line the surface of this groove could promote electrostatic interactions with negatively charged alginate polymers. A tyrosine residue, conserved in this family (Tyr237 in Niaso_2009), is located at the center of the groove. Given its location, orientation and conservation across this newly identified family, this residue could play a key catalytic role, potentially acting as both a Brønsted base and acid, such as observed in alginate lyase family PL7[3].

In most genomes, it is found as a single copy (~71%), with the remaining having an additional copy and only a handful with a third copy (*Aquimarina sp.* BL5, *Nonomuraea sp.* CA-141351, *Paenibacillus sp.* H1-7). PL_New2 proteins are mostly constituted of this sole catalytic module (89%), with few containing various CBMs (8%; mainly CBM32 and CBM35, and few combined with PL_New1 (3%). The PUL contexts show a strong association with PL6, PL17 and PL38 families, likely mainly driven

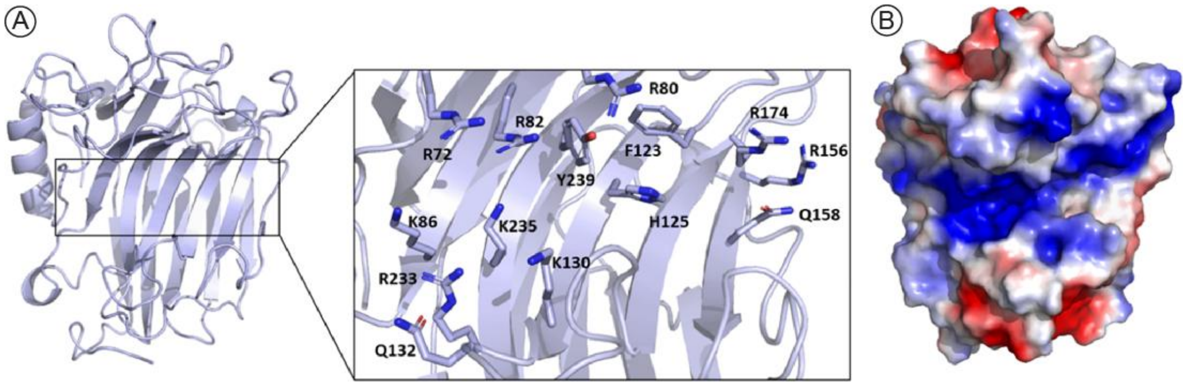


**Additional Figure 5 - AlphaFold models** of the characterized monomodular Niaso_2009. A) Overall structure of Niaso_2009 and zoom on the putative active site with candidate amino-acids for alginate coordination represented as sticks. B) Electrostatic surface highlighting the positive-charged groove.

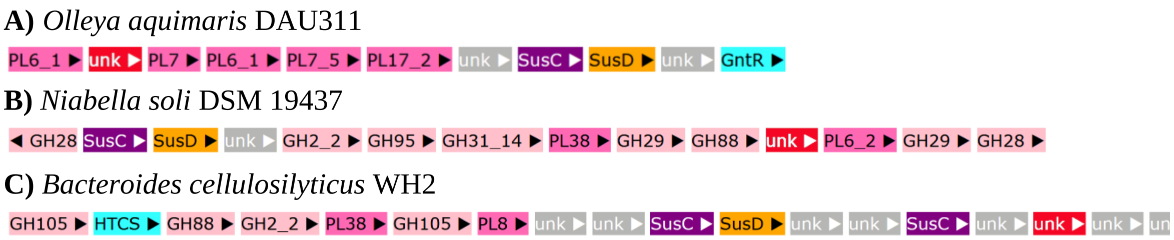


**Additional Figure 6: PUL organizations** around DZC78_07865 from the marine bacterium *Olleya aquimaris* DAU311, alginate (**A**); and the two characterized Bacteroidota members: Niaso_2009 in *N. soli* (B), and BcellWH2_01176 in *B. cellulosilyticus WH2* (C), all indicated in red.

by marine species as exemplified by its presence along with PL6, PL7 and PL17 in the marine bacterium *Olleya aquimaris* DAU31 (Additional Figure 6A). While this strongly supports the alginate lyase activity discovered in this work, the two additionally characterized members show more diverse PUL organizations. In the soil bacterium *Niabella soli* (Additional Figure 6B), the PUL includes not only PL6 and PL38 members but also notably α-fucosidases (GH29 and GH95), α-galacturonidases (two GH28), an α-galactosidase (GH31_14) and a β-glucuronidase (GH2_2). The PUL of the human gut *Bacteroides cellulosilyticus* (Additional Figure 6C) also contains PL38 and a GH2_2 members, as well as a noteworthy PL8, family mostly reported to recognize and cleave glucuronic acid residues in various glycans. These support PL_New2 intervention in the degradation of glycans containing, among others, guluronic acids.
