## Additional File 2 for "Screening for Polysaccharide Utilization Loci Targeting Marine Polysaccharides"

### In-depth characterization of a GH_New2 family member: B2K_10610

#### A. Results

##### A.1. Structural characterization of B2K_10610

The structure of native B2K_10610 was solved at 2.05 Å resolution by molecular replacement using an AlphaFold model[1] as template. Additional Table 1 provides the refinement statistics of B2K_10610 structure.

In the crystal structure, B2K_10610 adopts a dimeric form, predicted as a stable state by PISA analysis[2]. The interface between the two monomers covers an interaction surface area of 2208 Å², stabilized by an array of hydrogen bonds and salt bridges. Specifically, the 12 salt bridges involve His156, Lys173, His174, Arg181, Asp188, Asp190, Glu196, His201, Arg203, Asp542, Lys638, and Glu668 (Additional Figure 6A-B). These residues are not conserved across the GH_New2 family, suggesting that other family members could adopt a monomeric state and that dimerization is not crucial for catalysis.

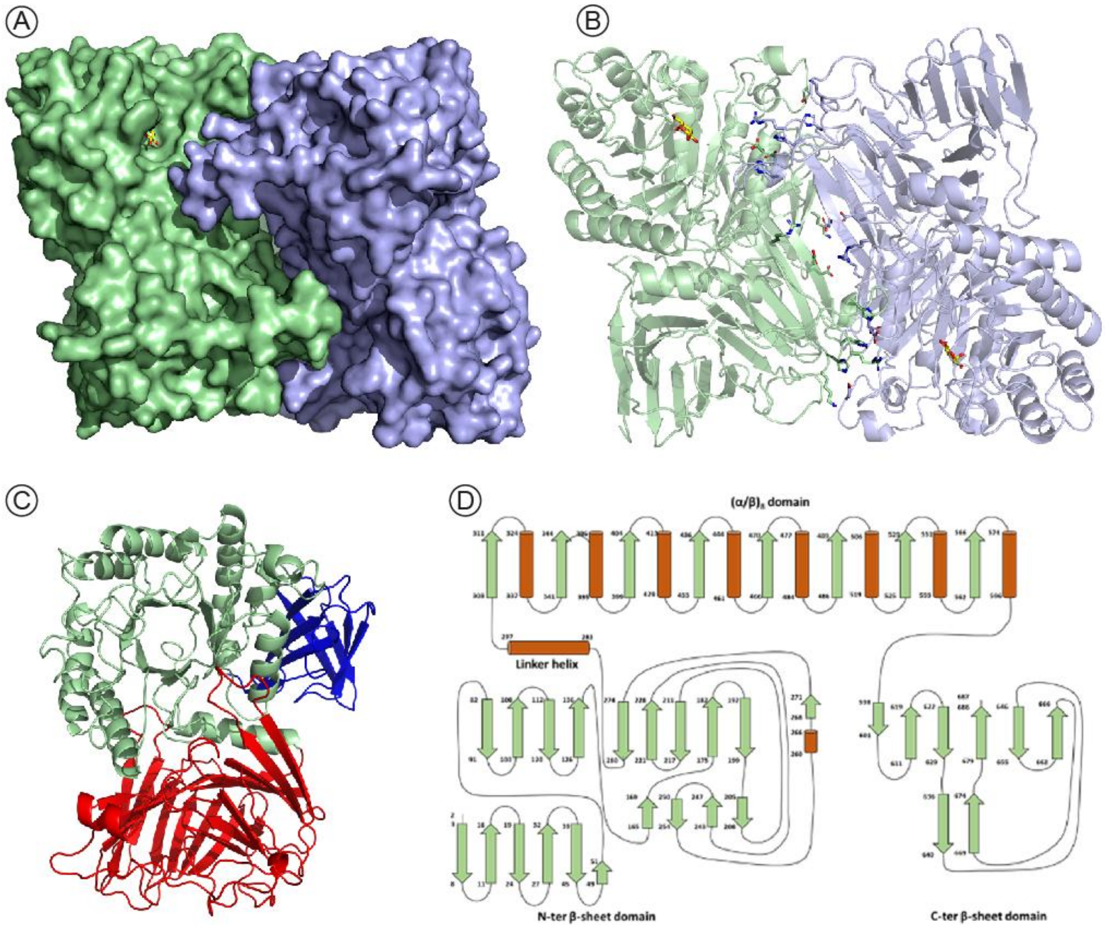

**Additional Figure 6: Crystal structure of α-galactosidase B2K_10610.** (**A** and **B**, idem Figure 4 A and B) Overall structure of the B2K_10610 dimer. Amino acids involved in intermolecular salt bridges shown as sticks. (**C**, **D**) Architecture of B2K_10610 monomer. Three domains are highlighted: the N-terminal β-sheet domain (red), the central TIM barrel domain (green), and the C-terminal β-sheet domain (blue).

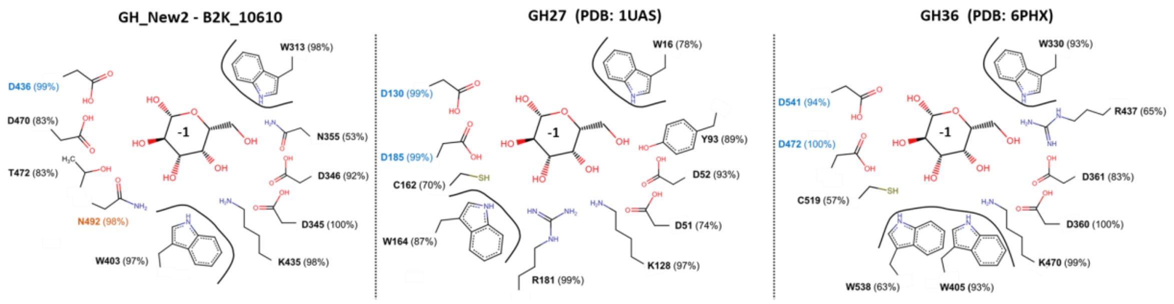

**Additional Figure 7 - Architecture and residue conservation of the -1 subsite in α-galactosidase B2K_10610 (GH_New2), 1UAS (GH27), and 6PHX (GH36)**. Conservation percentages correspond to residue frequencies within each respective enzyme family.

B2K_10610 comprises three folded domains: an N-terminal β-sandwich domain, an $(\alpha/\beta)_8$ central domain, and a C-terminal β-sandwich domain (Additional Figure 6C-D). The N-terminal β-sandwich is a large domain, comprising 281 amino acids that form 18 β-strands. The TIM barrel is connected to the N-terminal domain by an extended α-helix. The C-terminal domain forms a small β-sandwich composed of 91 amino acids arranged into 8 antiparallel β-strands. This overall structure should be conserved within the family, based on the high sequence similarity among its members.

To obtain insight into substrate recognition we co-crystallized the B2K_10610 D437A inactive mutant with *p*NP-α-D-galactopyranoside (α-Gal-pNP). Unfortunately, during the long incubation time of the co-crystallization assay α-Gal-pNP underwent spontaneous hydrolysis and only galactose molecules could be observed in the active site of each monomer of the B2K_10610-α-Gal crystal structure, obtained at 1.50 Å resolution (Main Figure 4C). The active site is located at the central axis of the $(\alpha/\beta)_8$ barrel domain and displays a "pocket" structure with only one negative subsite (-1). This structure arrangement supports the exo-acting activity of B2K_10610. The folding of this cavity involves several loops, including those connecting α-helices and β-sheets within the TIM domain, as well as a specific loop from the N-terminal β-sheet domain spanning residues 182-192. Interestingly, B-factor analysis shows low values of all these except the loop (346-386) suggesting the low dynamic of this protein's region. It is however worthwhile mentioning that no major structural rearrangement of the active site scaffold could be observed above binding of α-Gal in the chain A monomer, whereas in the chain B monomer the loop (346-386), connecting β2 with α2 in the TIM barrel, is partially disordered in the native structure, but undergoes a movement of about 1.2 Å upon substrate binding, thereby closely wrapping around the α-Gal substrate and providing stabilizing enzyme-substrate interactions.

The galactose molecule observed in the -1 subsite is stabilized through a combination of hydrogen bonds and aromatic stacking interactions (Main Figure 4C and Additional Figure 7, left panel). Notably, Trp313 and Trp403 are likely involved in the aromatic stacking, contributing to the glycan stabilization. The C2 hydroxyl group of galactose forms hydrogen bonds with Asp470 Oδ2, Thr472 Oγ1, and Asn492 Oδ1 atoms. Additionally, Asn492 plays a dual role by coordinating the C3 hydroxyl group via its Nδ2 atom, in collaboration with Lys435 Nζ atom. The C4 hydroxyl group establishes hydrogen bonds with

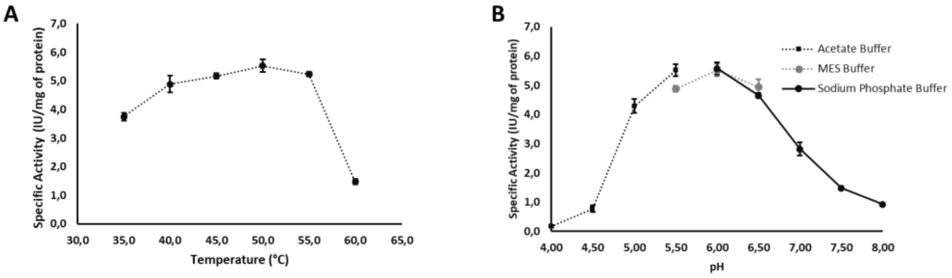

C

| | $k_{cat}$ ($s^{-1}$) | $K_M$ (mM) | $V_{max}$ ($\mu M.s^{-1}$) | $k_{cat}/K_M$ ($mM^{-1}.s^{-1}$) |
|---|---|---|---|---|
| **B2K_10610 - WT** | 17,94 ± 0,45 | 7,56 ± 0,45 | 0,897 ± 0,45 | 2,37 |
| **B2K_10610 - D437A** | n.a. | n.a. | n.a. | n.a. |
| **B2K_10610 - D470A** | 0,0024± | 7,09 ± 0,8 | 0,0024 ± 0,0001 | $3,38.10^{-4}$ |
| **B2K_10610 - N492A** | 0,014± | 89,95 ± 18,89 | 0,014 ± 002 | $1,56.10^{-4}$ |

**Additional Figure 8 - Biochemical characterization of B2K_10610.** For all the experiments, pNP-α-galactopyranoside was used as substrate. Effect of temperature (A) and pH (B) on α-galactosidase activity, and kinetic parameters (C) of wild-type and putative inactive mutants under optimal conditions were obtained by averaging three experiments with bars indicating ± the standard deviation.

Lys435 Nζ and Asp345 Oδ2. Finally, the C6 hydroxyl group interacts with Asp346 Oδ2 and Asn355 Nδ2 atoms. All these residues are highly conserved within the GH_New2 family, except Asn355, highlighting their importance in substrate recognition and the specificity of this family.

#### A.2. Biochemical characterization

During initial high-throughput screening, activity on *p*NP-α-D-galactopyranoside was detected for B2K_10610. This activity was further investigated. The enzyme exhibited maximum activity at pH 6.0 and 50°C (Additional Figure 8A-B). Under these conditions, B2K_10610 displayed a $K_M$ value of 7.56 mM and a catalytic efficiency of 2.37 $mM^{-1}\cdot s^{-1}$ (Additional Figure 8C). This α-galactosidase activity was further evaluated on natural substrates containing a terminal α-galactopyranose, including 6-α-D-galactopyranosyl-D-glucopyranose (melibiose) and 3-α-D-galactopyranosyl-D-galactoyranose (Gal-α1.3-Gal), using HPAEC-PAD. B2K_10610 released a galactose molecule from melibiose (Main Figure 4D) and, in lower amount, from Gal-α1,3-Gal (Additional Figure 9) demonstrating its clear exo-α-galactosidase activity.

To further investigate the molecular mechanism of B2K_10610, we analyzed the kinetic parameters of the several putative inactive mutants: D437A, D470A and N492A (Additional Figure 8C). The selection of single-point mutations was guided by the X-ray structure, specifically considering the distance between the anomeric carbon of the galactose molecule and potential catalytic residues.

Structural analysis revealed that the Oδ2 atom of Asp437 is positioned 2.3 Å from the C1 atom of galactose, making it well-located to act as the nucleophilic residue via its carboxylate group. Consistently, the D437A substitution completely abolished the enzymatic activity, confirming the essential catalytic role of Asp437. Interestingly, the D470A mutant retained the same $K_M$ as the wild-type but exhibited a dramatically reduced $k_{cat}$, indicating that Asp470 contributes to catalysis rather than substrate binding. Its proximity with Asp437 (putative nucleophile) and its orientation support that this residue could act as a pKa modulator, by assisting the nucleophile residue and promoting a proper protonation state for the catalysis.

Asn492 participates in coordinating the C2 and C3 hydroxyl groups of the substrate within the active site and is highly conserved across the GH_New2 family. The N492A mutant exhibited a tenfold reduced affinity for *p*NP-galactopyranoside, highlighting its critical role in substrate recognition. Interestingly, the N492A mutant also exhibited a $k_{cat}$ more than 1000-fold lower than the wild-type, strongly suggesting its critical role in catalysis. These observations support the hypothesis that Asn492 participates in the proton relay network and facilitates glycosidic bond cleavage[3].

##### A. 3. Comparison of GH_New2 with closely related GH27 and GH36

Remote evolutionary relationships were detected between GH_New2 and the GH27 and GH36 families at both sequence and structural levels and, interestingly, all exhibit α-galactosidase activities. However, important differences were observed notably in their overall domain architecture and in the residues involved in galactose recognition within the active site. The B2K_10610 tripartite domain architecture (N-terminal β-sheet, central $(\beta/\alpha)_8$ TIM barrel, and small C-terminal β-sheet) is highly similar to available crystal structures of the GH36 family. This tripartite organization is highly conserved across GH_New2 and GH36 families, but the sequences of their N-terminal domains show high divergence. Interestingly, GH27 proteins completely lack the N-terminal domain, suggesting family-specific structural adaptations.

GH_New2, GH27 and GH36 families also show conservation and divergence in the amino acids shaping the -1 subsite of their active site, involved in galactose recognition (Additional Figure 7). All three families feature: (i) two Asp and one Lys residues to coordinate the C6 and C4 hydroxyl groups; and

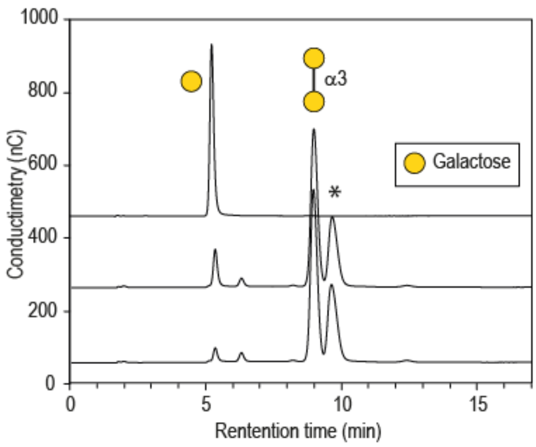

**Additional Figure 9 - HPAEC–PAD analysis of B2K_10610 activity.** Galactose was used as standards. Gal-α1,3-Gal was incubated with B2K_10610**.** * Peak attributed to HEPES buffer.

(ii) two conserved Trp residues playing an essential role in stabilizing the galactose ring through π–π stacking. Members of family GH27 display a highly conserved Tyr residue coordinating the C6 hydroxyl group, whereas members of families GH36 and GH_New2 display less conserved Arg and Asn residues, respectively. Furthermore, GH27 members exhibit a unique and conserved Arg interacting with the C3 hydroxyl group, making its active site more positively charged compared to the members of the other families. Regarding the catalytic amino-acids, all three families show high conservation of the structural positions of the two Asp residues playing the role of nucleophile and pKa modulator. The principal structural divergence at the -1 subsite between GH_New2 and the GH27/GH36 families lies in the residues positioned proximal to the C2 hydroxyl group and the anomeric carbon of galactose. GH27 and GH36 show similar Asp acid/base, while GH_New2 is characterized by a unique Asn and Thr (Asn492 and Thr472 in B2K_10610) occupying this region.

Altogether, these variations illustrate how evolution can fine-tune enzyme families with functional conservation, while family-specific adaptations remain to be further investigated. Consequently, GH_New2 will be classified as a novel family part of the CAZy clan GH-D.

#### B. Experimental procedures

##### B.1. Protein expression and purification

Like wild-type B2K_10610, its three mutants (D437A, D470A and N492A) were synthesized and cloned into pET28a vectors (Twist bioscience). The N-terminal His-tagged recombinant proteins were expressed in *Escherichia coli* Lemo21(DE3) competent cells (New England Biolabs). Cultures were grown in ZYP 5052 medium for 22 hours at 25 °C. The 200 mL culture were centrifuged, then pellets were resuspended in 40 mL of lysis buffer (50 mM phosphate buffer pH 7.4, 300 mM NaCl, 0.5 $mg.mL^{-1}$ Lysozyme) and stored at -80 °C. For the protein purification, resuspended bacterial pellets were disrupted by sonication. Soluble extracts were applied on 5 mL HisTrap columns (Cytiva) pre-equilibrated with 50 mM sodium phosphate buffer pH 7.4 and 300 mM NaCl. Resins were washed with 100 mL of 50 mM imidazole, 50 mM sodium phosphate pH 7.4 and 300 mM NaCl. Proteins were eluted using 30 mL of 50 mM sodium phosphate buffer pH 7.4 containing 250 mM imidazole and 300 mM NaCl. The elution fractions were analyzed by SDS-PAGE then pooled. Size exclusion chromatography was performed using an AKTA express system and a Superdex S200 16/200 column, upstream the biochemical characterization. Proteins were eluted in HEPES buffer 50 mM pH 7.0, using a flow of 1.5 $mL.min^{-1}$. Purified fractions were pooled together and concentrated using AmiconUltra-15 with a cut-off of 10 kDa. The enzyme concentration was determined by measuring $A_{280}$, 144 075 $M^{-1}$ $cm^{-1}$.

##### B.2. Enzymology

Enzyme kinetics assays were conducted in a 500 µL reaction mixture containing 5 mM *p*NP-α-Gal (Carbosynth), 50 mM sodium acetate, MES or sodium phosphate buffer 1 mg/mL BSA, and 50 µM enzyme. Optimum pH was determined using 50 mM sodium acetate buffer (pH 4.0, 4.5, 5.0, 5.5), MES buffer (pH 5.0, 5.5, 6.0) and sodium phosphate (pH 6.0, 6.5, 7.0, 7.5 and 8.0) at 50 °C. Optimal temperature was evaluated under the same conditions as above, in 50 mM sodium phosphate buffer at pH 6.0, across a temperature range from 30 °C to 60 °C. For all experiments, 50 µL of the reaction mixture was stopped in 50 µL of 1 M $Na_2CO_3$ every 2 min and during 16 min to ensure that the reaction velocity equaled to the initial velocity. Absorbance at 405 nm was measured at each time point using a PHERAstar FSX microplate reader (BMG Labtech) and converted to the equivalent amount of released galactose.

For wild-type B2K_10610, kinetic parameters were determined by varying the substrate concentration of *p*NP-α-Gal (0.05, 0.1, 0.25, 0.5, 0.75, 1, 1.75, 2.5, 4, 5, 7.5, 10, 12.5, 15, 17.5, and 20 mM) in 50 mM sodium phosphate buffer at pH 6.0 and 50 °C, using a final enzyme concentration of 50 nM. For B2K_10610 mutants, the kinetic assays were conducted using 10 µM enzyme and substrate concentrations of 5, 10, 20, 40, 60, 80, and 100 mM in 50 mM sodium phosphate buffer at pH 6.0 and 50 °C. Reactions were stopped as described above every 30 min and during 240 min. All quantitative assays were expressed as the mean ± SD from three independent experiments. Kinetic parameters were calculated using GraphPad Prism software (8.0.2).

**B.3. HPAEC-PAD analysis**

The activity of B2K_10610 on natural substrates was evaluated by High-Performance Anion-Exchange Chromatography with Pulsed Amperometric Detection (HPAEC-PAD), using a CarboPac™ PA100 guard column in series with a CarboPac™ PA100 analytical column (2 mm × 250 mm), at a flow rate of 0.250 $mL \cdot min^{-1}$. Enzyme kinetics were assessed using melibiose, type B antigen tetrasaccharide and galactopyranoside-α-1,3-galactopyranose substrates. Reactions were carried out in 200 µL of sodium phosphate buffer (pH 6.0) containing 1 µM enzyme and 1 mM substrate, at 40 °C during 1 hour. Reactions were stopped by heating the mixture at 90 °C for 10 min. 20 µL of each reaction mixture were diluted 10 times, and 10 µL of the diluted sample were injected. For HPAEC-PAD, Eluent A consisted of 150 mM NaOH, while Eluent B was 150 mM NaOH with 500 mM sodium acetate. Sugars were eluted at 20 °C using a linear gradient from 0% to 30% Eluent B over 50 min, followed by a 5 min wash with 100% Eluent B and a 10 min re-equilibration with 100% Eluent A. Peaks were identified by comparing retention times with those of individual monosaccharide standards (10 $mg \cdot L^{-1}$) injected separately.

**B.4. Crystallization and X-ray structure determination**

Crystallization trials were performed by the sitting-drop vapour-diffusion method in SwissCi crystallization plates (STP labtech) using a Mosquito® (STP labtech) crystallization robot and commercial screens. For the native protein at a concentration of 12 mg/ml crystal emerged from a condition of the JCSG+ Screen (Molecular Dimensions) containing 10% (w/v) PEG 8000, 0.1 M Hepes buffer pH 7.5 and 8% (v/v) ethylene glycol. The B2K_10610 D437A at 15 mg/ml was incubated with

30 mM α-Gal-PNP in 20 mM Hepes buffer at pH 7.0. Crystals were obtained with the JCSG Core I screen (Qiagen) from a condition containing 0.2 M lithium sulfate, 0.1 M TRIS buffer pH 8.5 and 30% (w/v) PEG 4000. Crystals were cryo-protected with reservoir solution supplemented with 15% (v/v) glycerol *prior* cooling in liquid nitrogen. Crystals of native B2K_10610 and the D437A mutant in complex with galactose belong to space group $P2_12_12_1$ with average unit cell axes of 97 × 108 × 143 Å and two molecules per asymmetric unit. X-ray diffraction data were acquired at beam lines Proxima1 and Proxima2 at the Synchrotron SOLEIL, Gif-sur-Yvette. All diffraction data were processed with XDS[4] and scaled and merged using the CCP4 (Winn *et al*, 2011) suite of programs Pointless, Aimless[5] and Truncate. The structure of native B2K_10610 was solved by molecular replacement with the program Phaser[6] using an Alphafold[1] model as search model (rotation function Z-score: 6.7; translation function Z-score: 21.5; number of clashes from packing analysis: 0; log likelihood gain: 3714). Maximum-likelihood refinement, including TLS refinement, and model adjustment were carried out with the programs Refmac[7] and Coot[8], respectively. For the dataset arising from native B2K_10610 a random set of 5% of reflections was set aside for cross-validation purposes. The cross-validation set for the B2K_10610 D437A galactose complex has been taken over from the parent (native B2K_10610) data set and randomly extended to higher resolution. Model quality was assessed with internal modules of Coot[8] and with the Molprobity server[9]. Figures were generated with Pymol[10] accessible *via* SBGRID[11]. The atomic coordinates and structure factors of native B2K_10610 and B2K_10610 D437A in complex with galactose have been deposited in the Protein Data Bank[12] with accession numbers 29RM, 29SO, respectively. Data collection and refinement statistic are provided in Additional Table 1.

**B.5. Sequence Analysis**

Sequences encoding families GH_New2, GH27 and GH36 were extracted from GeneBank (June 2025). To remove redundancies, sequences with over 95% identity were filtered out with CD-HIT version 4.8.1[13], and truncated sequences were manually curated. The final dataset comprised 1,340 GH27 sequences, 1,899 GH36 sequences, and 92 GH_New2 sequences. Multiple sequence alignments were generated using MAFFT webserver[18], and residue conservation scores were subsequently obtained with Jalview version 2.11.5.0 (Procter et al., 2021).

**Additional Table 1: Structural data collection and refinement statistics**

| | **B2K_10610 native** | **B2K_10610 D437A galactose complex** |
|---|---|---|
| PDB entry | 29RM | 29SO |
| **Data collection** | | |
| Synchrotron beam line | SOLEIL-Proxima2 | SOLEIL-Proxima1 |
| Space group | $P2_12_12_1$ | $P2_12_12_1$ |
| Cell dimensions□, *a*, *b*, *c* (Å) | 97.62, 107.39, 143.50 | 96.92, 108.60, 142.33 |
| Resolution (Å) | 48.81-2.05 (2.09-2.05) | 47.44-1.50 (1.53-1.50) |
| No. unique reflections | 95116 (4663) | 238020 (11687) |
| CC (1/2) | 0.997 (0.628) | 0.999 (0.614) |
| $R_{meas}$ | 0.214 (2.117) | 0.087 (1.672) |
| *I* / □*I* | 11.0 (2.0) | 17.1 (1.6) |
| Completeness (%) | 99.9 (99.7) | 99.5 (99.7) |
| Redundancy | 13.4 (13.3) | 12.9 (12.1) |
| Wilson B ($A^2$) | 29.2 | 18.0 |
| **Refinement** | | |
| Resolution (Å) | 48.86-2.05 (2.103-2.05) | 47.42-1.50 (1.539-1.50) |
| No. reflections working set | 90181 (6607) | 226091(16569) |
| No. reflections test set | 4842 (355) | 11806 (856) |
| $R_{work}$ / $R_{free}$ | 0.1915/0.2255 | 0.1630/0.1884 |
| No. atoms | | |
| Protein | 11084 | 11146 |
| Ligands/Ions | 68/2 | 62/4 |
| Waters | 503 | 1441 |
| B-factors (Å$^2$) | | |
| Protein main-chain/side-chain | 30.32/35.37 | 18.86/22.94 |
| Ligands/Ions | 44.91/60.28 | 28.63/31.57 |
| Waters | 34.45 | 31.67 |
| R.m.s. deviations | | |
| Bond lengths (Å) | 0.007 | 0.009 |
| Bond angles (°) | 0.998 | 1.185 |
| Ramachandran | | |
| Favoured (%) | 97.05 | 96.79 |
| Allowed (%) | 2.95 | 3.21 |
| Disallowed (%) | 0 | 0 |

Values in parentheses for the highest resolution shell
