## Additional File 3 for "Screening for Polysaccharide Utilization Loci Targeting Marine Polysaccharides"

### Structural analyses of the S1_30 *Bacteroides xylanisolvens* ι-carrageenan sulfatase

#### A. Results

In order to contribute to the understanding of substrate recognition in S1 enzymes we undertook further studies on *Bacteroides xylanisolvens* ι-carrageenan-sulfatase (hereafter *Bx*S1_30) which has been classified into sub-family S1_30 in the SulfAtlas database[1,2]. *Bx*S1_30 is predominantly monomeric in solution, with minor dimeric species according to our size exclusion chromatography assays (not shown), possibly resulting from heterologous expression.

##### A.1 Structural studies of *Bx*S1_30

To the best of our knowledge no three-dimensional structure has been reported so far of any member of the S1_30 sub-family. Crystal structures of a *Bx*S1_30 inactive mutant, for which the active site serine residue to be post-translationally modified to an FGly was mutated to an alanine, were solved by the molecular replacement method from *Bx*S1_30_S85A crystals diffracting X-rays to 1.41- and 1.7 Å resolution, respectively. Both crystallized in the monoclinic space group *C*2, with one molecule present in the asymmetric unit for *Bx*S1_30_S85A without ligand, whereas that with substrate bound displayed two molecules in the asymmetric unit. As all other S1 sulfatase structures determined to date[3], *Bx*S1_30 displays a two-domain structure with the N-terminal domain adopting an alkaline phosphatase α/β/α-fold formed by residues 21-397 and the C-terminal domain comprising residues 398-482 which

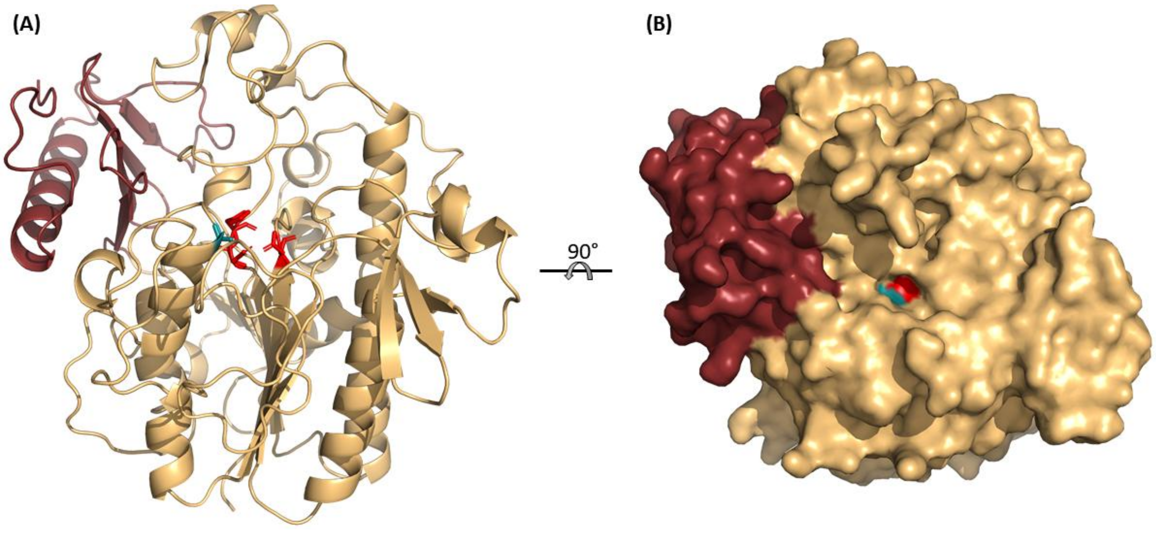


**Supplementary Figure 1**: Crystal structure of *B. xylanisolvens* ι-carrageenan sulfatase (only monomer A is shown). (**A**) The ribbon diagram is colored as a function of domains. The N-terminal domain (amino-acid residues 21-397) colored in light orange contains the catalytic domain. This domain hosts the active site and active site residues are colored in red (D32, D316, H317) and cyan (S85A). The smaller C-terminal domain (amino-acid residues 398-482) is colored in ruby. (**B**) Surface presentation and top-view of the two-domain structure rotated approximately 90° around the horizontal axis as compared to (**A**).

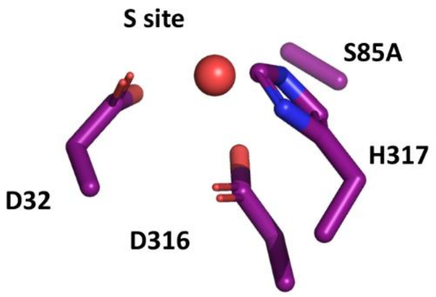


**Supplementary Figure 2: Calcium** binding site below the S site in the crystal structure of *B. xylanisolvens* ι-carrageenan sulfatase. The calcium ion has been substituted with a water molecule, and the active site serine has been mutated to an alanine (S85A).

form a four stranded anti-parallel β-sheet being situated in between an α-helix (also C-terminal domain) and the N-terminal domain (Supplementary Figure 1).

The active site is situated in the N-terminal domain and comprises residues Asp32, Ser85 (herein mutated to alanine), Asp316 and His317, with Ser85 being the residue to be converted into an FGly upon post-translational modification). This latter is a part of the S1 sulfatases signature motif, C/S-X-P-X-R[4]. As seen in Supplementary Figure 1, the active site region is flanked with flexible loops. The N-terminal domain also hosts the so-called S site (sulfate binding site), which in the major part of the known structures holds a bound calcium ion. Interestingly, in the two structures described herein of the S85A mutant, no calcium seems to be bound to this site in which a water molecule has been observed at the calcium ion position (Supplementary Figure 2). This latter performs hydrogen bonding interactions with Asp32, Asp316 and His317.

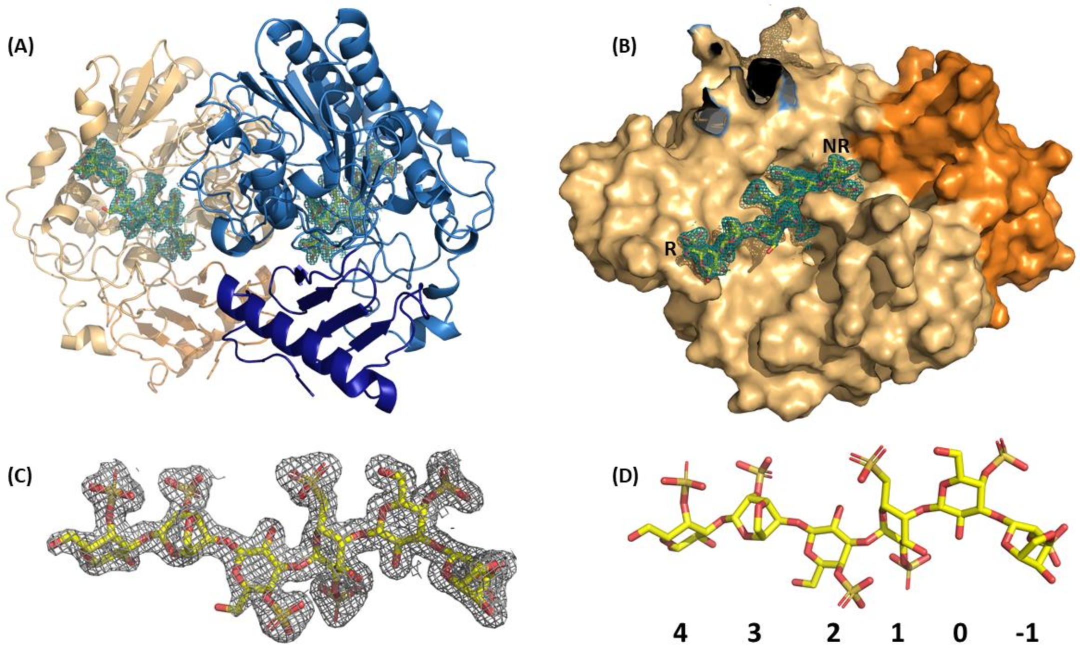


**Supplementary Figure 3:** Neo-ι-ν-ι-carrahexaose bound in the active site of *B. xylanisolvens* ι-carrageenan sulfatase. **A**) Ribbon diagram of the dimer colored as a function of domains. C-terminal domains are colored in dark orange (molA) and dark blue (molC), respectively, and the N-terminal domains in light orange (molA) and skyeblue (molC) contain the catalytic domain. This domain hosts the active site to which ινι-NC6 (shown with 2Fo-Fc electron density in green around the DP6) is bound within a long cleft with the non-reducing (NR) end pointing towards the C-domain (**B**). **C**) Close-up on the 2Fo-Fc electron density in grey contoured at 1σ around ινι-NC6. **D**) Stick-representation of ινι-NC6 and corresponding sub-sites.

In order to inform on substrate recognition and processing of substrate, data from crystals of *B. xylanisolvens* ι-carrageenan sulfatase (here *Bx*S1_30_S85A) grown in the presence of neo-ι-carrahexaose (ι-NC6) were analyzed. The structure solved by the molecular replacement method, employing the S85 mutant structure without substrate bound as search model, indicated that two molecules are present in the asymmetric unit. Only one direct interaction between the two monomers is observed with the sidechain of Glu140 performing hydrogen bonding to the backbone N of Asp139 of the other monomer. This is in agreement with PISA interface analysis[5] suggesting that the interface plays no role in formation of the complex, and likely is due to the crystal packing. When comparing this crystal structure (monomer A) to that without substrate bound in the active site, as expected, the structures were highly similar displaying an RMSD of 0.265 Å based on 422 atoms. For the *Bx*S1_30_S85A/DP6 complex, both molecules displayed a bound hexa-saccharide in the electron density in the active site region. Surprisingly, when studying the electron density corresponding to the DP6 in details, the density corresponded to a molecule of neo-ι-/ν-/ι-carrahexaose (hereafter referred to as ινι-NC6), and not ι-NC6 as expected (Supplementary Figure 3). This is explained by the composition of the DP6 fraction obtained after enzymatic degradation of ι-carrageenan which contains low amounts of ν-carrabiose, the biosynthetic precursor of ι-carrageenan. The purified DP6 fraction is composed of standard neo-ι-carrahexaose (ι-NC6) and hybrid ι-/ν-/ι- carrahexaose (ινι-NC6)[6].

When analyzing the crystal structure in details, most interactions between the enzyme and the substrate (Supplementary Table 1) are performed, by far, in subsites 0 (G4S) and +1 (D2S6S). The 4-sulfate of the G4S moiety present in the 0 subsite is coordinated in the S subsite by Asn106, Lys133, His135, His164, His225, His317, Lys329 and the water molecule substituting the calcium ion (Supplementary Figure 4). The alanine residue substituting the catalytic serine (S85A) is positioned

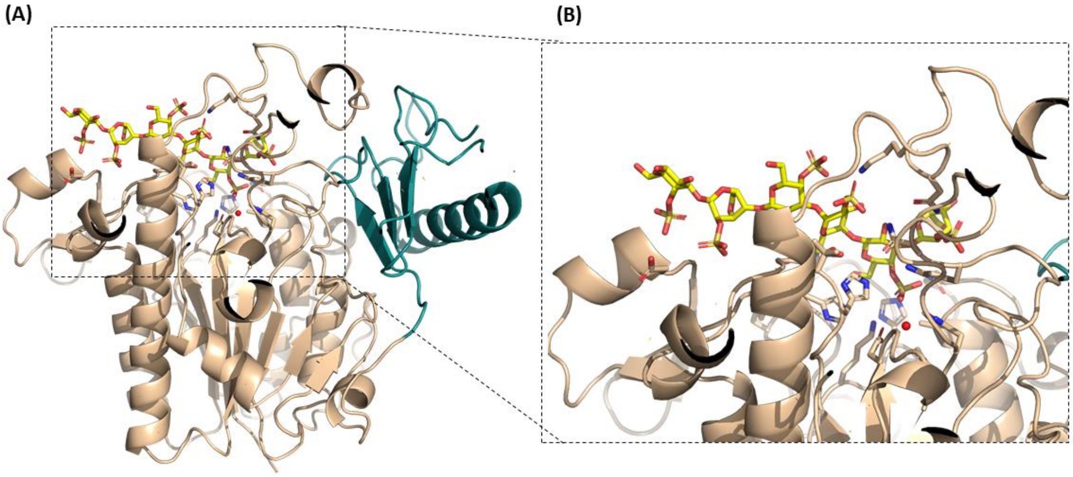


**Supplementary Figure 4.** Interactions between ινι-NC6 and *B. xylanisolvens* ι-carrageenan sulfatase are majorly salt-bridges and hydrogen bonds. (**A**) Overall structure of a monomer (N-terminal domain in wheat; C-terminal domain in deepteal) (A) highlighting amino acid residues participating in ινι-NC6 binding as sticks, and (**B**) close-up on the active site region.

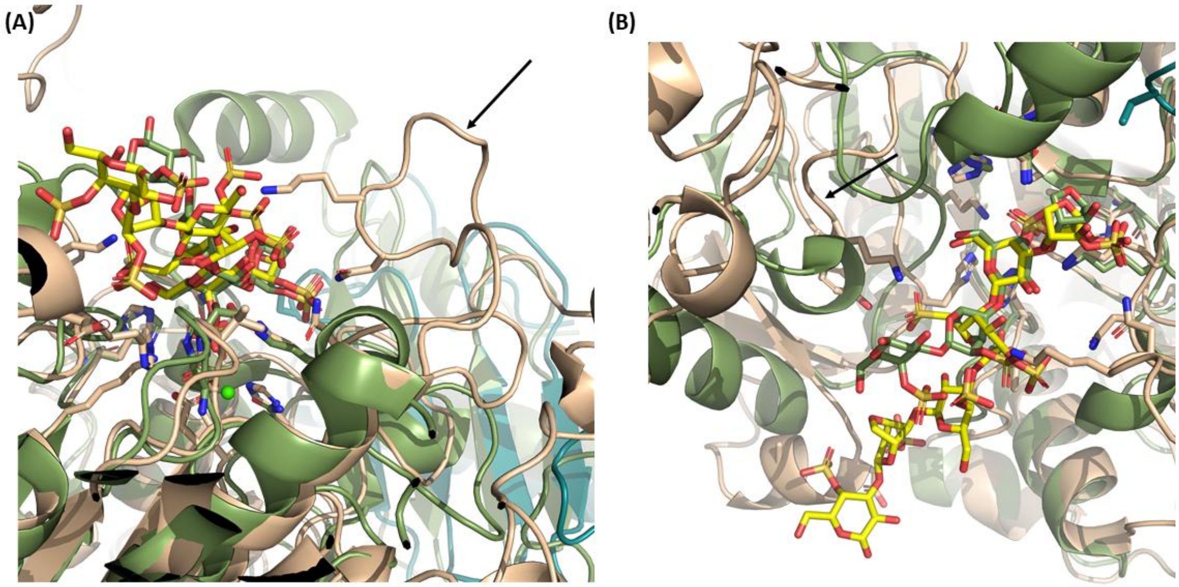


**Supplementary Figure 5**. Overlay of the crystal structure complex of *Ps*S1_19A/ι-NC4 (green) onto that of *Bx*S1_30_S85A/ ιvι-NC6 (N-terminal domain in wheat, C-terminal domain in teal). (**A**) Interactions in subsite -1 (but also subsite +1) are performed by residues situated in a loop (indicated by a black arrow) which is lacking in *Ps*S1_19A/ι-NC4. (**B**) Interactions in subsite +1 also are partly performed by residues situated in a loop (indicated by a black arrow) not present at the same position in *Ps*S1_19A/ι-NC4.

~3.15 Å below the targeted sulfate ester, and His225 is positioned ~2.9 Å from the scissile bond in an orientation being in agreement with the protonation of the ester oxygen, thereby making it a candidate as the catalytic acid. It should be noticed that Asn106 is present in a double conformation in the structure without ligand bound and that the imidazole ring of His135 changes orientation upon substrate binding. As concerns the non-reducing end 2-sulfate of 3,6-anhydro-D-galactose (DA2S) positioned in subsite -1, it is relatively loosely held with two hydrogen bonding interactions to the Lys329 peptide oxygen and nitrogen, but also to the Asn269 sidechain. For this latter, the carboxamide group is slightly shifted towards the substrate (~1.5 Å) as compared to the substrate free structure. For the remaining part of the hexaose (subsites +1 to +4), especially subsite +1 reveals that a high number of hydrogen bonds and salt-bridges stabilize the D2S6S moiety, involving Lys142, Tyr143, His164, Thr226, Lys268 and Asn 328 (Supplementary Table 1). As seen in the 0 subsite an asparagine carboxamide group (here Asn328) is shifted ~1.5 Å towards the substrate compared to the substrate free structure, and Thr226 adopts a double conformation in the complex. It can equally be highlighted that the G4S at the reducing end (subsite +4) is stabilized by interactions *via* the enzyme backbone (Gly180 and Ala183). Interestingly, no interactions are present between the substrate and residues from the C-terminal domain for which the closest residue, Cys 407, is situated approximately 6 Å from the non-reducing end 2-O-sulfated 3,6-anhydro-D-galactose.

#### A.2. Comparative studies of *B. xylanisolvens* ι-carrageenan sulfatase with ι-carrageenan sulfatases from other sub-families

In order to contribute to the understanding of the ι-carrageenan specificity of *Bx*S1_30, we compared the S1_30 three-dimensional structure of *Bx*S1_30 with those of an endo-4S-ι-carrageenan sulfatase from *Pseudoalteromonas sp*. PS47 (*Ps*S1_19A)[7] classified into S1 _19 and an S1_81 sulfatase (formerly referred to as *Pf*S1_NC)[8]. When superposing the structure of *Bx*S1_30_S85A/ ινι-NC6 with that of *Ps*S1_19A endo-4-sulfo-D-galactose sulfatase in complex with ι-NC4 (PDB 6B1V) and which also has been reported to be active on ι-carrageenan, the rmsd is 1.728 Å based on 244 Cα atoms. Careful inspection of the interactions in subsite -1, indicates that the interaction of Lys329 (Lys309 in *Ps*S1_19A) with the DA2S unit is conserved, whereas the interaction of Asn328 residing in a loop region in the three-dimensional structure of *Bx*S1_30 does not exist in *Ps*S1_19A since there is no loop here in this latter. This very loop seems to be responsible in big part for the recognition of the D2S6S moiety in subsite +1 in that the sidechain of Lys268 interacts with its 2-sulfate. A somewhat similar situation is

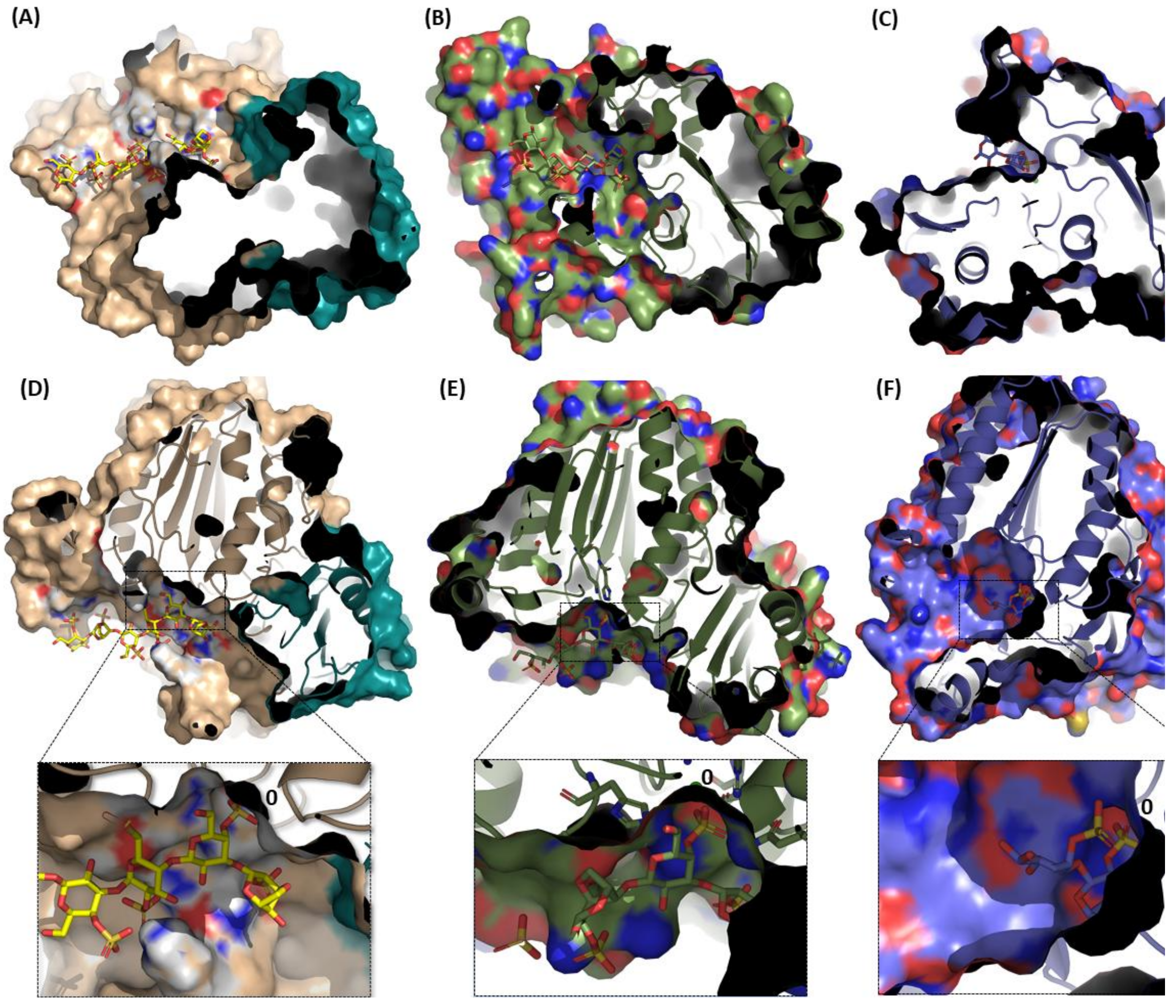


**Supplementary Figure 6.** Surface representations (**A,D**) *Bx*S1_30_S85A/ινι-NC6 with the C-terminal domain in teal, (**B,E**) *Ps*S1_19A/ι-NC4 and (**C,F**) *Pf*S1_81_C84S/ι-NC2. (**B**) Close-up on the substrate binding region with focus on the 0 subsite pocket. (**F**) The molecule is slightly rotated as compared to (**D**) and (**E**) to better appreciate the pocket shape and the fact that the entire sugar moiety, a DA2S, is located in the pocket.

observed when comparing the region of Lys142 and Tyr143, both interacting with the 6-sulfate of D2S6S. Indeed, these two amino acids are situated in a loop region radically different from what is observed in the same region in *Ps*S1_19A (Supplementary Figure 5). As concerns subsite 0 all residues involved in interactions with the substrate are conserved with the exception of His317 which is substituted by Asn292 in *Ps*S1_19A.

Superposition of the crystal structure of *Pseudoalteromonas fuliginea* exo-2-sulfo-3,6-D-anhydro-galactose sulfatase in complex with ι-NC4 (S1_81) onto that of *Bx*S1_30_S85A/ινι-NC6 indicates a rmsd of 2.226 Å based on 264 Cα atoms. It is complicated to make a direct comparison in that the substrate is different from those observed in *Bx*S1_30 and in *Ps*S1_19A, and for which G4S moieties are in subsite 0. In the *Pf*S1_81_C84S complex in which a ι-NC2 was observed, the non-reducing DA2S moiety is located in subsite 0. The amino acid residues which in the *Bx*S1_30_S85A/ινι-NC6 complex performed interactions with the hexa-saccharide in subsite 0 are all conserved in *Pf*S1_81_C84S/ι-NC2 with the exception of His164 for which no amino acid is present at that position and His317 which is substituted by Gln334. Indeed, the differences in substrate recognition and binding can be appreciated three enzymes are by comparing the surfaces of the three enzymes (Supplementary Figure 6).

Whereas *Pf*S1_81_C84S/ι-NC2 clearly has a narrow pocket corresponding to subsite 0 in which DA2S is complemented by the shape of the active site pocket, characteristic of exo-acting enzymes, the two other present large clefts to accommodate longer substrates confirming their endo activity.

Altogether, these studies have contributed to shed light on the understanding of the substrate specificity of different enzymes being active on a same substrate. We have reported the first structure of an S1_30 enzyme, and have shown that this latter has a higher specificity for ινι-neocarrahexaose than for ι-neocarrahexaose. We highlighted amino acid residues being essential for the recognition of ι-neocarrahexaose and its precursor ινι-neocarrahexaose and showed that the calcium ion needed for catalysis is weakly bound in that both crystal structures reported herein lacked this cation.

#### B. Experimental procedures

##### B.1. Protein expression and purification

A codon-optimized gene encoding for N-terminal 6xHis-tagged *B. xylanisolvens* ι-carrageenan sulfatase mutated on position serine 85 to an alanine, was synthesized and cloned into a pET-15b vector (Novagen) by GenScript. The resulting plasmid was transformed into *E. coli* T7 Express lysY/Iq cells (NEB, C3013). Cells were cultivated in LB medium supplemented with appropriate antibiotic (kanamycin and chloramphenicol or ampicillin) at 37 °C and 150 rpm. When the $OD_{600}$ reached 0.5, protein expression was induced by the addition of 0.5 mM isopropyl β-D-1-thiogalactopyranoside (IPTG) and cultures were incubated overnight (16-18 h) at 25 °C and 150 rpm. Cells were harvested by centrifugation (8200 × g for 15 min at 4 °C) and resuspended in lysis buffer (20 mM HEPES pH 7.5, 300 mM NaCl, 30 mM Imidazole, 10 µg/mL DNase I, 1 µg/mL of lysozyme and protease inhibitor cocktail: 1 µg/mL Chymostatin, 1 µg/mL Leupeptine, 1 µg/mL Antipain, 1 µg/mL Pepstatin, 5 µg/mL Aprotinin), and lysed by sonication. Following centrifugation (14500 × g for 10 min at 4 °C), the clarified supernatant was applied to a HisTrapTM HP column (Cytiva) using an ÄKTA pure™ system (Cytiva). The column was washed with lysis buffer and with 2 M NaCl, and the protein was eluted using a gradient up to 500 mM imidazole, followed by immediate desalting and buffer exchange into 20 mM HEPES (pH 7.5) with 150 mM NaCl using HiPrep™ columns (Cytiva). The protein was further purified using a Superdex® 200 Increase 10/300 GL column (Cytiva) pre-equilibrated with a buffer containing 20 mM HEPES (pH 7.5) and 150 mM NaCl. The eluate was collected, concentrated and buffer exchanged (20 mM HEPES pH 8.0, 150 mM NaCl) using an Amicon® Ultracel-30K centrifugal filter (Millipore) and applied to a HiTrap™ SP HP cation exchange chromatography column (Cytiva). The elution was performed by a linear gradient of 0–0.5 M NaCl in the same buffer. Fractions judged to be ≥95% pure by SDS–PAGE were combined, desalted and concentrated as described above. The protein concentration was determined by measuring the absorbance at 280 nm using the molar extinction coefficient calculated by ProtParam on the ExPasy server (https://web.expasy.org/protparam/).

##### B.2. Crystallization and X-ray crystallography studies

Crystallization experiments were carried out using a protein sample of recombinant *Bx*S1_30_S85A purified as described above, and for which monodispersity was confirmed by dynamic light scattering, DLS (Zetasizer Nano-S, Malvern) prior to crystallization screening. Crystals for X-ray diffraction were grown at 292 K using the sitting drop vapor diffusion technique employing a Mosquito® crystallization robot (SPT Labtech). All crystals were set up in 96-well MRC crystallization plates (Molecular Dimensions). For S85A mutant crystals, 100 nL of *Bx*S1_30_S85A at a concentration of 35 mg/mL was mixed with 100 nL reservoir solution containing 18 % w/v PEG 3350 and 100 mM sodium citrate tribasic-dihydrate buffer (pH 5.5), and equilibrated against 70 µL reservoir solution. Crystals appeared after 28 days. For crystals *Bx*S1_30_S85A grown in the presence of ι-NC6 (molecule synthesized as described earlier[6]), 35 mg/mL of the enzyme was pre-mixed with 10 mM of the hexaose. Hereafter, 100 nL of this mixture was mixed with 100 nL of precipitant solution containing 20% w/v PEG 3350 and 200 mM sodium malonate pH 6.0, and equilibrated against 70 µL of this reservoir solution. For crystals grown in the presence of ι-NC6, growth started at day 7, and the final size was reached upon 23 days. All crystals were mounted in cryo-loops and were flash frozen without adding any cryo-protectants for data collection.

##### B.3. Data collection and structure determination

X-ray diffraction data were collected on ESRF beamline ID30B (doi: 10.15151/ESRF-DC-2406973792), and data reduction was performed using programs from the autoPROC toolbox[9] and the

XDS package[10]. The Phenix suite (Adams et al., 2010) was used for structure determination employing the Phaser-MR module[11] for molecular replacement with the AlphaFold2 model of *Bx*S1_30[12] "AF-A0A7J5PXU8-F1-model_v4" as a search model. A single partial solution with an LLG of 1016 was obtained. The refined structure of *Bx*S1_30_S85A without ligand bound was used as search model to solve the structure of *Bx*S1_30_S85A in complex with ιvι-NC6 to 1.41 Å resolution. A single solution with two molecules in the asymmetric unit was found displaying an LLG of 34147. Three-dimensional structure refinement was done using phenix.refine[13] alternating with model building using COOT[14]. The structure of *Bx*S1_30_S85A was refined to R = 17.92 % and $R_{free}$ = 21.16 %, and that of the complex to R = 15.66 % and $R_{free}$ = 17.65 %, respectively.

##### B.4. Interaction analysis

Initial analyses of interactions between the enzyme and the substrate were performed using the PLIP (Protein Ligand Interaction Profiler) web server[15], followed by manual validation.

##### B.5. Figure rendering

Figures of three-dimensional structures were generated with PyMol (DeLano Scientific LLC, http://pymol.sourceforge.net/).

##### Accession codes

Coordinates and structure factors have been deposited in the RCSB Protein Data Bank under accession codes 30BD (*Bx*S1_30_S85A mutant without substrate) and 30SR (*Bx*S1_30_S85A mutant in complex with neo- ιvι-carrahexaose substrate).

**Supplementary Table 1.** Interactions between *Bx*S1_30 and ινι-NC6.

| **Subsite** | **ινι-NC6** | ***Bx*S1_30** | **Interaction** |
|---|---|---|---|
| -1 | DA2S | Asn269 Nδ2 | Hydrogen bond |
| -1 | DA2S | Lys329 N | Hydrogen bond |
| -1 | DA2S | Lys329 O | Hydrogen bond |
| 0 | G4S | Asp32 Oδ2 | Hydrogen bond |
| 0 | G4S | Ala85 N | Hydrogen bond |
| 0 | G4S | Asn106 Nδ2 | Hydrogen bond |
| 0 | G4S | Lys133 Nζ | Salt-bridge |
| 0 | G4S | His135 Nδ1 | Salt-bridge |
| 0 | G4S | His164 Nε2 | Hydrogen bond |
| 0 | G4S | His225 NE2 | Salt-bridge |
| 0 | G4S | His317 NE2 | Salt-bridge |
| 0 | G4S | Lys329 Nζ | Salt-bridge |
| 0 | G4S | Water substituting $Ca^{2+}$ | Hydrogen bond |
| +1 | D2S6S | Lys142 Nζ | Salt-bridge |
| +1 | D2S6S | Tyr143 | Hydrogen bond |
| +1 | D2S6S | His164 N | Hydrogen bond |
| +1 | D2S6S | Thr226 Oγ1 | Hydrogen bond |
| +1 | D2S6S | Lys268 Nζ | Hydrogen bond |
| +1 | D2S6S | Lys268 Nζ | Salt-bridge |
| +1 | D2S6S | Asn328 Nδ2 | Hydrogen bond |
| +2 | G4S | Lys268 Nζ | Salt-bridge |
| +3 | DA2S | Asp181 N | Hydrogen bond |
| +4 | G4S | Gly180 N | Hydrogen bond |
| +4 | G4S | Gly180 O | Hydrogen bond |
| +4 | G4S | Ala183 N | Hydrogen bond |

**Supplementary Table 2.** Data collection and refinement statistics.

| | ***Bx*S1_30_S85A** | ***Bx*S1_30_S85A/ neo-ι-ν-ι-carrahexaose** |
|---|---|---|
| **PDB ID** | **30BD** | **30SR** |
| Beamline | ID30B | ID30B |
| Wavelength (Å) | 0.87313 | 0.87313 |
| Resolution range (Å) | 63.93 – 1.70 (1.73 – 1.70) | 68.13 – 1.41 (1.45 – 1.41) |
| Space group | *C*2 | *C*2 |
| Unit cell dimensions<br>a, b, c (Å)<br>α, β, γ (°) | <br>95.30, 85.44, 73.94<br>90 120.16 90 | <br>110.72, 88.47, 122.73<br>90, 105.27, 90 |
| Unique reflections | 110296 (6750) | 431939 (31912) |
| Multiplicity | 3.8 (3.6) | 3.4 (2.8) |
| Completeness (%) | 99.3 (99.6) | 99.7 (99.3) |
| Mean I/σ(I) | 12.0 (2.6) | 7.7 (1.0) |
| †$R_{merge}$ | 0.175 (0.563) | 0.067 (0.919) |
| $CC_{1/2}$ | 0.765 | 0.998 (0.398) |
| **Refinement** | | |
| Protein atoms | 3684 | 7349 |
| Solvent atoms | 587 | 1388 |
| Ligand atoms | - | 92 (ινι-NC6) |
| $R_{work}$ | 0.1518 | 0.1566 |
| £$R_{free}$ | 0.1731 | 0.1765 |
| r.m.s.d. bonds (Å) | 0.006 | 0.010 |
| r.m.s.d. angles (°) | 0.817 | 1.129 |
| Average B-factor ($Å^2$) | 19.42 | 24.27 |
| Ramachandran plot<br>Favoured (%)<br>Allowed (%)<br>Outliers (%) | <br>97.4<br>2.6<br>0.0 | <br>97.1<br>2.9<br>0.0 |

Values in parentheses correspond to outermost resolution shells.

All diffraction data were collected on single crystals.

†$R_{merge} = \Sigma_{hkl}\Sigma^{N}_{i=1} |I_{hkl,i} - \langle I\rangle_{hkl}|/\Sigma_{hkl}\Sigma^{N}_{i=1} |I_{hkl,i}|$, where $I_{hkl}$ is the intensity of a reflection and $\langle I\rangle_{hkl}$ is the mean intensity of the reflection, and N is the number of observations of that reflection.

£$R_{free}$ calculated from 5% of the data excluded from refinement.
