## Supplementary figures and images for "Screening for Polysaccharide Utilization Loci Targeting Marine Polysaccharides"

### Figure_S2

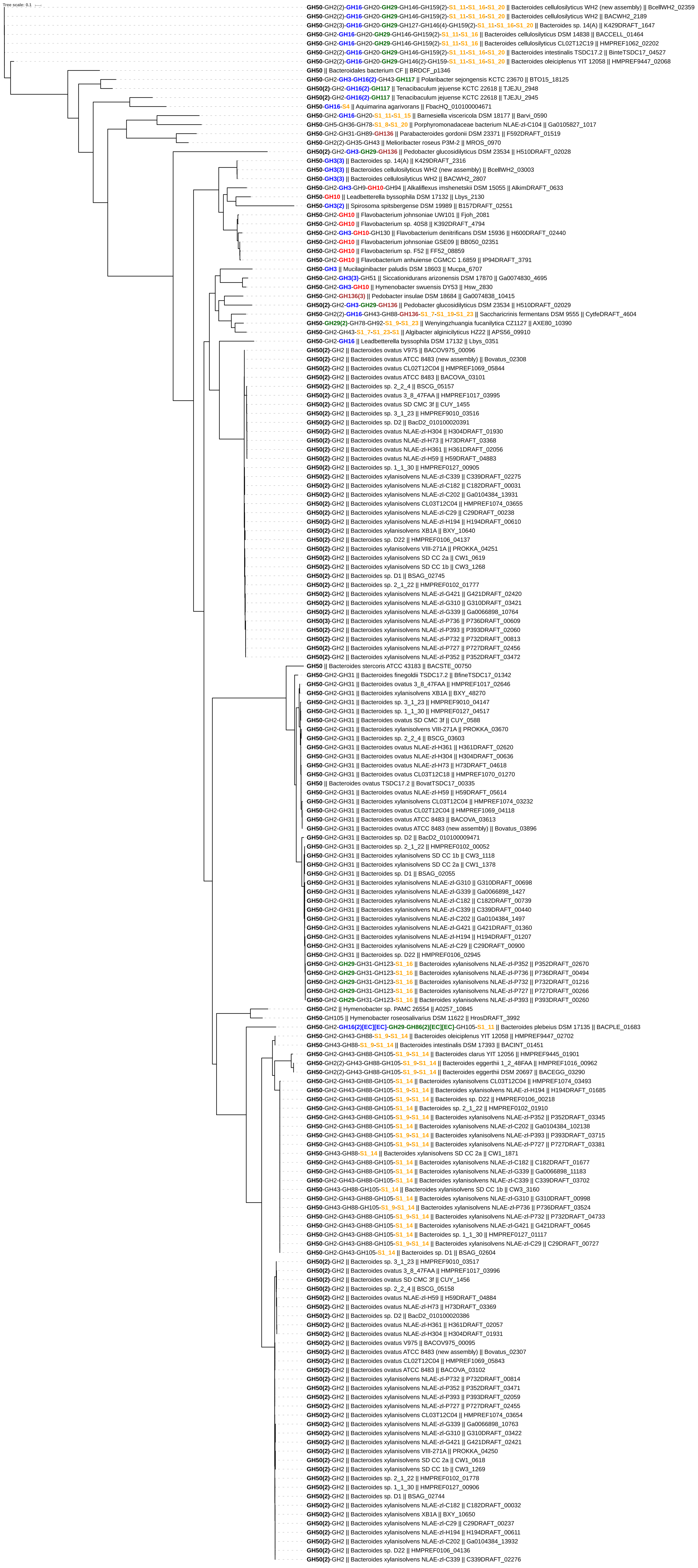

### Figure_S3

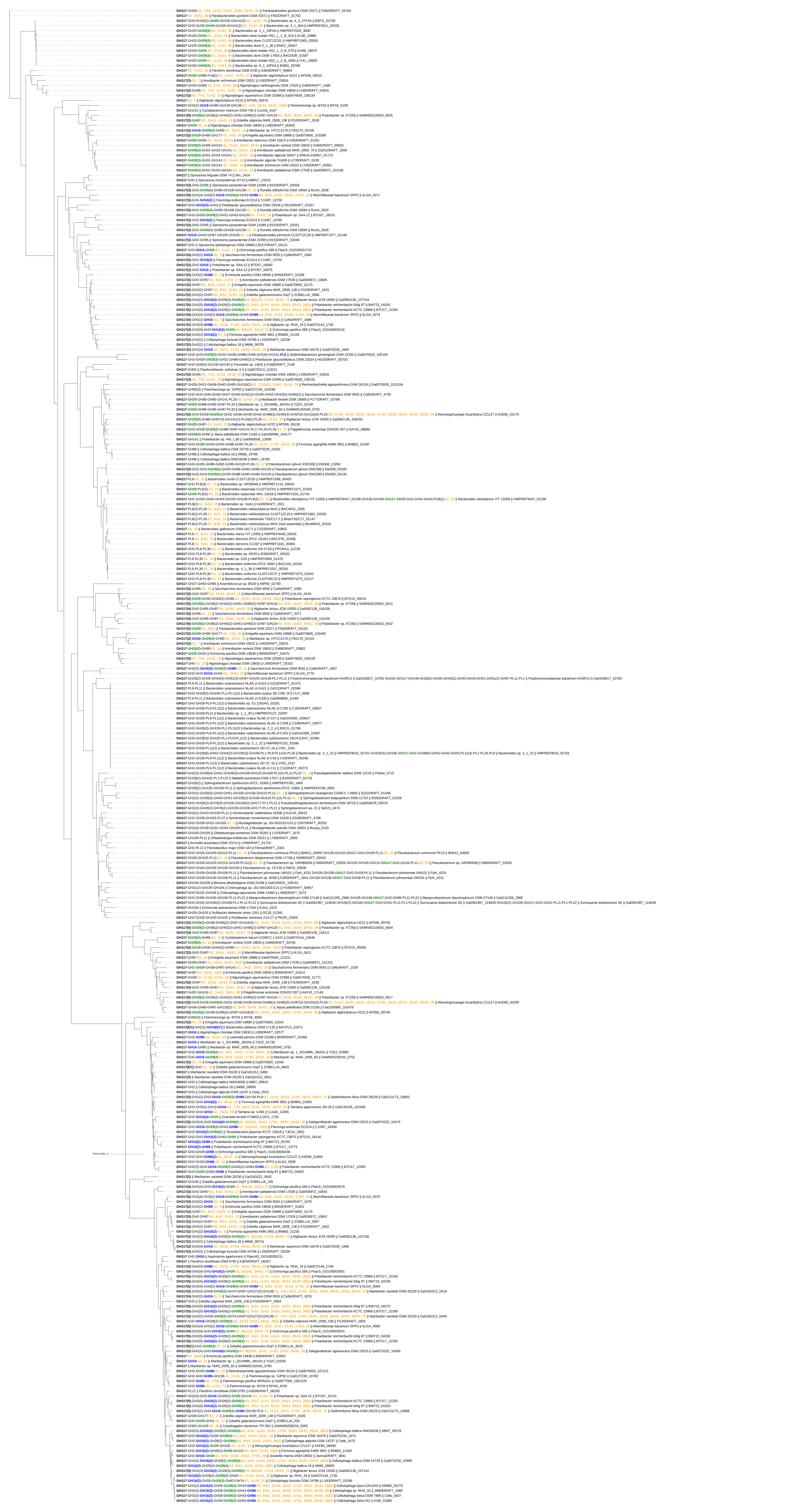

### Figure_S7

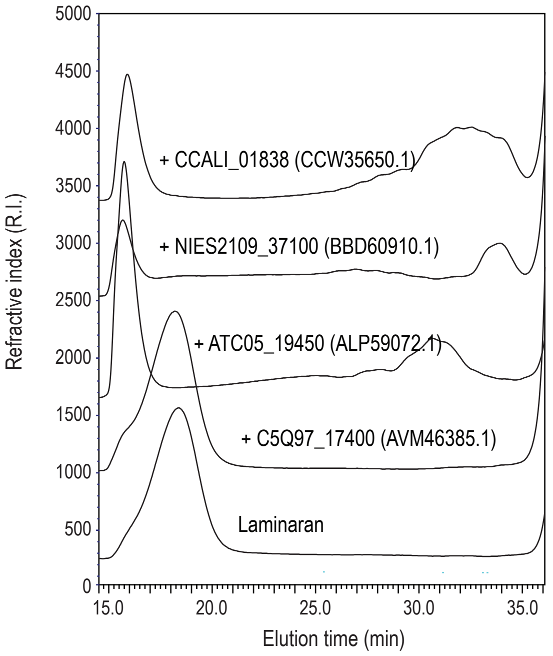


**Figure S7:** Size exclusion chromatography of laminaran incubated with selected GH50 enzymes.
