## Supplementary material for "Screening for Polysaccharide Utilization Loci Targeting Marine Polysaccharides": Figure_S4

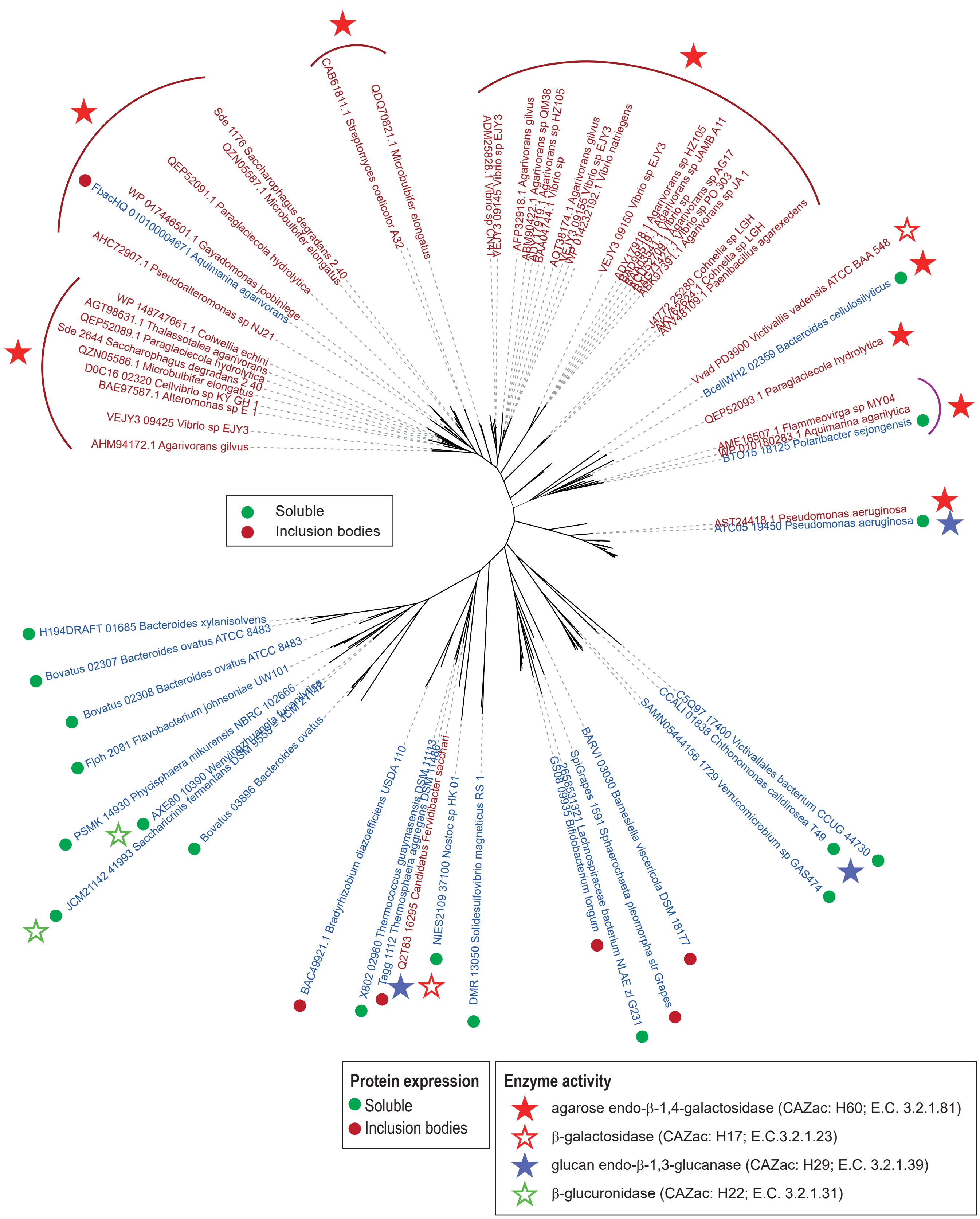


**Figure S4:** Diversity of the GH50 family. The phylogenetic tree was calculated from members of the GH50 family. Branch labels (loci and bacteria) in red correspond to enzymes studied and described in the literature. Branch labels in blue correspond to enzymes investigated in this study.
