## Supplementary material for "Screening for Polysaccharide Utilization Loci Targeting Marine Polysaccharides": Figure_S5

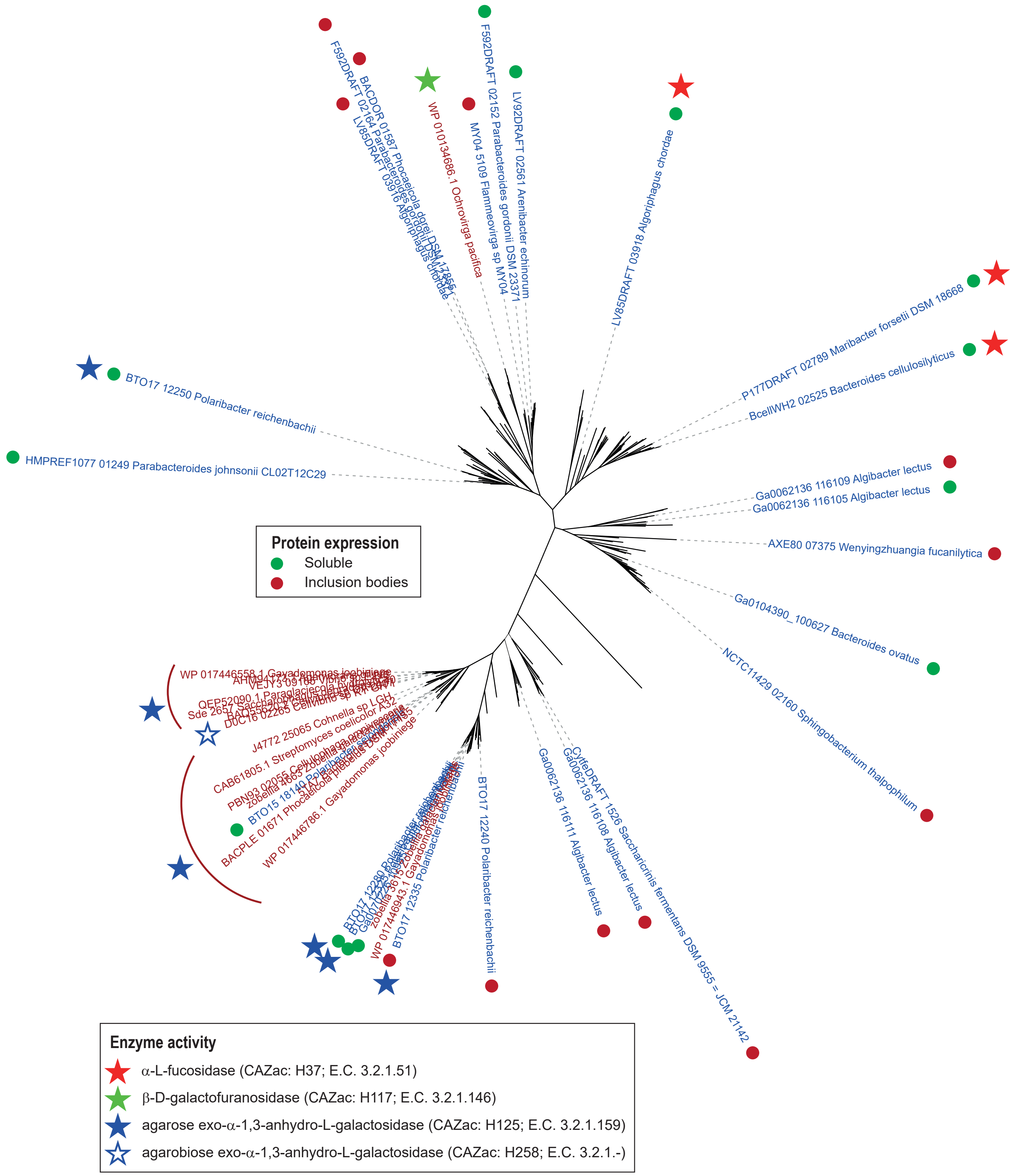


**Figure S5:** Diversity of the GH117 family. The phylogenetic tree was calculated with members of the GH117 family. Branch labels (loci and bacteria) in red correspond to enzymes reported in the literature. Branch labels in blue correspond to enzymes investigated in this study.
