## Supplementary material for "Screening for Polysaccharide Utilization Loci Targeting Marine Polysaccharides": Figure_S6

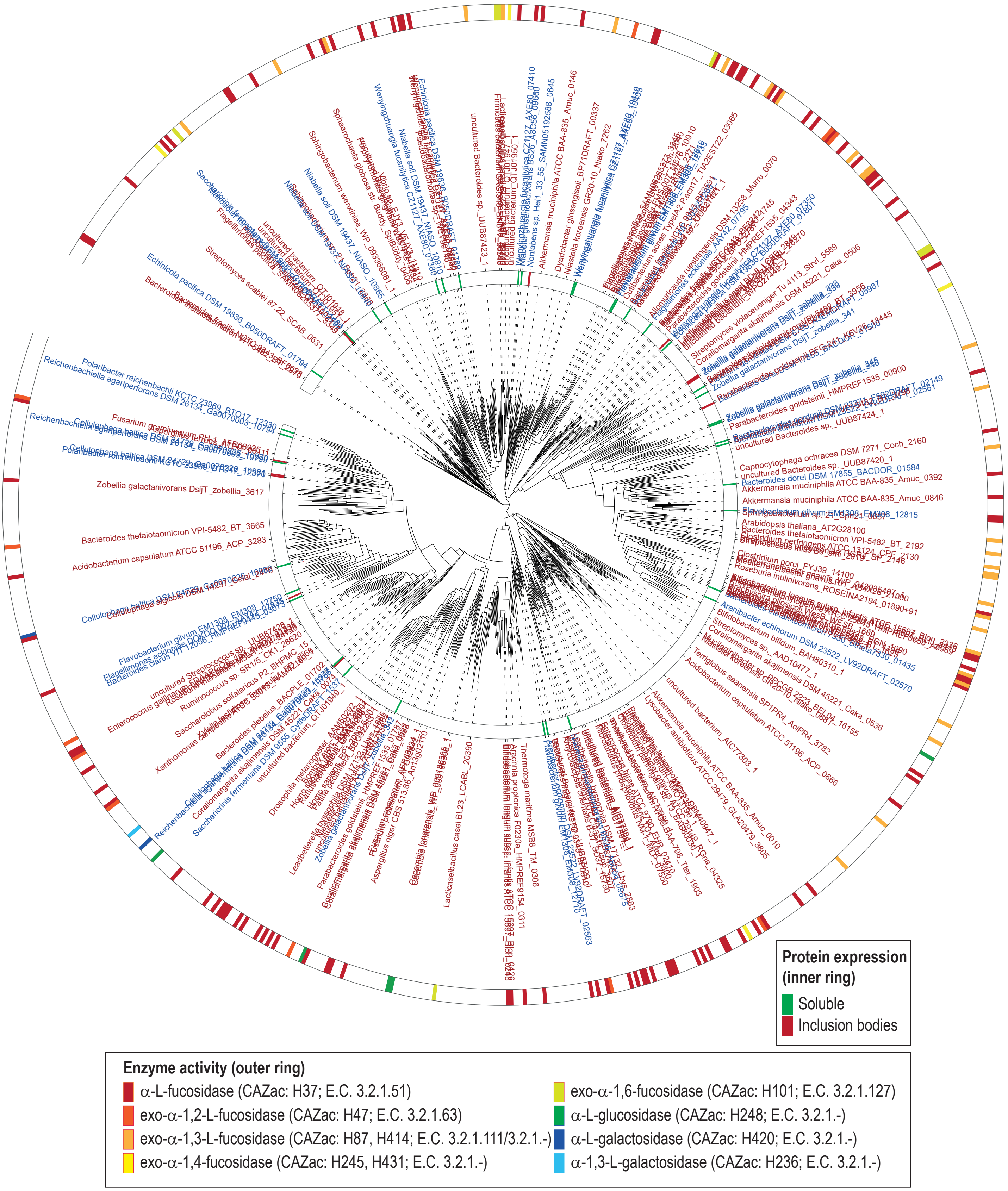


**Figure S6:** Diversity of the GH29 family. The phylogenetic tree was calculated with members of the GH29 family. Branch labels (loci and bacteria) in red correspond to enzymes studied and described in the literature. Branch labels in blue correspond to enzymes invetigated in this study.
