## Supplementary material for "Screening for Polysaccharide Utilization Loci Targeting Marine Polysaccharides": Figure_S8

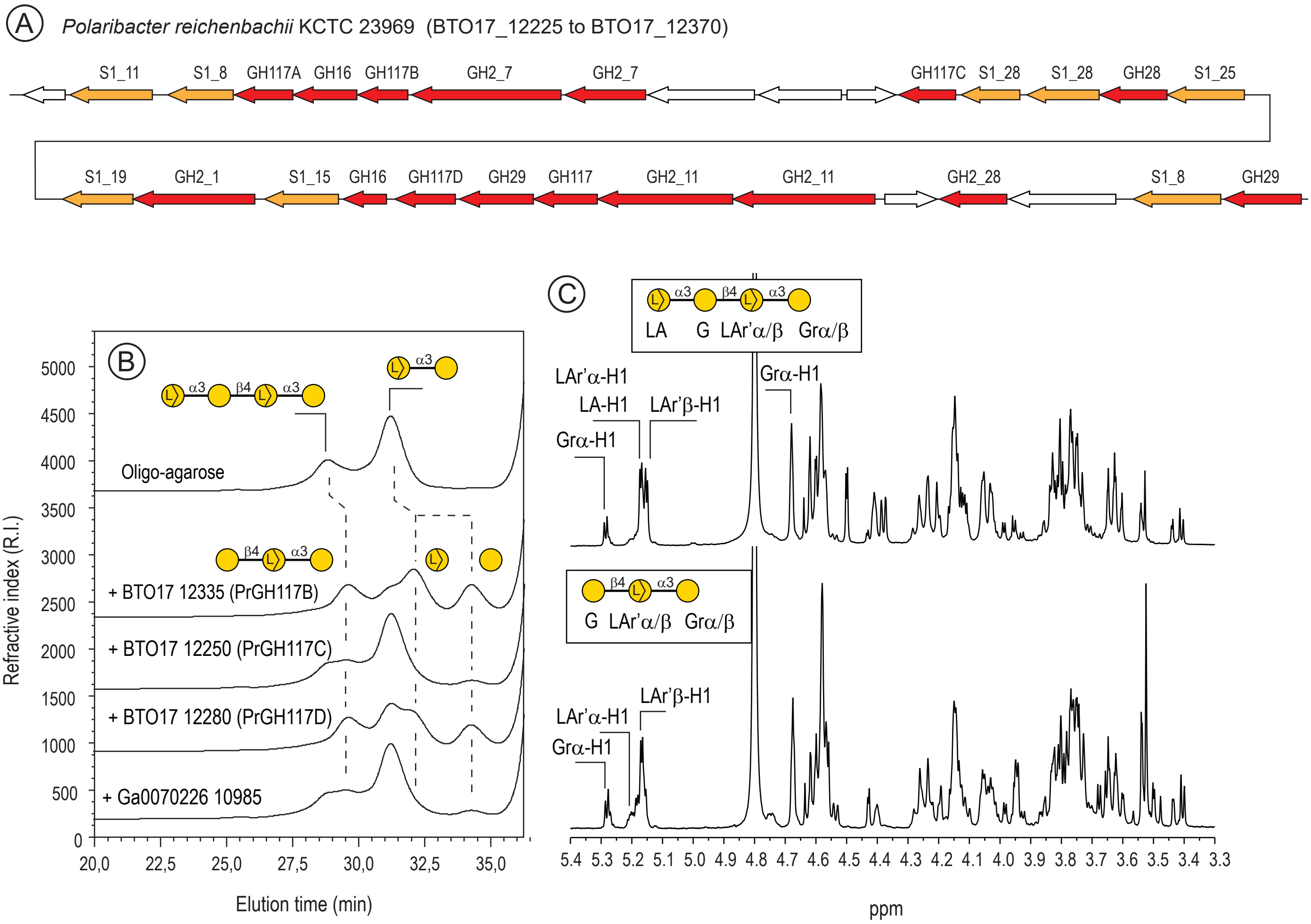


**Figure S8: A**) Organization of the selected PUL 18 dedicated to the degradation of agarose and prophyran. The PUL contain four enzymes grouped in the GH117 family. **B**) Chromatogram recorded after incubation of neo-oligoagarose with selected GH117 enzymes. **C**) $^{1}H$ NMR recorded on neo-agarotetraose inubated with α-L-anhydrogalactose hydrolase.
