## Supplementary material for "Screening for Polysaccharide Utilization Loci Targeting Marine Polysaccharides": Figure_S9

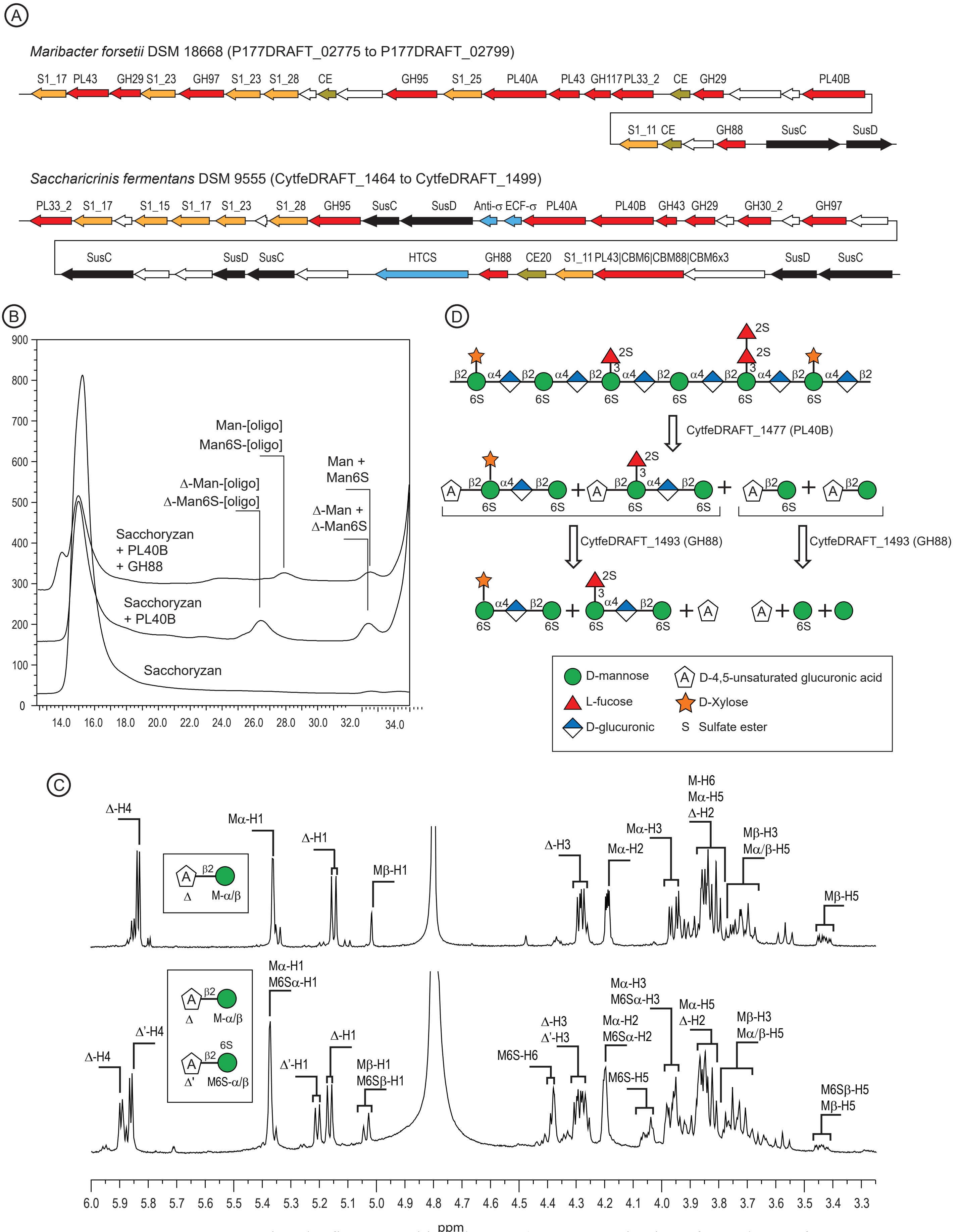


**Figure S9:** PUL encompassing the first saccorhizan lyase. **A**) PUL organization of *Maribacter forsetii* DSM 18668 and *Saccharicrinis fermentans* DSM 9555 dedicated to the degradation of saccorhizan. **B**) Size exclusion chromatography of saccorhizan incubated with the PL40 saccorhizan lyase. The oligo-saccharides obtained were further digested by the GH88 glucuronyl hydrolase. **C**) 1H NMR of the end-product of the saccorhizan lyase demonstrating the cleavage of the  -linked mannose and the formation an unsaturated residue at the non-reducing ends. **D**) Partial degradation pathway of the degradation pathway of sacchorizan. Other enzymes and especially sulfatases were not expressed solubles and actives.
