## Supplementary material for "Screening for Polysaccharide Utilization Loci Targeting Marine Polysaccharides": Figure_S10

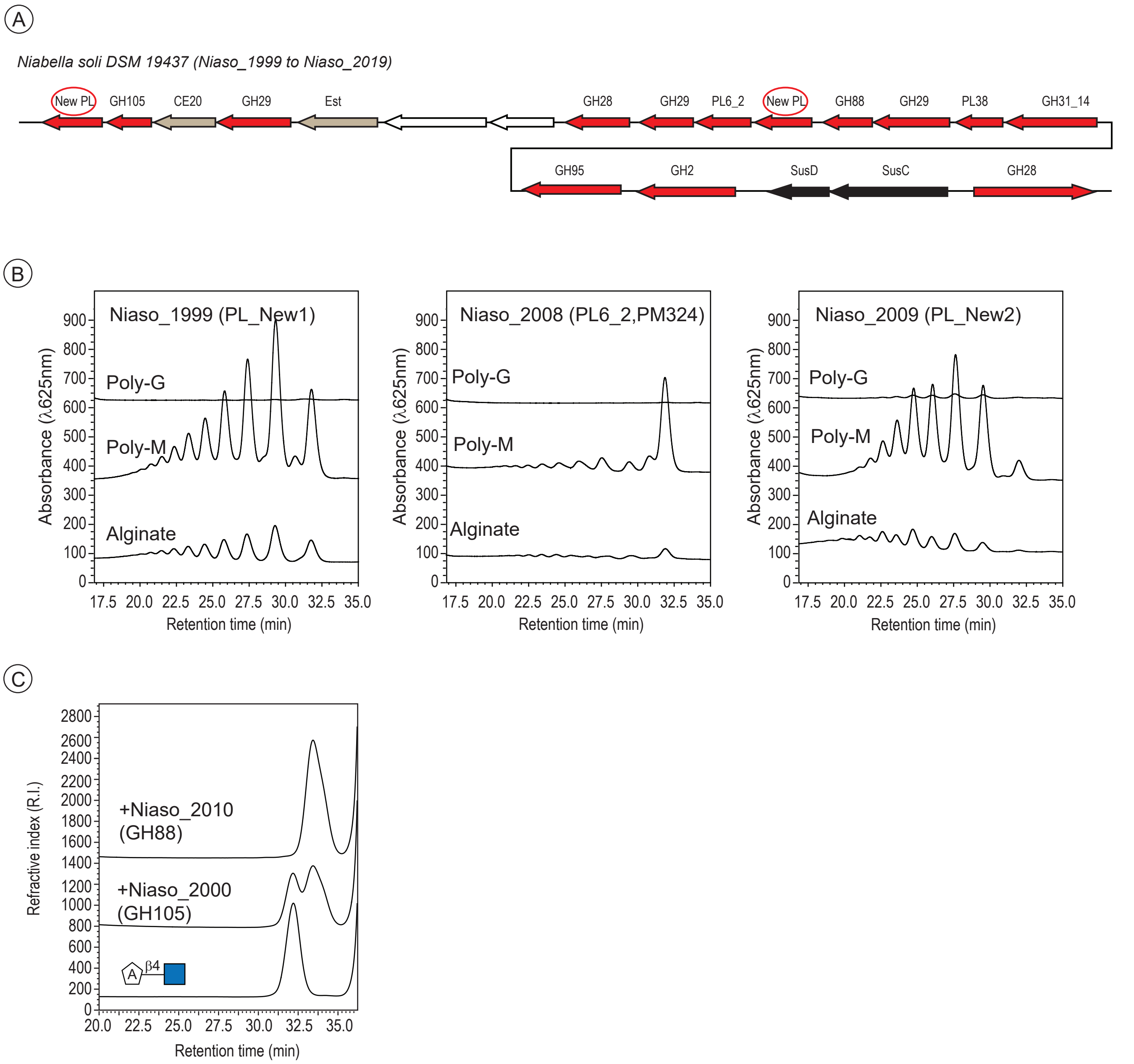


**Fig. S10:** Putative alginate lyase utilization loci. **A**) Genomic organization of the PUL encopassing three alginate lyases. (**B**) Size exclusion chromatograms of poly-mannuronic, poly-mannuronic/guluronic and poly-guluronic incubated with the first members of to new polysaccharide lyases families (Pl_New1 and PL_New2). The chromatograms demonstrated the poly-mannuronic specificities of these alginate lyases. (**C**) Size exclusion chromatograms of oligo-heparosan (DP2) incubated with Niaso_2000 (GH105) and Niaso_2010 (GH88) demonstrating the β-glcuronyl hydrolase activity of these enzymes.
