## Supplementary material for "Screening for Polysaccharide Utilization Loci Targeting Marine Polysaccharides": Figure_S11

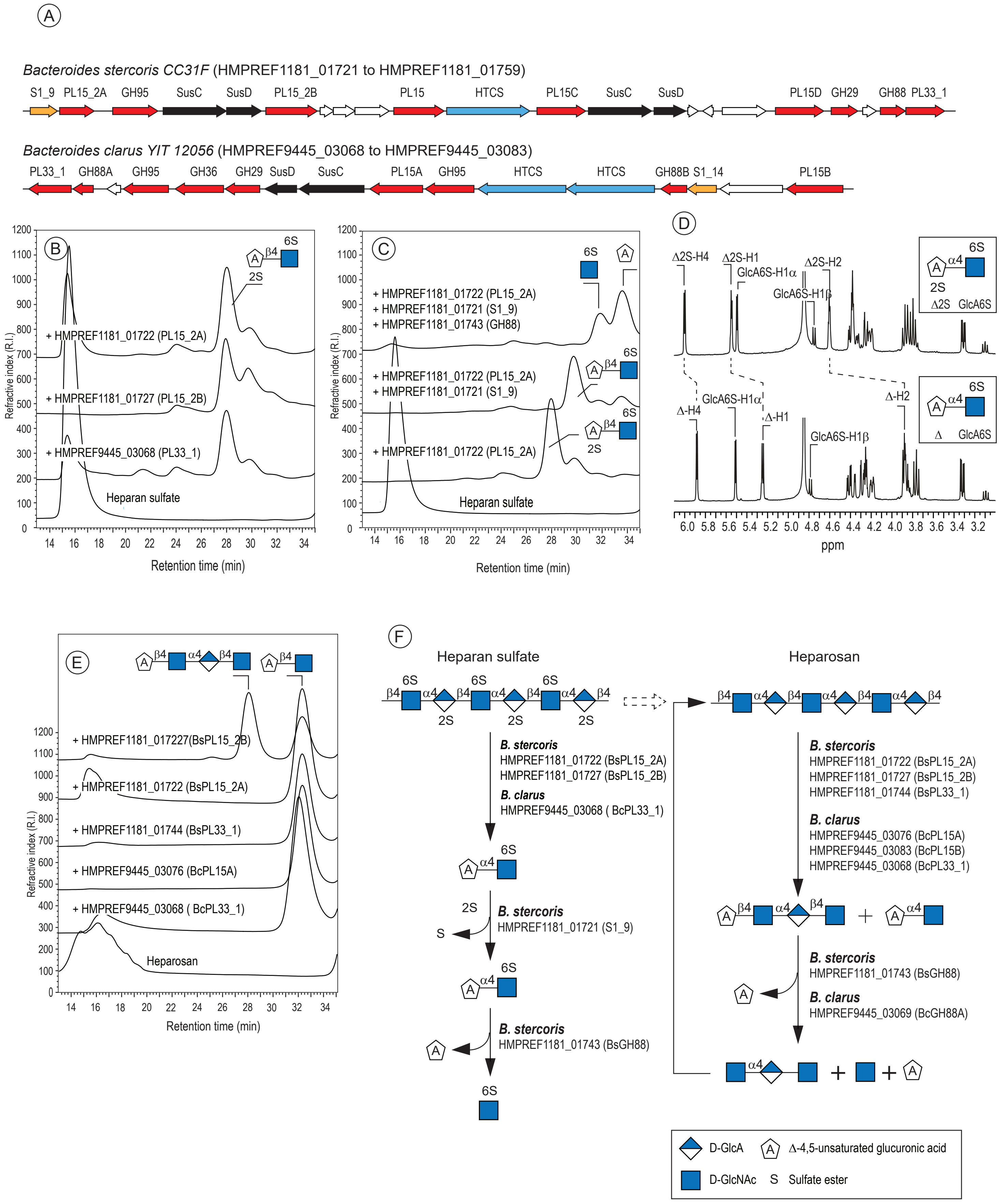


**Figure S11:** PUL involved in the degradation of heparan sulfate and heparosan. **A**) Organization of the PULs of *Bacteroides stercoris* CC31F and *Bacteroides clarus* YIT 12056. **B**) Chromatograms of the degradation products of heparan sulfate incubated with PL15_2 and PL33 heparan lyases. **C**) Observation of the multi-steps degradation of heparan sulfate. **D**) $^{1}$H NMR of the major heparin disaccharide incubated the S1_9 sulfatase demonstrating the removal of the sulfate at the position 2 of the unsaturated non-reducing end residue. **E**) Chromatogram of the degradation products of heparosan incubated with PL15 and PL33. Several enzymes are both heparan sulfate/heparosan lyases. **F**) Degradation pathway of heparan sulfate and heparosan deduced from the studied eznymes of the PUL.
