## Supplementary material for "Screening for Polysaccharide Utilization Loci Targeting Marine Polysaccharides": Figure_S12

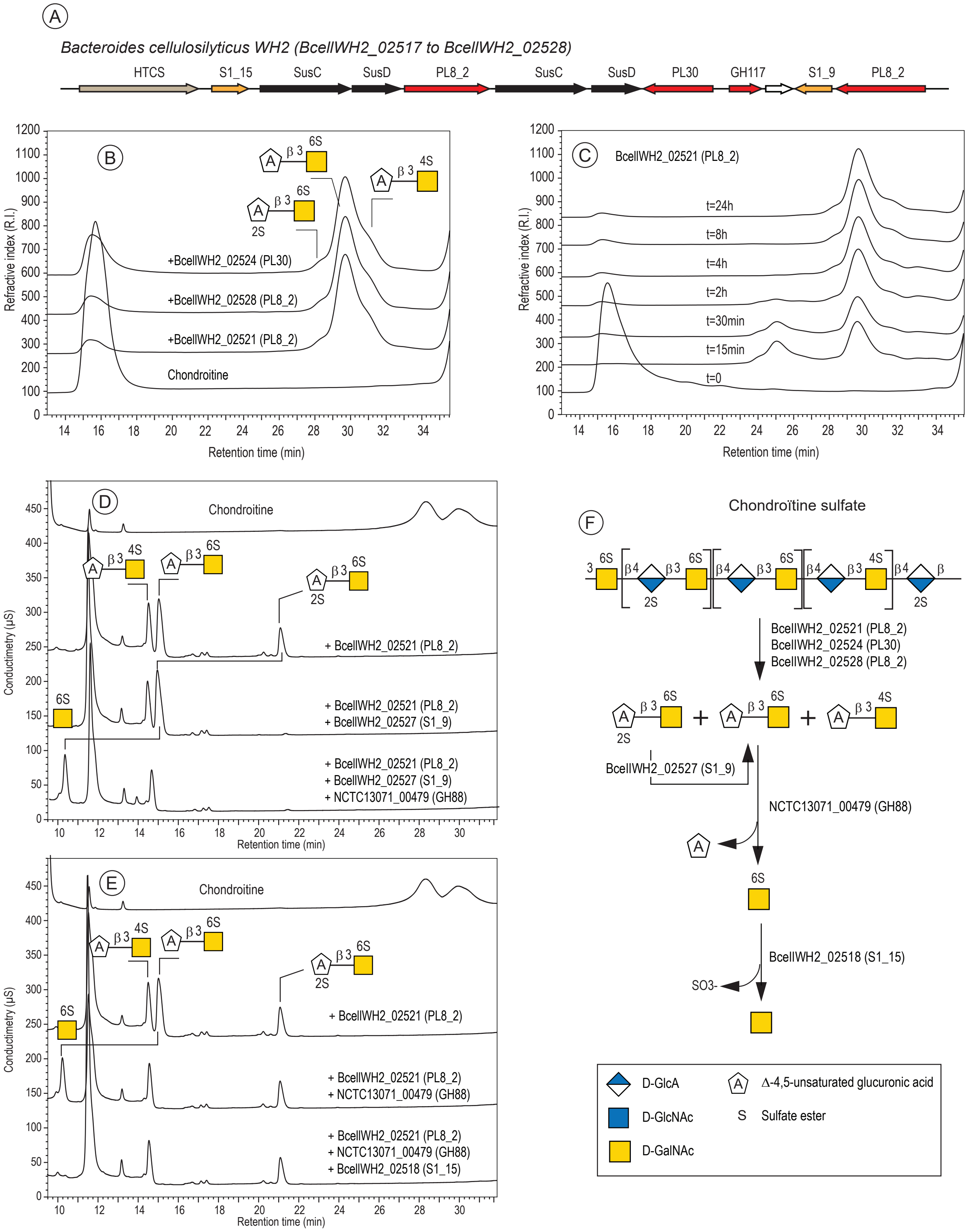


**Fig. S12:** Analysis of the PUL of *Bacteroides cellulosilyticus* WH2 involved in the degradation of the chondroitin sulfate. **A**) Organization of the PUL. **B**) Size exclusion chromatograms of the tree PL of the PUL showing similar end-products. **C**) Degradation kinetic of the PL8_2 (BcellWH2_02521). **D**) and **E**) Sequential degradation of chondroitin involving PL, GH and sulfatses. **F**) Degradation pathway of the chondroitin deduced from enzymology experiments.
