## Supplementary Table 1 for "Screening for Polysaccharide Utilization Loci Targeting Marine Polysaccharides"

**Table S1:** List of the selected PULs. The selected proteins listed glycoside hydrolases (GH), polysaccharides lyases (PL), sulfatases (S) and unclassified proteins (unk). * Not studied

| Eco | Strain | Locus (linked to PULDB) | Genbank | CAZy - SulfAtlas | Mw (kDa) |
|---|---|---|---|---|---|
| **Selected PUL 1: *Barnesiella viscericola DSM 18177* (Predicted PUL4)** | | | | | |
| **GH16 ▶ GH50 ▶ ◀ GH2 ◀ Sulf_1 ◀ GH20 ◀ Sulf_1 ◀ SusD ◀ SusC ◀ GH ◀ unk ◀ HTCS** | | | | | |
| Chicken gut | *Barnesiella viscericola DSM 18177* | Barvi_0589 | AHF11964.1 | GH16 | 48,2 |
| | *Barnesiella viscericola DSM 18177* | Barvi_0590 | AHF13634.1 | GH50 | 73,5 |
| | *Barnesiella viscericola DSM 18177* | Barvi_0592 | AHF11965.1 | GH2 | 116,7 |
| | *Barnesiella viscericola DSM 18177* | Barvi_0593 | AHF11966.1 | **S1_15** | 56,6 |
| | *Barnesiella viscericola DSM 18177* | Barvi_0594 | AHF11967.1 | GH20 | 86,6 |
| | *Barnesiella viscericola DSM 18177* | Barvi_0595 | AHF11968.1 | **S1_11** | 61,5 |
| | *Barnesiella viscericola DSM 18177* | Barvi_0598 | AHF11971.1 | unk | 94,7 |
| | *Barnesiella viscericola DSM 18177* | Barvi_0599 | AHF13635.1 | unk | 35,3 |
| **Selected PUL 2: *Polaribacter sejongensis KCTC 23670* (Predicted PUL 34)** | | | | | |
| **SusC ▶ SusD ▶ unk ▶ ◀ GH16 ◀ GH2 ◀ GH16 ◀ GH50 ◀ GH43 ◀ MFS ◀ GH117 ◀ EPI ◀ unk ◀ unk ◀ unk ◀ unk unk ▶ ◀ GH3 ◀ unk SusC ▶ SusD ▶ unk ▶** | | | | | |
| Marine | *Polaribacter sejongensis KCTC 23670* | BTO15_18110 | AUC23891.1 | GH16_16 | 45,8 |
| | *Polaribacter sejongensis KCTC 23670* | BTO15_18115 | AUC23892.1 | GH2 | 97,1 |
| | *Polaribacter sejongensis KCTC 23670* | BTO15_18120 | AUC23893.1 | GH16_15 | 46,8 |
| | *Polaribacter sejongensis KCTC 23670* | BTO15_18125 | AUC24144.1 | GH50 | 85,6 |
| | *Polaribacter sejongensis KCTC 23670* | BTO15_18130 | AUC23894.1 | GH43 | 52,6 |
| | *Polaribacter sejongensis KCTC 23670* | BTO15_18140 | AUC23895.1 | GH117 | 44,7 |
| | *Polaribacter sejongensis KCTC 23670* | BTO15_18175 | AUC23902.1 | GH3 | 81,3 |
| **Selected PUL 3: *Flavobacterium johnsoniae UW101* (Predicted PUL 12)** | | | | | |
| **HTCS ▶ SusC ▶ SusD ▶ GH10 ▶ unk ▶ GH50 ▶ GH2 ▶** | | | | | |
| Soil | *Flavobacterium johnsoniae UW101* | Fjoh_2079 | ABQ05109.1 | GH10 | 41,3 |
| | *Flavobacterium johnsoniae UW101* | Fjoh_2080 | ABQ05110.1 | unk | 32,8 |
| | *Flavobacterium johnsoniae UW101* | Fjoh_2081 | ABQ05111.1 | GH50 | 55,3 |
| | *Flavobacterium johnsoniae UW101* | Fjoh_2082 | ABQ05112.1 | GH2 | 69,8 |
| **Selected PUL 4: *Bacteroides ovatus ATCC 8483* (Pedicted PUL 39)** | | | | | |
| **◀ GH50 ◀ GH50 ◀ GH2\|CBM57 ◀ unk ◀ SusD ◀ SusC ◀ HTCS** | | | | | |
| H. gut | *Bacteroides ovatus ATCC 8483* | Bovatus_02307 | ALJ46940.1 | GH50 | 57,8 |
| | *Bacteroides ovatus ATCC 8483* | Bovatus_02308 | ALJ46941.1 | GH50 | 53,7 |
| | *Bacteroides ovatus ATCC 8483* | Bovatus_02309 | ALJ46942.1 | GH2\|CBM57 | 99,9 |
| **Selected PUL 5: *Wenyingzhuangia fucanilytica CZ1127* (Predicted PUL 14)** | | | | | |
| **◀ Sulf_1 ◀ CE6 ◀ unk ◀ GH50 ◀ Sulf_1 ◀ HTCS ◀ GH29 ◀ GH29 ◀ GH92 ◀ SusD ◀ SusC unk ▶ unk ▶ GH78** | | | | | |
| Marine | *Wenyingzhuangia fucanilytica CZ1127* | AXE80_10375 | ANW96654.1 | **S1_9*** | 49,0 |
| | *Wenyingzhuangia fucanilytica CZ1127* | AXE80_10380 | ANW96655.1 | CE6 | 26,2 |
| | *Wenyingzhuangia fucanilytica CZ1127* | AXE80_10385 | ANW97502.1 | unk | 32,8 |
| | *Wenyingzhuangia fucanilytica CZ1127* | AXE80_10390 | ANW96656.1 | GH50 | 54,8 |
| | *Wenyingzhuangia fucanilytica CZ1127* | AXE80_10395 | ANW97503.1 | **S1_23*** | 48,8 |
| | *Wenyingzhuangia fucanilytica CZ1127* | AXE80_10405 | ANW96658.1 | GH29 | 50,6 |
| | *Wenyingzhuangia fucanilytica CZ1127* | AXE80_10410 | ANW96659.1 | GH29 | 50,1 |
| | *Wenyingzhuangia fucanilytica CZ1127* | AXE80_10415 | ANW96660.1 | GH92 | 87,4 |
| | *Wenyingzhuangia fucanilytica CZ1127* | AXE80_10430 | ANW96663.1 | unk | 51,5 |
| | *Wenyingzhuangia fucanilytica CZ1127* | AXE80_10435 | ANW96664.1 | unk | 49,9 |
| | *Wenyingzhuangia fucanilytica CZ1127* | AXE80_10440 | ANW96665.1 | GH78 | 69,5 |
| **Selected PUL 6: *Saccharicrinis fermentans DSM 9555* (Predicted PUL 37)** | | | | | |
| **◀ GH2\|CBM57 ◀ GH136 ◀ CE6 ◀ unk ◀ SusD ◀ SusC ◀ SusD ◀ SusC ◀ Sulf_1 ◀ Sulf_1 ◀ GH50 ◀ Sulf_1 ◀ GH88 HTCS ▶ ◀ unk HTCS ▶ GH43_24 ▶ Sulf_1\|GH16 ▶ Sulf_1 ▶ GH2 ▶** | | | | | |
| Marine | *Saccharicrinis fermentans DSM 9555* | CytfeDRAFT_4602 | GAF03328.1 | **S1_7 *** | 61,3 |
| | *Saccharicrinis fermentans DSM 9555* | CytfeDRAFT_4603 | WP_052343392.1 | **S1_7*** | 60,3 |
| | *Saccharicrinis fermentans DSM 9555* | CytfeDRAFT_4604 | GAF03325.1 | GH50 | 53,7 |
| | *Saccharicrinis fermentans DSM 9555* | CytfeDRAFT_4605 | GAF03324.1 | **S1_23** | 53,4 |
| | *Saccharicrinis fermentans DSM 9555* | CytfeDRAFT_4606 | GAF03323.1 | GH88 | 46,4 |
| | *Saccharicrinis fermentans DSM 9555* | CytfeDRAFT_4610 | GAF03317.1 | GH43_24 | 43,5 |
| | *Saccharicrinis fermentans DSM 9555* | CytfeDRAFT_4611 | WP_081736078.1 | **S1_19\|GH16_** | 84,8 |
| | *Saccharicrinis fermentans DSM 9555* | CytfeDRAFT_4612 | GAF03314.1 | **S1_19*** | 57,1 |
| | *Saccharicrinis fermentans DSM 9555* | CytfeDRAFT_4613 | WP_081736079.1 | GH2 | 119,7 |
| **Selected PUL 7: *Aquimarina agarivorans* (Cazymes cluster 1)** | | | | | |
| **◀ unk ◀ unk ◀ Sulf_4 GH16 ▶ ◀ GH50 ◀ unk** | | | | | |
| ne | *Aquimarina agarivorans* | FbacHQ_010100004671 | WP_010521076.1 | GH50 | 68,4 |

| | | | | | |
|---|---|---|---|---|---|
| Mari | *Aquimarina agarivorans* | FbacHQ_010100004666 | WP_010521075.1 | GH16_26\|CBI | 58,0 |
| | *Aquimarina agarivorans* | FbacHQ_010100004661 | WP_010521074.1 | **Sulf_4** | 38,4 |
| **Selected PUL 8:** ***Bacteroides cellulosilyticus WH2*** **(Predicted PUL 45)** | | | | | |
| **◄ GH16\|Sulf_1 ◄ GH29 ◄ GH2 ◄ GH20 ◄ Sulf_1 ◄ SusD ◄ SusC HTCS ► ◄ GH159 ◄ unk ◄ GH146 ◄ GH50 ◄ GH159 ◄ unk ◄ SusD ◄ SusC ◄ SusD ◄ SusC HTCS ► ◄ Sulf_1 ◄ GH2** | | | | | |
| Human gut | *Bacteroides cellulosilyticus WH2* | BcellWH2_02348 | ALJ59588.1 | **S1_16** | 91,0 |
| | *Bacteroides cellulosilyticus WH2* | BcellWH2_02349 | ALJ59589.1 | GH29 | 70,1 |
| | *Bacteroides cellulosilyticus WH2* | BcellWH2_02350 | ALJ59590.1 | GH2 | 116,5 |
| | *Bacteroides cellulosilyticus WH2* | BcellWH2_02351 | ALJ59591.1 | GH20 | 87,3 |
| | *Bacteroides cellulosilyticus WH2* | BcellWH2_02352 | ALJ59592.1 | **S1_11** | 61,5 |
| | *Bacteroides cellulosilyticus WH2* | BcellWH2_02356 | ALJ59596.1 | GH159 | 36,2 |
| | *Bacteroides cellulosilyticus WH2* | BcellWH2_02357 | ALJ59597.1 | unk | 44,2 |
| | *Bacteroides cellulosilyticus WH2* | BcellWH2_02358 | ALJ59598.1 | GH146 | 88,2 |
| | *Bacteroides cellulosilyticus WH2* | BcellWH2_02359 | ALJ59599.1 | GH50 | 76,6 |
| | *Bacteroides cellulosilyticus WH2* | BcellWH2_02360 | ALJ59600.1 | GH159 | 39,8 |
| | *Bacteroides cellulosilyticus WH2* | BcellWH2_02361 | ALJ59601.1 | CE20 | 71,5 |
| | *Bacteroides cellulosilyticus WH2* | BcellWH2_02367 | ALJ59607.1 | **S1_20** | 57,6 |
| | *Bacteroides cellulosilyticus WH2* | BcellWH2_02368 | ALJ59608.1 | GH2 | 101,1 |
| **Selected PUL 9:** ***Bacteroides xylanisolvens NLAE-zl-H194*** **(Predited PUL 60)** | | | | | |
| **◄ GH105 ◄ GH154 ◄ GH154\|Sulf_1 ◄ GH43_28 ◄ Sulf_1 ◄ GH88\|GH154 ◄ GH50 ◄ unk ◄ SusD ◄ SusC ◄** | | | | | |
| Human gut | *Bacteroides xylanisolvens NLAE-zl-H194* | H194DRAFT_01679 | WP_008641054.1 | GH105 | 42,7 |
| | *Bacteroides xylanisolvens NLAE-zl-H194* | H194DRAFT_01680 | WP_008641056.1 | GH154 | 46,0 |
| | *Bacteroides xylanisolvens NLAE-zl-H194* | H194DRAFT_01681 | WP_074706425.1 | **S1_14\|GH154** | 96,2 |
| | *Bacteroides xylanisolvens NLAE-zl-H194* | H194DRAFT_01682 | WP_195564478.1 | GH43_28 | 69,0 |
| | *Bacteroides xylanisolvens NLAE-zl-H194* | H194DRAFT_01683 | WP_195564479.1 | **S1_9*** | 51,8 |
| | *Bacteroides xylanisolvens NLAE-zl-H194* | H194DRAFT_01684 | WP_032850949.1 | GH88\|GH154 | 90,1 |
| | *Bacteroides xylanisolvens NLAE-zl-H194* | H194DRAFT_01685 | WP_004313499.1 | GH50 | 58,3 |
| | *Bacteroides xylanisolvens NLAE-zl-H194* | H194DRAFT_01686 | WP_195564480.1 | unk | 66,3 |
| | *Bacteroides xylanisolvens NLAE-zl-H194* | H194DRAFT_01689 | WP_195564481.1 | CBM\|CBM\|CE | 114,9 |
| | *Bacteroides xylanisolvens NLAE-zl-H194* | H194DRAFT_01691 | WP_195564483.1 | GH2 | 109,0 |
| **Selected PUL 10:** ***Bacteroides ovatus ATCC 8483*** **(Predicted PUL 81)** | | | | | |
| **HTCS ► SusC ► SusD ► unk ► unk ► GH50 ► unk ► GH ► GH31 ► GH2\|CBM57 ► CE6 ►** | | | | | |
| Human gut | *Bacteroides ovatus ATCC 8483* | Bovatus_03901 | ALJ48504.1 | CE6 | 29,9 |
| | *Bacteroides ovatus ATCC 8483* | Bovatus_03900 | ALJ48503.1 | GH2\|CBM | 102,6 |
| | *Bacteroides ovatus ATCC 8483* | Bovatus_03899 | ALJ48502.1 | GH31_4 | 90,3 |
| | *Bacteroides ovatus ATCC 8483* | Bovatus_03898 | ALJ48501.1 | unk | 69,1 |
| | *Bacteroides ovatus ATCC 8483* | Bovatus_03897 | ALJ48500.1 | unk | 38,0 |
| | *Bacteroides ovatus ATCC 8483* | Bovatus_03896 | ALJ48499.1 | GH50 | 42,0 |
| | *Bacteroides ovatus ATCC 8483* | Bovatus_03895 | ALJ48498.1 | unk | 56,2 |
| **Selected PUL 11: Bacteroides thetaiotaomicron 7330 (Predicted PUL 29)** | | | | | |
| **◄ HTCS unk ► unk ► GH18 ► SusC ► SusD ► GH ► Sulf_1 ► GH20 ► GH2 ► GH29 ► Sulf_1 ► unk ►** | | | | | |
| Human gut | *Bacteroides thetaiotaomicron 7330* | Btheta7330_01428 | ALJ40996.1 | GH18 | 60,4 |
| | *Bacteroides thetaiotaomicron 7330* | Btheta7330_01431 | ALJ40999.1 | unk | 65,5 |
| | *Bacteroides thetaiotaomicron 7330* | Btheta7330_01432 | ALJ41000.1 | **S1_11** | 61,8 |
| | *Bacteroides thetaiotaomicron 7330* | Btheta7330_01433 | ALJ41001.1 | GH20 | 87,8 |
| | *Bacteroides thetaiotaomicron 7330* | Btheta7330_01434 | ALJ41002.1 | GH2 | 116,1 |
| | *Bacteroides thetaiotaomicron 7330* | Btheta7330_01435 | ALJ41003.1 | GH29 | 68,9 |
| | *Bacteroides thetaiotaomicron 7330* | Btheta7330_01436 | ALJ41004.1 | **S1_15** | 56,4 |
| **Selected PUL 12:** ***Parabacteroides gordonii DSM 23371*** **(Predicted PUL 51)** | | | | | |
| **AraC ► ◄ unk ◄ Sulf_1 ◄ GH29 ◄ Sulf_1 ◄ unk ◄ GH117\|GH117 ◄ SusD ◄ SusC ◄ Anti-σ ECF-σ ►** | | | | | |
| Human gut | *Parabacteroides gordonii DSM 23371* | F592DRAFT_02147 | KKB57897.1 | Esterase | 32,3 |
| | *Parabacteroides gordonii DSM 23371* | F592DRAFT_02148 | KKB57898.1 | **S1_16*** | 59,1 |
| | *Parabacteroides gordonii DSM 23371* | F592DRAFT_02149 | KKB57899.1 | GH29 | 52,6 |
| | *Parabacteroides gordonii DSM 23371* | F592DRAFT_02150 | KKB57900.1 | **S1_16*** | 55,8 |
| | *Parabacteroides gordonii DSM 23371* | F592DRAFT_02152 | KKB57902.1 | GH117\|GH11 | 76,6 |
| | *Parabacteroides gordonii DSM 23371* | F592DRAFT_02157 | KKB57907.1 | **S1_7*** | 54,9 |
| | *Parabacteroides gordonii DSM 23371* | F592DRAFT_02158 | KKB57908.1 | **S1_15*** | 57,3 |
| | *Parabacteroides gordonii DSM 23371* | F592DRAFT_02159 | KKB57909.1 | **S1_24*** | 49,0 |
| | *Parabacteroides gordonii DSM 23371* | F592DRAFT_02160 | KKB57910.1 | **S1_11*** | 60,8 |
| | *Parabacteroides gordonii DSM 23371* | F592DRAFT_02161 | KKB57911.1 | **S1_14*** | 52,2 |
| | *Parabacteroides gordonii DSM 23371* | F592DRAFT_02162 | KKB57912.1 | **S1_15*** | 55,6 |
| | *Parabacteroides gordonii DSM 23371* | F592DRAFT_02163 | KKB57913.1 | GH20\|CBM | 85,9 |
| | *Parabacteroides gordonii DSM 23371* | F592DRAFT_02164 | KKB57914.1 | GH117 | 45,5 |
| | *Parabacteroides gordonii DSM 23371* | F592DRAFT_02165 | KKB57915.1 | **S1_23*** | 50,8 |
| **Selected PUL 13:** ***Algoriphagus chordae*** **DSM 19830 (Predicted PUL 34)** | | | | | |

| | | | | | |
|---|---|---|---|---|---|
| **◄ Sulf_1 ◄ Sulf_1 unk ► Sulf_1 ► unk ► Sulf_1 ► GH117 ► unk ► GH117 ► SusC ► SusD ► unk ► Sulf_1 ► ◄ GH95** | | | | | |
| Marine | *Algoriphagus chordae DSM 19830* | LV85DRAFT_03910 | PZX47561.1 | **S1_11*** | 65,3 |
| | *Algoriphagus chordae DSM 19830* | LV85DRAFT_03911 | PZX47562.1 | **S1_11*** | 61,5 |
| | *Algoriphagus chordae DSM 19830* | LV85DRAFT_03913 | PZX47564.1 | **S1_15*** | 57,3 |
| | *Algoriphagus chordae DSM 19830* | LV85DRAFT_03915 | PZX47566.1 | **S1_7*** | 53,4 |
| | *Algoriphagus chordae DSM 19830* | LV85DRAFT_03916 | PZX47567.1 | GH117 | 51,2 |
| | *Algoriphagus chordae DSM 19830* | LV85DRAFT_03917 | PZX47568.1 | unk | 48,7 |
| | *Algoriphagus chordae DSM 19830* | LV85DRAFT_03918 | PZX47569.1 | GH117 | 40,8 |
| | *Algoriphagus chordae DSM 19830* | LV85DRAFT_03922 | PZX47573.1 | **S1_28*** | 55,5 |
| | *Algoriphagus chordae DSM 19830* | LV85DRAFT_03923 | PZX47574.1 | GH95 | 98,9 |
| **Selected PUL 14: *Bacteroides dorei DSM 17855* (Cazymes cluster 3)** | | | | | |
| **GH29 ► unk ► unk ► Sulf_1 ► GH29 ► Sulf_1 ► GH20 ► GH117 ►** | | | | | |
| Human gut | *Phocaeicola dorei DSM 17855* | BACDOR_01585 | EEB26054.1 | **S1_46** | 57,5 |
| | *Phocaeicola dorei DSM 17855* | BACDOR_01583 | EEB26052.1 | **S1_11*** | 60,1 |
| | *Phocaeicola dorei DSM 17855* | BACDOR_01584 | EEB26053.1 | GH29 | 77,7 |
| | *Phocaeicola dorei DSM 17855* | BACDOR_01587 | EEB26056.1 | GH117 | 45,9 |
| | *Phocaeicola dorei DSM 17855* | BACDOR_01586 | EEB26055.1 | GH20 | 62,1 |
| | *Phocaeicola dorei DSM 17855* | BACDOR_01580 | EEB26049.1 | GH29 | 37,6 |
| | *Phocaeicola dorei DSM 17855* | BACDOR_01574 | EEB26043.1 | unk | 96,4 |
| **Selected PUL 15: *Chitinophaga pinensis DSM 2588* (Predicted PUL 70)** | | | | | |
| **SusC ► SusD ► ◄ GH20 ◄ unk GH127 ► Sulf_4 ► unk ►** | | | | | |
| Soil | *Chitinophaga pinensis DSM 2588* | Cpin_4994 | ACU62427.1 | GH20 | 85,6 |
| | *Chitinophaga pinensis DSM 2588* | Cpin_4996 | ACU62429.1 | GH127 | 90,3 |
| | *Chitinophaga pinensis DSM 2588* | Cpin_4997 | ACU62430.1 | **S4*** | 30,5 |
| **Selected PUL 16: *Bacteroides cellulosilyticus WH2* (Predicted PUL 43)** | | | | | |
| **HTCS ► Sulf_1 ► SusC ► SusD ► PL8_2 ► SusC ► SusD ► ◄ PL30 GH117 ► unk ► ◄ Sulf_1 ◄ PL8_2 ◄ unk** | | | | | |
| Human gut | *Bacteroides cellulosilyticus WH2* | BcellWH2_02518 | ALJ59757.1 | **S1_15** | 0,0 |
| | *Bacteroides cellulosilyticus WH2* | BcellWH2_02521 | ALJ59760.1 | PL8_2 | 110,7 |
| | *Bacteroides cellulosilyticus WH2* | BcellWH2_02524 | ALJ59763.1 | PL30 | 92,5 |
| | *Bacteroides cellulosilyticus WH2* | BcellWH2_02525 | ALJ59764.1 | GH117 | 42,3 |
| | *Bacteroides cellulosilyticus WH2* | BcellWH2_02527 | ALJ59766.1 | **S1_9** | 51,8 |
| | *Bacteroides cellulosilyticus WH2* | BcellWH2_02528 | ALJ59767.1 | PL8_2 | 116,2 |
| | *Bacteroides cellulosilyticus WH2* | BcellWH2_02529 | ALJ59768.1 | unk | 26,5 |
| **Selected PUL 17: *Arenibacter echinorum DSM 23522* (Cazymes cluster 4)** | | | | | |
| **unk ► unk ► GH ► GH33 ► GH ► unk ► unk ► ◄ unk ◄ unk ◄ unk ◄ MFS ◄ unk ◄ GH29\|GH117 ◄ Sulf_1 ◄** | | | | | |
| Marine | *Arenibacter echinorum DSM 23522* | LV92DRAFT_02561 | RAJ11630.1 | GH117\|GH29 | 90,7 |
| | *Arenibacter echinorum DSM 23522* | LV92DRAFT_02562 | RAJ11631.1 | **S1_16*** | 57,7 |
| | *Arenibacter echinorum DSM 23522* | LV92DRAFT_02563 | RAJ11632.1 | GH29 | 51,9 |
| | *Arenibacter echinorum DSM 23522* | LV92DRAFT_02564 | RAJ11633.1 | CE20 | 52,6 |
| | *Arenibacter echinorum DSM 23522* | LV92DRAFT_02565 | RAJ11634.1 | Esterase | 78,7 |
| | *Arenibacter echinorum DSM 23522* | LV92DRAFT_02566 | RAJ11635.1 | GH31_14 | 82,0 |
| | *Arenibacter echinorum DSM 23522* | LV92DRAFT_02568 | RAJ11637.1 | **S1_15*** | 59,4 |
| | *Arenibacter echinorum DSM 23522* | LV92DRAFT_02569 | RAJ11638.1 | unk | 87,9 |
| | *Arenibacter echinorum DSM 23522* | LV92DRAFT_02570 | RAJ11639.1 | GH29 | 84,3 |
| **Selected PUL 18: *Polaribacter reichenbachii KCTC 23969* (Cazymes cluster 5)** | | | | | |
| **◄ GntR ◄ Sulf_1 ◄ Sulf_1 ◄ GH117 ◄ GH16 ◄ GH117 ◄ GH2 ◄ GH2 ◄ unk ◄ unk unk ► ◄ GH117 ◄ Sulf_1 ◄ Sulf_1 ◄ GH28 ◄ Sulf_1 ◄ Sulf_1 ◄ GH2 ◄ Sulf_1 ◄ GH16 ◄ GH117 ◄ GH29 ◄ GH117 ◄ GH2 ◄ GH2 unk ►** | | | | | |
| Marine | *Polaribacter reichenbachii KCTC 23969* | BTO17_12240 | AUC19412.1 | GH117 | 48,6 |
| | *Polaribacter reichenbachii KCTC 23969* | BTO17_12245 | AUC19413.1 | GH16 | 48,9 |
| | *Polaribacter reichenbachii KCTC 23969* | BTO17_12250 | AUC19414.1 | GH117 | 42,5 |
| | *Polaribacter reichenbachii KCTC 23969* | BTO17_12255 | AUC19415.1 | GH2 | 120,7 |
| | *Polaribacter reichenbachii KCTC 23969* | BTO17_12260 | AUC19416.1 | GH2 | 66,2 |
| | *Polaribacter reichenbachii KCTC 23969* | BTO17_12280 | AUC19420.1 | GH117 | 48,6 |
| | *Polaribacter reichenbachii KCTC 23969* | BTO17_12285 | AUC19421.1 | **S1_28** | 49,3 |
| | *Polaribacter reichenbachii KCTC 23969* | BTO17_12290 | AUC19422.1 | **S1_28** | 59,4 |
| | *Polaribacter reichenbachii KCTC 23969* | BTO17_12295 | AUC19423.1 | GH28 | 52,1 |
| | *Polaribacter reichenbachii KCTC 23969* | BTO17_12300 | AUC19424.1 | **S1_25** | 62,7 |
| | *Polaribacter reichenbachii KCTC 23969* | BTO17_12305 | AUC19425.1 | **S1_29** | 52,7 |
| | *Polaribacter reichenbachii KCTC 23969* | BTO17_12310 | AUC20527.1 | GH2 | 93,0 |
| | *Polaribacter reichenbachii KCTC 23969* | BTO17_12315 | AUC19426.1 | **S1_15** | 57,1 |
| | *Polaribacter reichenbachii KCTC 23969* | BTO17_12320 | AUC19427.1 | GH16_11 | 32,8 |
| | *Polaribacter reichenbachii KCTC 23969* | BTO17_12325 | AUC19428.1 | GH117 | 48,4 |
| | *Polaribacter reichenbachii KCTC 23969* | BTO17_12330 | AUC19429.1 | GH29 | 58,4 |
| | *Polaribacter reichenbachii KCTC 23969* | BTO17_12335 | AUC19430.1 | GH117 | 49,6 |
| | *Polaribacter reichenbachii KCTC 23969* | BTO17_12340 | AUC19431.1 | GH2 | 108,5 |

| | | | | | |
|---|---|---|---|---|---|
| | *Polaribacter reichenbachii KCTC 23969* | BTO17_12345 | AUC19432.1 | GH2 | 108,5 |
| | *Polaribacter reichenbachii KCTC 23969* | BTO17_12355 | AUC19434.1 | GH28 | 52,5 |
| | *Polaribacter reichenbachii KCTC 23969* | BTO17_12360 | AUC19435.1 | unk | 91,6 |
| | *Polaribacter reichenbachii KCTC 23969* | BTO17_12365 | AUC19436.1 | **S1_8** | 71,0 |
| | *Polaribacter reichenbachii KCTC 23969* | BTO17_12370 | AUC19437.1 | GH29 | 58,9 |
| **Selected PUL 19: *Algibacter lectus JCM 19300* (Predicted PUL 23)** | | | | | |
| **◄ SusD ◄ unk ◄ SusC Sulf_1 ► unk ► ◄ GH117 ◄ GH95 ◄ unk ◄ GH117 ◄ GH117 ◄ GH2 ◄ GH117 ◄ unk ◄ GH97 ◄ Sulf_1 ◄ CBM47 ◄ Sulf_1 ◄ Sulf_1 SusC ► SusD ► SusC ► SusD ► unk ►** | | | | | |
| Marine | *Algibacter lectus JCM 19300* | Ga0062136_116103 | GAL64199.1 | **S1_16*** | 54,0 |
| | *Algibacter lectus JCM 19300* | Ga0062136_116104 | GAL64200.1 | unk | 36,0 |
| | *Algibacter lectus JCM 19300* | Ga0062136_116105 | GAL64201.1 | GH117 | 40,8 |
| | *Algibacter lectus JCM 19300* | Ga0062136_116106 | GAL64202.1 | GH95 | 86,8 |
| | *Algibacter lectus JCM 19300* | Ga0062136_116108 | GAL64204.1 | GH117 | 37,6 |
| | *Algibacter lectus JCM 19300* | Ga0062136_116109 | GAL64205.1 | GH117 | 39,2 |
| | *Algibacter lectus JCM 19300* | Ga0062136_116110 | GAL64206.1 | GH2 | 96,7 |
| | *Algibacter lectus JCM 19300* | Ga0062136_116111 | GAL64207.1 | GH117 | 36,6 |
| | *Algibacter lectus JCM 19300* | Ga0062136_116112 | GAL64208.1 | Esterase | 56,7 |
| | *Algibacter lectus JCM 19300* | Ga0062136_116113 | GAL64209.1 | GH97 | 71,4 |
| | *Algibacter lectus JCM 19300* | Ga0062136_116114 | GAL64210.1 | **S1_15*** | 59,8 |
| | *Algibacter lectus JCM 19300* | Ga0062136_116115 | GAL64211.1 | CBM47 | 57,6 |
| | *Algibacter lectus JCM 19300* | Ga0062136_116116 | GAL64212.1 | **S1_20*** | 54,0 |
| | *Algibacter lectus JCM 19300* | Ga0062136_116117 | GAL64213.1 | **S1_15*** | 54,0 |
| **Selected PUL 20: *Bacteroides ovatus NLAE-zl-C57* (Predicted PUL 19)** | | | | | |
| **◄ GH2 ◄ PL11_1 GH28 ► ◄ unk ◄ unk ◄ GH117 ◄ unk ◄ HTCS SusC ► SusD ► unk ► unk ► unk ► ◄ PL11_1 PL9_1 ► unk ►** | | | | | |
| cow gut | *Bacteroides ovatus NLAE-zl-C57* | Ga0104390_100622 | SDH45369.1 | GH2 | 112,4 |
| | *Bacteroides ovatus NLAE-zl-C57* | Ga0104390_100623 | SDH45386.1 | PL11_1 | 71,0 |
| | *Bacteroides ovatus NLAE-zl-C57* | Ga0104390_100624 | SDH45426.1 | GH28 | 54,0 |
| | *Bacteroides ovatus NLAE-zl-C57* | Ga0104390_100625 | SDH45462.1 | unk | 121,9 |
| | *Bacteroides ovatus NLAE-zl-C57* | Ga0104390_100627 | SDH45527.1 | GH117 | 41,5 |
| | *Bacteroides ovatus NLAE-zl-C57* | Ga0104390_100628 | SDH45570.1 | unk | 28,7 |
| | *Bacteroides ovatus NLAE-zl-C57* | Ga0104390_100632 | SDH45691.1 | unk | 122,1 |
| | *Bacteroides ovatus NLAE-zl-C57* | Ga0104390_100635 | SDH45794.1 | PL11_1 | 74,3 |
| | *Bacteroides ovatus NLAE-zl-C57* | Ga0104390_100636 | SDH45820.1 | PL9_1 | 56,3 |
| **Selected PUL 21: *Sphingobacterium thalpophilum DSM 11723* (Predicted PUL 13)** | | | | | |
| **CE12 ► unk ► SusC ► SusD ► unk ► unk ► SusC ► SusD ► GH51 ► GH115 ► GH43_18 ► unk ► GH2 ► unk ► ◄ GH28 GH28 ► PL11_1 ► CE12 ► CE12 ► Sulf_1 ► GH105 ► GH2 ► unk ► GH106 ► SusC ► SusD ►** | | | | | |
| H. wound | *Sphingobacterium thalpophilum DSM 1172[illegible]* | BS55DRAFT_01026 | VTR39317.1 | GH117 | 40,0 |
| | *Sphingobacterium thalpophilum DSM 1172[illegible]* | BS55DRAFT_01027 | VTR39324.1 | GH28 | 52,3 |
| | *Sphingobacterium thalpophilum DSM 1172[illegible]* | BS55DRAFT_01028 | VTR39331.1 | unk | 135,5 |
| | *Sphingobacterium thalpophilum DSM 1172[illegible]* | BS55DRAFT_01029 | VTR39338.1 | unk | 81,7 |
| **Selected PUL 22: *Maribacter forsetii DSM 18668* (Predicted PUL 13)** | | | | | |
| **◄ GH95 ◄ Sulf_1 ◄ PL40 ◄ unk ◄ GH117 ◄ PL33_2 ◄ unk ◄ GH29 ◄ GH ◄ Pept_SC ◄ PL40 ◄ Sulf_1 ◄ unk ◄ MFS ◄ GH88 SusC ► SusD ► ◄ HTCS** | | | | | |
| Marine | *Maribacter forsetii DSM 18668* | P177DRAFT_02785 | WP_036155605.1 | GH95 | 93,0 |
| | *Maribacter forsetii DSM 18668* | P177DRAFT_02786 | WP_084684762.1 | **S1_25*** | 59,2 |
| | *Maribacter forsetii DSM 18668* | P177DRAFT_02787 | WP_209435192.1 | PL40 | 104,4 |
| | *Maribacter forsetii DSM 18668* | P177DRAFT_02788 | WP_157486574.1 | PL43 | 50,9 |
| | *Maribacter forsetii DSM 18668* | P177DRAFT_02789 | WP_209435193.1 | GH117 | 41,1 |
| | *Maribacter forsetii DSM 18668* | P177DRAFT_02790 | WP_209435194.1 | PL33 | 72,5 |
| | *Maribacter forsetii DSM 18668* | P177DRAFT_02792 | WP_036155612.1 | GH29 | 51,2 |
| | *Maribacter forsetii DSM 18668* | P177DRAFT_02793 | WP_036155614.1 | unk | 88,7 |
| | *Maribacter forsetii DSM 18668* | P177DRAFT_02795 | WP_084684696.1 | PL40 | 106,5 |
| | *Maribacter forsetii DSM 18668* | P177DRAFT_02796 | WP_084684697.1 | **S1_11*** | 64,8 |
| | *Maribacter forsetii DSM 18668* | P177DRAFT_02797 | WP_036155618.1 | Esterase | 31,8 |
| | *Maribacter forsetii DSM 18668* | P177DRAFT_02799 | WP_036155622.1 | GH88 | 45,5 |
| **Selected PUL 23: *Saccharicrinis fermentans DSM 9555* (Cazymes cluster 6)** | | | | | |
| **◄ GH ◄ PL40 ◄ unk ◄ unk ◄ unk ◄ GH39 ◄ unk ◄ CBM6 ◄ GH117 ◄ unk ◄ Sulf_1 ◄ GH2 ◄ unk ◄ Sulf_1 ◄ unk ◄ unk ◄ GH97 ◄ Sulf_1 HTCS ► ◄ GH29** | | | | | |
| | *Saccharicrinis fermentans DSM 9555* | CytfeDRAFT_1518 | GAF01948.1 | unk | 89,5 |
| | *Saccharicrinis fermentans DSM 9555* | CytfeDRAFT_1519 | GAF01947.1 | PL40 | 101,0 |
| | *Saccharicrinis fermentans DSM 9555* | CytfeDRAFT_1520 | GAF01945.1 | Esterase | 31,0 |
| | *Saccharicrinis fermentans DSM 9555* | CytfeDRAFT_1521 | GAF01944.1 | Esterase | 26,5 |
| | *Saccharicrinis fermentans DSM 9555* | CytfeDRAFT_1522 | GAF01943.1 | unk | 101,6 |
| | *Saccharicrinis fermentans DSM 9555* | CytfeDRAFT_1523 | GAF01942.1 | GH39 | 63,5 |
| | *Saccharicrinis fermentans DSM 9555* | CytfeDRAFT_1524 | GAF01941.1 | PL43 | 51,1 |

| | | | | | |
|---|---|---|---|---|---|
| Marine | *Saccharicrinis fermentans DSM 9555* | CytfeDRAFT_1525 | GAF01940.1 | CBM6\|CBM96 | 109,4 |
| | *Saccharicrinis fermentans DSM 9555* | CytfeDRAFT_1526 | GAF01939.1 | GH117 | 37,2 |
| | *Saccharicrinis fermentans DSM 9555* | CytfeDRAFT_1527 | WP_044212102.1 | GH172 | 24,0 |
| | *Saccharicrinis fermentans DSM 9555* | CytfeDRAFT_1528 | WP_027471340.1 | **S1_20*** | 53,8 |
| | *Saccharicrinis fermentans DSM 9555* | CytfeDRAFT_1529 | GAF01935.1 | GH2 | 90,9 |
| | *Saccharicrinis fermentans DSM 9555* | CytfeDRAFT_1530 | WP_081735939.1 | CE20 | 57,2 |
| | *Saccharicrinis fermentans DSM 9555* | CytfeDRAFT_1531 | GAF01932.1 | **S1_14*** | 23,9 |
| | *Saccharicrinis fermentans DSM 9555* | CytfeDRAFT_1532 | GAF01931.1 | unk | 29,1 |
| | *Saccharicrinis fermentans DSM 9555* | CytfeDRAFT_1534 | GAF01930.1 | GH97 | 72,2 |
| | *Saccharicrinis fermentans DSM 9555* | CytfeDRAFT_1535 | GAF01929.1 | **S1_16*** | 53,9 |
| | *Saccharicrinis fermentans DSM 9555* | CytfeDRAFT_1537 | WP_027471346.1 (G | GH29 | 59,1 |
| **Selected PUL 24: *Flammeovirga sp. MY04* (Predicted PUL 37)** | | | | | |
| **◄ GH117 Sulf_1 ► Sulf_1 ► Sulf_1 ► GH2 ► Sulf_1 ► Sulf_1 ► GH136\|CBM16\|CBM16 ► ◄ GH95 ◄ Sulf_1 ◄ GH16 ◄ GH2 ◄ GH128 ◄ SusD ◄ SusC Sulf_1 ► HTCS ► GH136 ► unk ► ◄ GH136 ◄ GH136 ◄ Sulf_1 ◄ GH2 ◄ GH2 ◄ GH136 ◄ Sulf_1 ◄ SusD ◄ SusC** | | | | | |
| Marine | *Flammeovirga sp. MY04* | MY04_5109 | ANQ52444.1 | GH117 | 41,8 |
| | *Flammeovirga sp. MY04* | MY04_5110 | ANQ52445.1 | **S1_16*** | 58,4 |
| | *Flammeovirga sp. MY04* | MY04_5111 | ANQ52446.1 | **S1_4*** | 62,3 |
| | *Flammeovirga sp. MY04* | MY04_5112 | ANQ52447.1 | **S1_19*** | 53,3 |
| | *Flammeovirga sp. MY04* | MY04_5113 | ANQ52448.1 | GH2 | 92,9 |
| | *Flammeovirga sp. MY04* | MY04_5114 | ANQ52449.1 | **S1_19*** | 52,6 |
| | *Flammeovirga sp. MY04* | MY04_5115 | ANQ52450.1 | **S1_15*** | 57,1 |
| | *Flammeovirga sp. MY04* | MY04_5116 | ANQ52451.1 | GH136\|CBM1 | 121,5 |
| | *Flammeovirga sp. MY04* | MY04_5117 | ANQ52452.1 | GH95 | 89,3 |
| | *Flammeovirga sp. MY04* | MY04_5118 | ANQ52453.1 | **S1_19*** | 53,2 |
| | *Flammeovirga sp. MY04* | MY04_5119 | ANQ52454.1 | GH16 | 51,3 |
| | *Flammeovirga sp. MY04* | MY04_5120 | ANQ52455.1 | GH2 | 93,1 |
| | *Flammeovirga sp. MY04* | MY04_5121 | ANQ52456.1 | GH128 | 86,3 |
| | *Flammeovirga sp. MY04* | MY04_5124 | ANQ52459.1 | **S1_16*** | 60,2 |
| **Selected PUL 25: *Parabacteroides johnsonii CL02T12C29* (Predicted PUL 23)** | | | | | |
| **ECF-σ ► Anti-σ ► SusC ► SusC ► SusD ► GH16 ► unk ► unk ► GH97 ► GH106 ► GH117\|GH43_24 ► Sulf_1** | | | | | |
| Human gut | *Parabacteroides johnsonii CL02T12C29* | HMPREF1077_01244 | EKN12468.1 | GH16 | 30,9 |
| | *Parabacteroides johnsonii CL02T12C29* | HMPREF1077_01247 | EKN12471.1 | GH97 | 71,4 |
| | *Parabacteroides johnsonii CL02T12C29* | HMPREF1077_01249 | EKN12473.1 | GH117\|GH43 | 75,8 |
| | *Parabacteroides johnsonii CL02T12C29* | HMPREF1077_01250 | EKN12474.1 | **S1_4** | 57,9 |
| | *Parabacteroides johnsonii CL02T12C29* | HMPREF1077_01254 | EKN12478.1 | CE | 54,0 |
| | *Parabacteroides johnsonii CL02T12C29* | HMPREF1077_01255 | EKN12479.1 | GH105 | 42,5 |
| **Selected PUL 26: *Tamlana sp. UJ94* (Predicted PUL 13)** | | | | | |
| **SusC ► SusD ► Sulf_1 ► GH127 ► unk ► MFS ► unk ► Sulf_1 ► GH16 ► unk ► unk ► GH167 ► GH16 ►** | | | | | |
| Marine | *Tamlana carrageenivorans UJ94* | C1A40_08395 | AUS05484.1 | **S1_19** | 54,1 |
| | *Tamlana carrageenivorans UJ94* | C1A40_08400 | AUS05485.1 | GH127 | 75,6 |
| | *Tamlana carrageenivorans UJ94* | C1A40_08405 | AUS05486.1 | unk | 68,9 |
| | *Tamlana carrageenivorans UJ94* | C1A40_08420 | AUS07299.1 | **S1_7** | 63,6 |
| | *Tamlana carrageenivorans UJ94* | C1A40_08425 | AUS05489.1 | GH16_13 | 61,7 |
| | *Tamlana carrageenivorans UJ94* | C1A40_08430 | AUS05490.1 | unk | 68,4 |
| | *Tamlana carrageenivorans UJ94* | C1A40_08440 | AUS05491.1 | GH167 | 90,8 |
| | *Tamlana carrageenivorans UJ94* | C1A40_08445 | AUS05492.1 | GH16_17 | 36,2 |
| **Selected PUL 27: *Wenyingzhuangia fucanilytica CZ1127* (Predicted PUL 9)** | | | | | |
| **◄ GH167 ◄ GH167 ◄ GH110 ◄ unk ◄ unk ◄ Sulf_1 ◄ unk ◄ GH150 ◄ unk ◄ unk ◄ GH150 ◄ unk ◄ unk ◄** | | | | | |
| Marine | *Wenyingzhuangia fucanilytica CZ1127* | AXE80_08050 | ANW96232.1 | GH167 | 87,6 |
| | *Wenyingzhuangia fucanilytica CZ1127* | AXE80_08055 | ANW96233.1 | GH167 | 114,1 |
| | *Wenyingzhuangia fucanilytica CZ1127* | AXE80_08060 | ANW96234.1 | GH110 | 68,7 |
| | *Wenyingzhuangia fucanilytica CZ1127* | AXE80_08065 | ANW96235.1 | unk | 43,3 |
| | *Wenyingzhuangia fucanilytica CZ1127* | AXE80_08075 | ANW96237.1 | **S1_8** | 55,6 |
| | *Wenyingzhuangia fucanilytica CZ1127* | AXE80_08080 | ANW96238.1 | unk | 41,9 |
| | *Wenyingzhuangia fucanilytica CZ1127* | AXE80_08085 | ANW96239.1 | GH150 | 109,3 |
| | *Wenyingzhuangia fucanilytica CZ1127* | AXE80_08090 | ANW96240.1 | unk | 20,0 |
| | *Wenyingzhuangia fucanilytica CZ1127* | AXE80_08095 | ANW96241.1 | unk | 45,5 |
| | *Wenyingzhuangia fucanilytica CZ1127* | AXE80_08100 | ANW96242.1 | GH150 | 108,1 |
| | *Wenyingzhuangia fucanilytica CZ1127* | AXE80_08110 | ANW96244.1 | unk | 59,2 |
| | *Wenyingzhuangia fucanilytica CZ1127* | AXE80_08130 | ANW96248.1 | GH127 | 76,5 |
| | *Wenyingzhuangia fucanilytica CZ1127* | AXE80_08135 | ANW96249.1 | GH2 | 124,8 |
| **Selected PUL 28: *Bacteroides ovatus CL02T12C04* (Predicted PUL 25)** | | | | | |
| **◄ GntR ◄ Sulf_1 unk ► ◄ unk ◄ unk ◄ unk ◄ unk ◄ unk ◄ GH16 ◄ unk ◄ unk ◄ unk ◄ SusD ◄ SusC ◄ GH16 ◄ unk ◄ GH2 ◄ GH167 ◄ GH2 ◄ Sulf_1 ◄ unk ◄ unk ◄ unk ◄ Sulf_1 ◄ Sulf_1 ◄ GH127 ◄ unk ◄ unk** | | | | | |
| | *Bacteroides ovatus CL02T12C04* | HMPREF1069_02086 | EIY64557.1 | **S1_30** | 60,8 |

| | | | | | |
|---|---|---|---|---|---|
| Human gut | *Bacteroides ovatus CL02T12C04* | HMPREF1069_02093 | EIY64564.1 | GH16_17 | 60,8 |
| | *Bacteroides ovatus CL02T12C04* | HMPREF1069_02094 | EIY64565.1 | unk | 74,4 |
| | *Bacteroides ovatus CL02T12C04* | HMPREF1069_02099 | EIY64570.1 | GH16_17 | 41,4 |
| | *Bacteroides ovatus CL02T12C04* | HMPREF1069_02101 | EIY64572.1 | GH2 | 123,9 |
| | *Bacteroides ovatus CL02T12C04* | HMPREF1069_02102 | EIY64573.1 | GH167 | 87,9 |
| | *Bacteroides ovatus CL02T12C04* | HMPREF1069_02103 | EIY64574.1 | GH2 | 98,1 |
| | *Bacteroides ovatus CL02T12C04* | HMPREF1069_02104 | EIY64575.1 | **S1_30** | 59,4 |
| | *Bacteroides ovatus CL02T12C04* | HMPREF1069_02108 | EIY64579.1 | **S1_16** | 54,5 |
| | *Bacteroides ovatus CL02T12C04* | HMPREF1069_02109 | EIY64580.1 | **S1_81** | 59,7 |
| | *Bacteroides ovatus CL02T12C04* | HMPREF1069_02110 | EIY64581.1 | GH127 | 78,3 |
| **Selected PUL 29: *Cyclobacterium marinum DSM 745* (Predicted PUL 24)** | | | | | |
| **unk ► PL10_1 ► PL1_2 ► GH127 ► unk ► ◄ unk Sulf_1 ► ◄ unk ◄ unk ◄ Sulf_1 ◄ SusD ◄ SusC** | | | | | |
| Marine | *Cyclobacterium marinum DSM 745* | Cycma_2891 | AEL26627.1 | unk | 47,8 |
| | *Cyclobacterium marinum DSM 745* | Cycma_2892 | AEL26628.1 | PL10_1 | 41,8 |
| | *Cyclobacterium marinum DSM 745* | Cycma_2893 | AEL26629.1 | PL1_2 | 50,4 |
| | *Cyclobacterium marinum DSM 745* | Cycma_2894 | AEL26630.1 | GH127 | 76,1 |
| | *Cyclobacterium marinum DSM 745* | Cycma_2895 | AEL26631.1 | unk | 25,1 |
| | *Cyclobacterium marinum DSM 745* | Cycma_2897 | AEL26633.1 | **S1_23*** | 56,7 |
| | *Cyclobacterium marinum DSM 745* | Cycma_2899 | AEL26635.1 | unk | 75,8 |
| | *Cyclobacterium marinum DSM 745* | Cycma_2900 | AEL26636.1 | **S1_8*** | 58,4 |
| **Selected PUL 30: *Nonlabens sp. Hel1_33_55* (Predicted PUL 1)** | | | | | |
| **HTCS ► SusC ► SusD ► GH3 ► Sulf_1 ► Sulf_1 ► Sulf_1 ► unk ► GH29 ► Sulf_1 ► unk ► Sulf_1 ► GH10 ► unk ► Sulf_1 ► PL ► Sulf_1 ► Sulf_1 ► unk ► unk ►** | | | | | |
| Marine | *Nonlabens sp. Hel1_33_55* | SAMN05192588_0640 | SCX99618.1 | GH3 | 87,2 |
| | *Nonlabens sp. Hel1_33_55* | SAMN05192588_0641 | SCX99637.1 | **S1_11*** | 64,0 |
| | *Nonlabens sp. Hel1_33_55* | SAMN05192588_0642 | SCX99659.1 | **S1_7*** | 59,7 |
| | *Nonlabens sp. Hel1_33_55* | SAMN05192588_0643 | SCX99673.1 | **S1_16*** | 54,2 |
| | *Nonlabens sp. Hel1_33_55* | SAMN05192588_0644 | SCX99690.1 | Esterase | 34,6 |
| | *Nonlabens sp. Hel1_33_55* | SAMN05192588_0645 | SCX99714.1 | GH29 | 68,4 |
| | *Nonlabens sp. Hel1_33_55* | SAMN05192588_0646 | SCX99726.1 | **S1_16*** | 53,3 |
| | *Nonlabens sp. Hel1_33_55* | SAMN05192588_0647 | SCX99748.1 | GH | 65,7 |
| | *Nonlabens sp. Hel1_33_55* | SAMN05192588_0648 | SCX99764.1 | **S1_16*** | 54,4 |
| | *Nonlabens sp. Hel1_33_55* | SAMN05192588_0649 | SCX99783.1 | CBM85\|GH10 | 66,7 |
| | *Nonlabens sp. Hel1_33_55* | SAMN05192588_0650 | SCX99797.1 | unk | 52,8 |
| | *Nonlabens sp. Hel1_33_55* | SAMN05192588_0651 | SCX99820.1 | **S1_8*** | 63,3 |
| | *Nonlabens sp. Hel1_33_55* | SAMN05192588_0652 | SCX99839.1 | PL | 60,8 |
| | *Nonlabens sp. Hel1_33_55* | SAMN05192588_0653 | SCX99860.1 | **S1_7*** | 56,7 |
| | *Nonlabens sp. Hel1_33_55* | SAMN05192588_0654 | SCX99878.1 | **S1_7*** | 59,1 |
| | *Nonlabens sp. Hel1_33_55* | SAMN05192588_0655 | SCX99895.1 | unk | 63,7 |
| **Selected PUL 31: *Bacteroides stercoris CC31F* (Predicted PUL 28)** | | | | | |
| **Sulf_1 ► PL15_2 ► GH95 ► ◄ unk SusC ► SusD ► PL15_2 ► unk ► unk ► unk ► PL15 ► HTCS ► PL15 ► SusC ► SusD ► unk ► ◄ unk unk ► PL15 ► ◄ unk GH29 ► unk ► GH88 ► PL33_1 ►** | | | | | |
| Human gut | *Bacteroides stercoris CC31F* | HMPREF1181_01721 | EPH20505.1 | **S1_9** | 52,2 |
| | *Bacteroides stercoris CC31F* | HMPREF1181_01722 | EPH20506.1 | PL15_2 | 99,2 |
| | *Bacteroides stercoris CC31F* | HMPREF1181_01723 | EPH20507.1 | GH95 | 83,1 |
| | *Bacteroides stercoris CC31F* | HMPREF1181_01727 | EPH20511.1 | PL15_2 | 99,3 |
| | *Bacteroides stercoris CC31F* | HMPREF1181_01730 | EPH20514.1 | unk | 63,0 |
| | *Bacteroides stercoris CC31F* | HMPREF1181_01731 | EPH20515.1 | PL15 | 96,9 |
| | *Bacteroides stercoris CC31F* | HMPREF1181_01733 | EPH20517.1 | PL15 | 96,2 |
| | *Bacteroides stercoris CC31F* | HMPREF1181_01738 | EPH20522.1 | unk | 86,6 |
| | *Bacteroides stercoris CC31F* | HMPREF1181_01739 | EPH20523.1 | PL15 | 95,0 |
| | *Bacteroides stercoris CC31F* | HMPREF1181_01741 | EPH20525.1 | GH29 | 54,3 |
| | *Bacteroides stercoris CC31F* | HMPREF1181_01742 | EPH20526.1 | unk | 26,0 |
| | *Bacteroides stercoris CC31F* | HMPREF1181_01743 | EPH20527.1 | GH88 | 47,7 |
| | *Bacteroides stercoris CC31F* | HMPREF1181_01744 | EPH20528.1 | PL33_1 | 74,5 |
| **Selected PUL 32: *Echinicola pacifica DSM 19836* (Predicted PUL 33)** | | | | | |
| **◄ SusD ◄ SusC ◄ Sulf_1 ◄ unk ◄ Sulf_1 ◄ Sulf_1 ◄ GH29 ◄ Sulf_1 ◄ unk ◄ Sulf_1 ◄ unk ◄ GH29 ◄ unk ◄ GH ◄ Sulf_1 ◄ Sulf_1 ◄ Sulf_1 ◄ Sulf_1 ◄ GH29 ◄ GH95 ◄ GH ◄ SusD ◄ SusC** | | | | | |
| e | *Echinicola pacifica DSM 19836* | B050DRAFT_01785 | WP_018474051.1 | **S1_25*** | 56,6 |
| | *Echinicola pacifica DSM 19836* | B050DRAFT_01786 | WP_083923270.1 | **S1_22*** | 21,3 |
| | *Echinicola pacifica DSM 19836* | B050DRAFT_01787 | WP_169336255.1 | **S1_22*** | 27,0 |
| | *Echinicola pacifica DSM 19836* | B050DRAFT_01788 | WP_083923272.1 | Esterase | 56,8 |
| | *Echinicola pacifica DSM 19836* | B050DRAFT_01789 | WP_229802408.1 | GH29 | 66,6 |
| | *Echinicola pacifica DSM 19836* | B050DRAFT_01790 | WP_157492747.1 | **S1_25*** | 61,9 |
| | *Echinicola pacifica DSM 19836* | B050DRAFT_01791 | WP_018474055.1 | unk | 37,5 |
| | *Echinicola pacifica DSM 19836* | B050DRAFT_01792 | WP_018474056.1 | **S1_22*** | 52,8 |

| | | | | | |
|---|---|---|---|---|---|
| Marine | *Echinicola pacifica DSM 19836* | B050DRAFT_01793 | WP_018474057.1 | unk | 37,9 |
| | *Echinicola pacifica DSM 19836* | B050DRAFT_01794 | WP_018474058.1 | GH29 | 70,1 |
| | *Echinicola pacifica DSM 19836* | B050DRAFT_01796 | WP_205744990.1 | GH141 | 81,6 |
| | *Echinicola pacifica DSM 19836* | B050DRAFT_01797 | WP_018474060.1 | **S1_15*** | 58,0 |
| | *Echinicola pacifica DSM 19836* | B050DRAFT_01798 | WP_018474061.1 | **S1_17*** | 67,9 |
| | *Echinicola pacifica DSM 19836* | B050DRAFT_01799 | WP_018474062.1 | **S1_17*** | 0,0 |
| | *Echinicola pacifica DSM 19836* | B050DRAFT_01800 | WP_083923274.1 | **S1_17*** | 0,0 |
| | *Echinicola pacifica DSM 19836* | B050DRAFT_01801 | WP_044202327.1 | GH29 | 42,3 |
| | *Echinicola pacifica DSM 19836* | B050DRAFT_01802 | WP_205744991.1 | GH95 | 108,4 |
| | *Echinicola pacifica DSM 19836* | B050DRAFT_01803 | WP_018474066.1 | GH141 | 82,6 |
| **Selected PUL 33:** ***Wenyingzhuangia fucanilytica CZ1127*** **(Predicted PUL 8)** | | | | | |
| **◄ GH ◄ GH107 ◄ GH107 ◄ unk ◄ unk ◄ unk ◄ unk ◄ Sulf_1 ◄ Sulf_1 ◄ unk ◄ GH29 ◄ GH95 ◄ unk ◄ unk ◄ GH ◄ GH ◄ GH29 ◄ Sulf_1 ◄ SusD ◄ SusC ◄ Sulf_1 ◄ unk ◄ GH29 ◄ Sulf_1 ◄ GH107 ◄ GH107** | | | | | |
| Marine | *Wenyingzhuangia fucanilytica CZ1127* | AXE80_07300 | ANW96096.1 | GH141 | 80,8 |
| | *Wenyingzhuangia fucanilytica CZ1127* | AXE80_07305 | ANW96097.1 | GH107 | 91,0 |
| | *Wenyingzhuangia fucanilytica CZ1127* | AXE80_07310 | ANW96098.1 | GH107 | 96,4 |
| | *Wenyingzhuangia fucanilytica CZ1127* | AXE80_07315 | ANW96099.1 | GH141 | 80,8 |
| | *Wenyingzhuangia fucanilytica CZ1127* | AXE80_07320 | WP_083194610.1 | GH43_2 | 34,1 |
| | *Wenyingzhuangia fucanilytica CZ1127* | AXE80_07330 | ANW96101.1 | unk | 88,4 |
| | *Wenyingzhuangia fucanilytica CZ1127* | AXE80_07345 | ANW97461.1 | GH141 | 82,8 |
| | *Wenyingzhuangia fucanilytica CZ1127* | AXE80_07350 | ANW97462.1 | GH29 | 43,9 |
| | *Wenyingzhuangia fucanilytica CZ1127* | AXE80_07355 | ANW96103.1 | GH95 | 101,7 |
| | *Wenyingzhuangia fucanilytica CZ1127* | AXE80_07365 | ANW96105.1 | GH168 | 44,0 |
| | *Wenyingzhuangia fucanilytica CZ1127* | AXE80_07370 | ANW96106.1 | unk | 59,0 |
| | *Wenyingzhuangia fucanilytica CZ1127* | AXE80_07375 | ANW96107.1 | GH117 | 39,8 |
| | *Wenyingzhuangia fucanilytica CZ1127* | AXE80_07380 | ANW96108.1 | GH29 | 69,2 |
| | *Wenyingzhuangia fucanilytica CZ1127* | AXE80_07405 | ANW96112.1 | unk | 32,3 |
| | *Wenyingzhuangia fucanilytica CZ1127* | AXE80_07410 | ANW96113.1 | GH29 | 55,9 |
| | *Wenyingzhuangia fucanilytica CZ1127* | AXE80_07420 | ANW96115.1 | GH107 | 109,2 |
| | *Wenyingzhuangia fucanilytica CZ1127* | AXE80_07425 | ANW96116.1 | GH107 | 87,4 |
| **Selected PUL 34:** ***Flagellimonas eckloniae DOKDO 007*** **(Predicted PUL 6)** | | | | | |
| **◄ GntR ◄ PL7 ◄ unk ◄ unk ◄ SusD ◄ SusC ◄ unk ◄ PL17_2 ◄ PL6_1\|PL6_1 unk ► ◄ GH92 ◄ unk ◄ CBM9 ◄ GH20 ◄ GH92 ◄ GH20 ◄ GH29 ◄ Sulf_1 ◄ GH92 ◄ GH3 ◄ Sulf_1 ◄ GH95 ◄ Sulf_1 ◄ Sulf_1 ◄ GH29 ◄** | | | | | |
| Marine | *Flagellimonas eckloniae DOKDO 007* | AAY42_07765 | KQC31659.1 | GH92 | 84,4 |
| | *Flagellimonas eckloniae DOKDO 007* | AAY42_07770 | KQC31660.1 | unk | 42,5 |
| | *Flagellimonas eckloniae DOKDO 007* | AAY42_07775 | KQC29796.1 | CBM9 | 41,5 |
| | *Flagellimonas eckloniae DOKDO 007* | AAY42_07780 | KQC29797.1 | GH20 | 88,0 |
| | *Flagellimonas eckloniae DOKDO 007* | AAY42_07785 | KQC29798.1 | GH92 | 84,3 |
| | *Flagellimonas eckloniae DOKDO 007* | AAY42_07790 | KQC31661.1 | GH20 | 87,9 |
| | *Flagellimonas eckloniae DOKDO 007* | AAY42_07795 | KQC29799.1 | GH29 | 50,0 |
| | *Flagellimonas eckloniae DOKDO 007* | AAY42_07800 | KQC31662.1 | **S1_15*** | 53,2 |
| | *Flagellimonas eckloniae DOKDO 007* | AAY42_07805 | KQC31663.1 | GH92 | 88,2 |
| | *Flagellimonas eckloniae DOKDO 007* | AAY42_07810 | KQC31664.1 | GH3 | 100,0 |
| | *Flagellimonas eckloniae DOKDO 007* | AAY42_07815 | KQC29800.1 | **S1_14*** | 49,2 |
| | *Flagellimonas eckloniae DOKDO 007* | AAY42_07820 | KQC31665.1 | GH95 | 86,8 |
| | *Flagellimonas eckloniae DOKDO 007* | AAY42_07825 | KQC29801.1 | **S1_17 *** | 68,9 |
| | *Flagellimonas eckloniae DOKDO 007* | AAY42_07830 | KQC31666.1 | **S1_11*** | 59,6 |
| | *Flagellimonas eckloniae DOKDO 007* | AAY42_07835 | WP_139063687.1 | GH29 | 48,9 |
| | *Flagellimonas eckloniae DOKDO 007* | AAY42_07840 | KQC29802.1 | **S1_28*** | 56,1 |
| | *Flagellimonas eckloniae DOKDO 007* | AAY42_07845 | KQC31667.1 | **S1_11*** | 60,2 |
| | *Flagellimonas eckloniae DOKDO 007* | AAY42_07850 | KQC29803.1 | GH130 | 39,0 |
| | *Flagellimonas eckloniae DOKDO 007* | AAY42_07855 | KQC29804.1 | unk | 72,9 |
| | *Flagellimonas eckloniae DOKDO 007* | AAY42_07860 | KQC29805.1 | GH92 | 84,7 |
| **Selected PUL 35:** ***Flexithrix dorotheae DSM 6795*** **(Predicted PUL 40)** | | | | | |
| **unk ► unk ► unk ► Sulf_4 ► ◄ unk ◄ GH28 ◄ Sulf_1 ◄ Sulf_1 ◄ Sulf_1 ◄ unk ◄ Sulf_1 ◄ GH ◄ unk ◄ unk** | | | | | |
| Marine | *Flexithrix dorotheae DSM 6795* | A3EMDRAFT_05972 | WP_020529959.1 | unk | 71,5 |
| | *Flexithrix dorotheae DSM 6795* | A3EMDRAFT_05973 | WP_083925270.1 | GH28 | 55,2 |
| | *Flexithrix dorotheae DSM 6795* | A3EMDRAFT_05974 | WP_052323370.1 | **S1_14*** | 53,3 |
| | *Flexithrix dorotheae DSM 6795* | A3EMDRAFT_05975 | WP_044239112.1 | **S1_20*** | 52,4 |
| | *Flexithrix dorotheae DSM 6795* | A3EMDRAFT_05976 | WP_020529963.1 | **S1_16*** | 56,5 |
| | *Flexithrix dorotheae DSM 6795* | A3EMDRAFT_05978 | WP_020529965.1 | **S1_20*** | 51,9 |
| | *Flexithrix dorotheae DSM 6795* | A3EMDRAFT_05979 | WP_020529966.1 | unk | 61,6 |
| | *Flexithrix dorotheae DSM 6795* | A3EMDRAFT_05980 | WP_026210427.1 | unk | 41,9 |
| | *Flexithrix dorotheae DSM 6795* | A3EMDRAFT_05981 | WP_026210428.1 | unk | 74,9 |
| | *Flexithrix dorotheae DSM 6795* | A3EMDRAFT_05982 | WP_020529969.1 | GH42 | 81,7 |
| | *Flexithrix dorotheae DSM 6795* | A3EMDRAFT_05983 | WP_020529970.1 | unk | 124,9 |

| | | | | | |
|---|---|---|---|---|---|
| | *Flexithrix dorotheae DSM 6795* | A3EMDRAFT_05987 | WP_020529975.1 | GH29 | 43,0 |
| | *Flexithrix dorotheae DSM 6795* | A3EMDRAFT_05988 | WP_020529976.1 | PL29 | 61,5 |
| | *Flexithrix dorotheae DSM 6795* | A3EMDRAFT_05989 | WP_026210429.1 | GH88 | 47,7 |
| | *Flexithrix dorotheae DSM 6795* | A3EMDRAFT_05991 | WP_157637982.1 | unk | 83,5 |
| | *Flexithrix dorotheae DSM 6795* | A3EMDRAFT_05992 | WP_044239140.1 | unk | 87,4 |
| **Selected PUL 36: *Zobellia galactanivorans DsijT* (Predicted PUL 9)** | | | | | |
| **◄ GH ◄ unk ◄ Sulf_1 ◄ PL\|CBM16 ◄ GH29\|CBM47 ◄ GH29\|CBM47 ◄ unk ◄ GH29 ◄ GH29 ◄ GH95 ◄ GH3 ◄** | | | | | |
| Marine | *Zobellia galactanivorans DsijT* | ZOBELLIA_334 | CAZ94407.1 | unk | 80,3 |
| | *Zobellia galactanivorans DsijT* | ZOBELLIA_335 | CAZ94408.1 | unk | 85,2 |
| | *Zobellia galactanivorans DsijT* | ZOBELLIA_336 | CAZ94409.1 | **S1_16** | 54,9 |
| | *Zobellia galactanivorans DsijT* | ZOBELLIA_337 | CAZ94410.1 | unk | 87,5 |
| | *Zobellia galactanivorans DsijT* | ZOBELLIA_338 | CAZ94411.1 | GH29\|CBM47 | 55,0 |
| | *Zobellia galactanivorans DsijT* | ZOBELLIA_339 | CAZ94412.1 | GH29\|CBM47 | 56,2 |
| | *Zobellia galactanivorans DsijT* | ZOBELLIA_340 | CAZ94413.1 | GH141 | 90,4 |
| | *Zobellia galactanivorans DsijT* | ZOBELLIA_341 | CAZ94414.1 | GH29 | 41,5 |
| | *Zobellia galactanivorans DsijT* | ZOBELLIA_342 | CAZ94415.1 | GH29 | 56,4 |
| | *Zobellia galactanivorans DsijT* | ZOBELLIA_343 | CAZ94416.1 | GH95 | 87,7 |
| | *Zobellia galactanivorans DsijT* | ZOBELLIA_344 | CAZ94417.1 | GH3 | 81,9 |
| | *Zobellia galactanivorans DsijT* | ZOBELLIA_345 | CAZ94418.1 | GH29 | 56,4 |
| | *Zobellia galactanivorans DsijT* | ZOBELLIA_346 | CAZ94419.1 | GH29\|CBM51 | 74,4 |
| | *Zobellia galactanivorans DsijT* | ZOBELLIA_347 | CAZ94420.1 | GH3 | 89,2 |
| **Selected PUL 37: *Saccharicrinis fermentans DSM 9555* (Predicted PUL 14)** | | | | | |
| **◄ PL40 ◄ PL40 ◄ GH43 ◄ GH29 ◄ unk ◄ GH30_2 ◄ unk ◄ GH97 ◄ SusD ◄ SusC ◄ unk ◄ unk ◄ SusD ◄** | | | | | |
| Marine | *Saccharicrinis fermentans DSM 9555* | CytfeDRAFT_1464 | GAF02015.1 | PL33_2 | 78,8 |
| | *Saccharicrinis fermentans DSM 9555* | CytfeDRAFT_1465 | GAF02014.1 | **S1_17** | 67,9 |
| | *Saccharicrinis fermentans DSM 9555* | CytfeDRAFT_1467 | GAF02012.1 | **S1_15** | 59,5 |
| | *Saccharicrinis fermentans DSM 9555* | CytfeDRAFT_1468 | GAF02011.1 | **S1_17** | 68,3 |
| | *Saccharicrinis fermentans DSM 9555* | CytfeDRAFT_1469 | GAF02010.1 | **S1_23** | 54,7 |
| | *Saccharicrinis fermentans DSM 9555* | CytfeDRAFT_1471 | GAF02008.1 | **S1_28** | 55,7 |
| | *Saccharicrinis fermentans DSM 9555* | CytfeDRAFT_1472 | WP_052343062.1 | GH95 | 63,9 |
| | *Saccharicrinis fermentans DSM 9555* | CytfeDRAFT_1477 | GAF01999.1 | PL40 | 105,7 |
| | *Saccharicrinis fermentans DSM 9555* | CytfeDRAFT_1478 | WP_211238157.1 | PL40 | 108,0 |
| | *Saccharicrinis fermentans DSM 9555* | CytfeDRAFT_1479 | GAF01996.1 | GH43 | 36,5 |
| | *Saccharicrinis fermentans DSM 9555* | CytfeDRAFT_1480 | GAF01995.1 | GH29 | 51,9 |
| | *Saccharicrinis fermentans DSM 9555* | CytfeDRAFT_1481 | WP_052343064.1 | unk | 28,5 |
| | *Saccharicrinis fermentans DSM 9555* | CytfeDRAFT_1482 | GAF01993.1 | GH30_2 | 54,4 |
| | *Saccharicrinis fermentans DSM 9555* | CytfeDRAFT_1484 | GAF01991.1 | GH97 | 74,0 |
| | *Saccharicrinis fermentans DSM 9555* | CytfeDRAFT_1487 | GAF01988.1 | unk | 56,7 |
| | *Saccharicrinis fermentans DSM 9555* | CytfeDRAFT_1488 | GAF01987.1 | unk | 60,6 |
| | *Saccharicrinis fermentans DSM 9555* | CytfeDRAFT_1491 | GAF01983.1 | unk | 86,9 |
| | *Saccharicrinis fermentans DSM 9555* | CytfeDRAFT_1493 | GAF01980.1 | GH88 | 45,3 |
| | *Saccharicrinis fermentans DSM 9555* | CytfeDRAFT_1494 | GAF01979.1 | CE | 60,8 |
| | *Saccharicrinis fermentans DSM 9555* | CytfeDRAFT_1495 | GAF01978.1 | **S1_11** | 64,0 |
| | *Saccharicrinis fermentans DSM 9555* | CytfeDRAFT_1496 | GAF01977.1 | CBM6\|CBM6\| | 141,5 |
| | *Saccharicrinis fermentans DSM 9555* | CytfeDRAFT_1497 | WP_052343067.1 | unk | 131,9 |
| **Selected PUL 38: *Niabella soli DSM 19437* (Cazymes cluster 5)** | | | | | |
| **◄ GH28 ◄ GH29 ◄ PL6_2 ◄ unk ◄ GH88 ◄ GH29 ◄ PL38 ◄ GH31 ◄ GH95 ◄ GH2 ◄ unk ◄ SusD ◄ SusC** | | | | | |
| Soil | *Niabella soli DSM 19437* | Niaso_1999 | AHF15512.1 | unk | 55,1 |
| | *Niabella soli DSM 19437* | Niaso_2000 | AHF15513.1 | GH105 | 43,0 |
| | *Niabella soli DSM 19437* | Niaso_2002 | AHF15515.1 | GH29 | 66,7 |
| | *Niabella soli DSM 19437* | Niaso_2003 | AHF15516.1 | unk | 72,9 |
| | *Niabella soli DSM 19437* | Niaso_2004 | AHF15517.1 | unk | 94,6 |
| | *Niabella soli DSM 19437* | Niaso_2005 | AHF15518.1 | unk | 53,9 |
| | *Niabella soli DSM 19437* | Niaso_2006 | AHF15519.1 | GH28 | 55,8 |
| | *Niabella soli DSM 19437* | Niaso_2007 | AHF15520.1 | GH29 | 52,8 |
| | *Niabella soli DSM 19437* | Niaso_2008 | AHF15521.1 | PL6_2 | 49,4 |
| | *Niabella soli DSM 19437* | Niaso_2009 | AHF15522.1 | unk | 51,2 |
| | *Niabella soli DSM 19437* | Niaso_2010 | AHF15523.1 | GH88 | 45,7 |
| | *Niabella soli DSM 19437* | Niaso_2011 | AHF15524.1 | GH29 | 70,5 |
| | *Niabella soli DSM 19437* | Niaso_2012 | AHF15525.1 | PL38 | 45,3 |
| | *Niabella soli DSM 19437* | Niaso_2013 | AHF15526.1 | GH31 | 80,5 |
| | *Niabella soli DSM 19437* | Niaso_2014 | AHF15527.1 | GH95 | 95,7 |
| | *Niabella soli DSM 19437* | Niaso_2015 | AHF15528.1 | GH2 | 92,1 |
| | *Niabella soli DSM 19437* | Niaso_2016 | AHF17606.1 | unk | 25,8 |
| | *Niabella soli DSM 19437* | Niaso_2019 | AHF15530.1 | GH28 | 102,9 |

| | | | | | |
|---|---|---|---|---|---|
| **Selected PUL 39:** ***Bacteroides clarus YIT 12056*** **(Predicted PUL 41)** | | | | | |
| **◄ PL33_1 ◄ GH88 ◄ unk ◄ GH95 ◄ GH36 ◄ GH29 ◄ SusD ◄ SusC ◄ PL15 ◄ GH95 ◄ HTCS ◄ HTCS ◄ GH88** | | | | | |
| Human gut | *Bacteroides clarus YIT 12056* | HMPREF9445_03068 | EGF49696.1 | PL33_1 | 73,9 |
| | *Bacteroides clarus YIT 12056* | HMPREF9445_03069 | EGF49697.1 | GH88 | 47,6 |
| | *Bacteroides clarus YIT 12056* | HMPREF9445_03070 | EGF49698.1 | unk | 27,2 |
| | *Bacteroides clarus YIT 12056* | HMPREF9445_03071 | EGF49699.1 | GH95 | 84,0 |
| | *Bacteroides clarus YIT 12056* | HMPREF9445_03072 | EGF49700.1 | GH36 | 86,6 |
| | *Bacteroides clarus YIT 12056* | HMPREF9445_03073 | EGF49701.1 | GH29 | 63,2 |
| | *Bacteroides clarus YIT 12056* | HMPREF9445_03076 | EGF49704.1 | PL15 | 96,6 |
| | *Bacteroides clarus YIT 12056* | HMPREF9445_03077 | EGF49705.1 | GH95 | 103,5 |
| | *Bacteroides clarus YIT 12056* | HMPREF9445_03080 | EGF49708.1 | GH88 | 46,7 |
| | *Bacteroides clarus YIT 12056* | HMPREF9445_03081 | EGF49709.1 | **S1_14** | 51,8 |
| | *Bacteroides clarus YIT 12056* | HMPREF9445_03082 | EGF49710.1 | unk | 112,9 |
| | *Bacteroides clarus YIT 12056* | HMPREF9445_03083 | EGF49711.1 | PL15 | 98,1 |
| **Selected PUL 40:** ***Cellulophaga baltica DSM 24729*** **(Predicted PUL 22)** | | | | | |
| **unk ► unk ► unk ► unk ► GH29 ► SusC ► SusD ► unk ► GH16 ► Sulf_1 ► unk ► GH29 ► unk ► GH16 ►** | | | | | |
| Marine | *Cellulophaga baltica DSM 24729* | Ga0070226_10966 | SDF23599.1 | unk | 122,9 |
| | *Cellulophaga baltica DSM 24729* | Ga0070226_10967 | SDF23614.1 | unk | 58,7 |
| | *Cellulophaga baltica DSM 24729* | Ga0070226_10968 | SDF23646.1 | unk | 41,0 |
| | *Cellulophaga baltica DSM 24729* | Ga0070226_10969 | SDF23678.1 | unk | 52,8 |
| | *Cellulophaga baltica DSM 24729* | Ga0070226_10970 | SDF23691.1 | GH29 | 57,4 |
| | *Cellulophaga baltica DSM 24729* | Ga0070226_10973 | SDF23759.1 | unk | 57,2 |
| | *Cellulophaga baltica DSM 24729* | Ga0070226_10974 | SDF23778.1 | GH16_16 | 37,9 |
| | *Cellulophaga baltica DSM 24729* | Ga0070226_10975 | SDF23801.1 | **S1_11** | 64,1 |
| | *Cellulophaga baltica DSM 24729* | Ga0070226_10976 | SDF23816.1 | unk | 63,9 |
| | *Cellulophaga baltica DSM 24729* | Ga0070226_10977 | SDF23840.1 | GH29 | 59,0 |
| | *Cellulophaga baltica DSM 24729* | Ga0070226_10978 | SDF23855.1 | unk | 33,4 |
| | *Cellulophaga baltica DSM 24729* | Ga0070226_10979 | SDF23882.1 | GH16_11 | 32,6 |
| | *Cellulophaga baltica DSM 24729* | Ga0070226_10980 | SDF23901.1 | **S1_15** | 55,9 |
| | *Cellulophaga baltica DSM 24729* | Ga0070226_10981 | SDF23921.1 | GH29 | 60,1 |
| | *Cellulophaga baltica DSM 24729* | Ga0070226_10982 | SDF23942.1 | GH29 | 57,3 |
| | *Cellulophaga baltica DSM 24729* | Ga0070226_10983 | SDF23970.1 | unk | 52,1 |
| | *Cellulophaga baltica DSM 24729* | Ga0070226_10984 | SDF23985.1 | **S1_28** | 58,5 |
| | *Cellulophaga baltica DSM 24729* | Ga0070226_10985 | SDF24009.1 | GH117 | 47,0 |
| | *Cellulophaga baltica DSM 24729* | Ga0070226_10986 | SDF24033.1 | GH28 | 52,2 |
| | *Cellulophaga baltica DSM 24729* | Ga0070226_10987 | SDF24056.1 | GH16_11 | 33,6 |
| | *Cellulophaga baltica DSM 24729* | Ga0070226_10988 | SDF24072.1 | **S1_8** | 70,7 |
| | *Cellulophaga baltica DSM 24729* | Ga0070226_10989 | SDF24099.1 | GH28 | 51,9 |
| | *Cellulophaga baltica DSM 24729* | Ga0070226_10990 | SDF24118.1 | **S1_28** | 49,2 |
| | *Cellulophaga baltica DSM 24729* | Ga0070226_10991 | SDF24137.1 | **S1_24** | 24,8 |
| | *Cellulophaga baltica DSM 24729* | Ga0070226_10992 | SDF24160.1 | GH2 | 94,0 |
| | *Cellulophaga baltica DSM 24729* | Ga0070226_10993 | SDF24178.1 | GH16_11 | 38,9 |
| | *Cellulophaga baltica DSM 24729* | Ga0070226_10994 | SDF24205.1 | **S1_25** | 48,4 |
| | *Cellulophaga baltica DSM 24729* | Ga0070226_10995 | SDF24225.1 | **S1_25*** | 15,3 |
| | *Cellulophaga baltica DSM 24729* | Ga0070226_10996 | SDF24245.1 | **S1_17** | 69,3 |
| | *Cellulophaga baltica DSM 24729* | Ga0070226_10997 | SDF24267.1 | GH2 | 108,5 |
| | *Cellulophaga baltica DSM 24729* | Ga0070226_10998 | SDF24289.1 | **S1_15** | 58,4 |
| | *Cellulophaga baltica DSM 24729* | Ga0070226_10999 | SDF24306.1 | **S1_19** | 52,4 |
| | *Cellulophaga baltica DSM 24729* | Ga0070226_109100 | SDF24334.1 | GH2 | 93,1 |
| | *Cellulophaga baltica DSM 24729* | Ga0070226_109101 | SDF24350.1 | **S1_28** | 57,2 |
| **Selected PUL 41:** ***Reichenbachiella agariperforans DSM 26134*** **(Predicted PUL 21)** | | | | | |
| **SusC ► SusD ► SusC ► SusD ► unk ► unk ► GH29 ► Sulf_1 ► Sulf_1 ► PL ► GH28 ► GH28 ► Sulf_1 ► GH2** | | | | | |
| Marine | *Reichenbachiella agariperforans DSM 26134* | Ga0070003_10784 | SHK67534.1 | unk | 33,6 |
| | *Reichenbachiella agariperforans DSM 26134* | Ga0070003_10786 | SHK67590.1 | GH29 | 58,3 |
| | *Reichenbachiella agariperforans DSM 26134* | Ga0070003_10787 | SHK67621.1 | **S1_11** | 64,5 |
| | *Reichenbachiella agariperforans DSM 26134* | Ga0070003_10788 | SHK67653.1 | **S1_19** | 52,6 |
| | *Reichenbachiella agariperforans DSM 26134* | Ga0070003_10789 | SHK67680.1 | unk | 81,6 |
| | *Reichenbachiella agariperforans DSM 26134* | Ga0070003_10790 | SHK67709.1 | GH28 | 51,6 |
| | *Reichenbachiella agariperforans DSM 26134* | Ga0070003_10791 | SHK67740.1 | GH28 | 51,2 |
| | *Reichenbachiella agariperforans DSM 26134* | Ga0070003_10792 | SHK67771.1 | **S1_28** | 59,0 |
| | *Reichenbachiella agariperforans DSM 26134* | Ga0070003_10793 | SHK67796.1 | GH2 | 109,0 |
| | *Reichenbachiella agariperforans DSM 26134* | Ga0070003_10794 | SHK67824.1 | GH29 | 57,4 |
| | *Reichenbachiella agariperforans DSM 26134* | Ga0070003_10795 | SHK67854.1 | GH29 | 57,9 |
| | *Reichenbachiella agariperforans DSM 26134* | Ga0070003_10796 | SHK67883.1 | **S1_15** | 55,8 |
| | *Reichenbachiella agariperforans DSM 26134* | Ga0070003_10797 | SHK67911.1 | unk | 82,5 |
| | *Reichenbachiella agariperforans DSM 26134* | Ga0070003_10798 | SHK67940.1 | GH2 | 92,5 |

| | | | | | |
|---|---|---|---|---|---|
| | *Reichenbachiella agariperforans DSM 26134* | Ga0070003_10799 | SHK67965.1 | GH16 | 33,6 |
| | *Reichenbachiella agariperforans DSM 26134* | Ga0070003_107100 | SHK67996.1 | **S1_8** | 71,4 |
| | *Reichenbachiella agariperforans DSM 26134* | Ga0070003_107101 | SHK68026.1 | **S1_72** | 89,1 |
