## Supplementary Table 2 for "Screening for Polysaccharide Utilization Loci Targeting Marine Polysaccharides"

**Table S2:** Activity screening of the enzymes grouped in 31 established CAZy families. * Not in PUL.

| Family | Organism | Genbank | Substrate | Activity | EC number |
|---|---|---|---|---|---|
| **GH2** | *Barnesiella viscericola DSM 18177* | AHF11965.1 | pNP-b-D-Galp | b-D-galactopyranosidase | 3.2.1.23 |
| **GH2** | *Flavobacterium johnsoniae UW101* | ABQ05112.1 | pNP-b-D-GlcA | b-D-glucuropyranosidase | 3.2.1.31 |
| **GH2** | *Saccharicrinis fermentans DSM 9555* | WP_081736079.1 | pNP-b-D-Galp | b-D-galactopyranosidase | 3.2.1.23 |
| **GH2** | *Bacteroides cellulosilyticus WH2* | ALJ59590.1 | pNP-b-D-Galp | b-D-galactopyranosidase | 3.2.1.23 |
| **GH2** | *Bacteroides cellulosilyticus WH2* | ALJ59608.1 | b(1,3)-Galactan | b-D-1,3-galactopyranosidase | 3.2.1.23 |
| **GH2** | *Bacteroides thetaiotaomicron 7330* | ALJ41002.1 | pNP-b-D-Galp | b-D-galactopyranosidase | 3.2.1.23 |
| **GH2** | *Polaribacter reichenbachii KCTC 23969* | AUC20527.1 | pNP-b-D-Galp | b-D-galactopyranosidase | 3.2.1.23 |
| **GH2** | *Algibacter lectus JCM 19300* | GAL64206.1 | pNP-a-D-GlcA | b-D-glucuropyranosidase | 3.2.1.31 |
| **GH2** | *Flammeovirga sp. MY04* | ANQ52455.1 | pNP-b-D-Galp | b-D-galactopyranosidase | 3.2.1.23 |
| **GH2** | *Wenyingzhuangia fucanilytica CZ1127* | ANW96249.1 | pNP-b-D-Galp | b-D-galactopyranosidase | 3.2.1.23 |
| **GH2** | *Bacteroides ovatus CL02T12C04* | EIY64572.1 | pNP-b-D-Galp | b-D-galactopyranosidase | 3.2.1.23 |
| **GH2** | *Bacteroides ovatus CL02T12C04* | EIY64574.1 | pNP-b-D-Galp | b-D-galactopyranosidase | 3.2.1.23 |
| **GH2** | *Niabella soli DSM 19437* | AHF15528.1 | pNP-b-D-GlcA | b-D-galactopyranosidase, a-L-arabinopyranosidase | 3.2.1.23 |
| **GH2** | *Cellulophaga baltica DSM 24729* | SDF24160.1 | pNP-b-D-Galp | b-D-galactopyranosidase, a-L-arabinopyranosidase | 3.2.1.23 |
| **GH2** | *Reichenbachiella agariperforans DSM 26134* | SHK67940.1 | pNP-b-D-Galp | b-D-galactopyranosidase, b-D-fucopyranosidase | 3.2.1.23 |
| **GH2\|CBM57** | *Bacteroides ovatus ATCC 8483* | ALJ46942.1 | pNP-b-D-GlcA | b-D-glucuropyranosidase | 3.2.1.31 |
| **GH2\|CBM57** | *Bacteroides ovatus ATCC 8483* | ALJ48503.1 | pNP-b-D-GlcA | b-D-glucuropyranosidase | 3.2.1.31 |
| **GH3** | *Polaribacter sejongensis KCTC 23670* | AUC23902.1 | pNP- b-D-Glcp, pNP-b-D-Xylp | b-D-glucopyranosidase, b-D-xylopyranosidase, | 3.2.1.21 3.2.1.37 |
| **GH3** | *Nonlabens sp. Hel1_33_55* | SCX99618.1 | pNP-b-D-Glcp, pNP-b-D-cellobiosides | b-D-glucopyranosidase | 3.2.1.21 |
| **GH10** | *Flavobacterium johnsoniae UW101* | ABQ05109.1 | pNP-b-D-Cellobiose, Arabinoxylan | Arabinoxylanase | 3.2.1.- |
| **GH16** | *Cellulophaga baltica DSM 24729* | SDF23778.1 | Porphyran, agarose | b-agarase | 3.2.1.81 |
| **GH16** | *Reichenbachiella agariperforans DSM 26134* | SHK67965.1 | Porphyran, oligo-porphyran | porphyranase | 3.2.1.178 |
| **GH16_11** | *Polaribacter reichenbachii KCTC 23969* | AUC19427.1 | Porphyran | Porphyranase | 3.2.1.178 |
| **GH16_11** | *Cellulophaga baltica DSM 24729* | SDF24056.1 | Porphyran | porphyranase | 3.2.1.178 |
| **GH16_13** | *Tamlana carrageenivorans UJ94* | AUS05489.1 | a-carrageenan | a-carrageenase | 3.2.1. |
| **GH16_17** | *Tamlana carrageenivorans UJ94* | AUS05492.1 | k-carrageenan | k-carrageenase | 3.2.1.83 |
| **GH16_17** | *Bacteroides ovatus CL02T12C04* | EIY64570.1 | k-carrageenan, i-carrageenan | k-carrageenase, i-carrageenase | 3.2.1.83, 3.2.1.157 |
| **GH20** | *Barnesiella viscericola DSM 18177* | AHF11967.1 | pNP-b-D-GlcNAc6S, pNP-b-D-GlcNAc, pNP-b-D-GalNAc | b-D-N-Acetyl-6-sulfo-hexosaminidase | 3.2.1.52 3.2.1.- |
| **GH20** | *Bacteroides cellulosilyticus WH2* | ALJ59591.1 | pNP-b-D-GlcNAc6S, pNP-b-D-GlcNAc, pNP-b-D-GalNAc | b-D-N-Acetyl-6-sulfo-hexosaminidase | 3.2.1.52 3.2.1.- |
| **GH20** | *Bacteroides thetaiotaomicron 7330* | ALJ41001.1 | pNP-b-D-GlcNAc6S, pNP-b-D-GlcNAc, pNP-b-D-GalNAc | b-D-N-Acetyl-6-sulfo-hexosaminidase | 3.2.1.52 3.2.1.- |
| **GH20** | *Parabacteroides gordonii DSM 23371* | KKB57913.1 | pNP-b-D-GlcNAc, pNP-b-D-GalNAc | b-D-N-Acetyl-hexosaminidase | 3.2.1.52 |
| **GH20** | *Phocaeicola dorei DSM 17855* | EEB26055.1 | pNP-b-D-GlcNAc6S, pNP-b-D-GlcNAc, pNP-b-D-GalNAc | b-D-N-Acetyl-6-sulfo-hexosaminidase | 3.2.1.52 3.2.1.- |
| **GH20** | *Chitinophaga pinensis DSM 2588* | ACU62427.1 | pNP-b-D-GlcNAc, pNP-b-D-GalNAc | b-D-N-Acetyl-hexosaminidase | 3.2.1.52 |
| **GH20** | *Flagellimonas eckloniae DOKDO 007* | KQC29797.1 | pNP-b-D-GlcNAc, pNP-b-D-GalNAc | b-D-N-Acetyl-hexosaminidase | 3.2.1.52 |
| **GH20** | *Flagellimonas eckloniae DOKDO 007* | KQC31661.1 | pNP-b-D-GlcNAc, pNP-b-D-GalNAc | b-D-N-Acetyl-hexosaminidase | 3.2.1.52 |
| **GH28** | *Polaribacter reichenbachii KCTC 23969* | AUC19434.1 | pNP-a-D-GlcA | a-D-glucuronidase | 3.2.1.139 |
| **GH28** | *Sphingobacterium thalpophilum DSM 11723* | SDH45426.1 | Polygalacturonic acid (RG I), pNP-a-D-GlcA | a-D-1,4-galacturonase | 3.2.1.15 |
| **GH29** | *Wenyingzhuangia fucanilytica CZ1127* | ANW96658.1 | pNP-a-L-Fucp | a-L-fucopyranosidase | 3.2.1.51 |
| **GH29** | *Wenyingzhuangia fucanilytica CZ1127* | ANW96659.1 | pNP-a-L-Fucp | a-L-fucopyranosidase | 3.2.1.51 |
| **GH29** | *Bacteroides thetaiotaomicron 7330* | ALJ41003.1 | pNP-a-L-Fucp | a-L-fucopyranosidase | 3.2.1.51 |
| **GH29** | *Arenibacter echinorum DSM 23522* | RAJ11632.1 | pNP-a-L-Fucp | a-L-fucopyranosidase | 3.2.1.51 |
| **GH29** | *Bacteroides stercoris CC31F* | EPH20525.1 | pNP-a-L-Fucp | a-L-fucopyranosidase | 3.2.1.51 |
| **GH29** | *Echinicola pacifica DSM 19836* | WP_229802408.1 | pNP-a-L-Fucp | a-L-fucopyranosidase | 3.2.1.51 |

| | | | | | |
|---|---|---|---|---|---|
| **GH29** | *Echinicola pacifica DSM 19836* | WP_044202327.1 | pNP-a-L-Fucp | a-L-fucopyranosidase | 3.2.1.51 |
| **GH29** | *Wenyingzhuangia fucanilytica CZ1127* | ANW96108.1 | pNP-a-L-Fucp | a-L-fucopyranosidase | 3.2.1.51 |
| **GH29** | *Wenyingzhuangia fucanilytica CZ1127* | ANW96113.1 | pNP-a-L-Fucp | a-L-fucopyranosidase | 3.2.1.51 |
| **GH29** | *Flagellimonas eckloniae DOKDO 007* | KQC29799.1 | pNP-a-L-Fucp | a-L-fucopyranosidase | 3.2.1.51 |
| **GH29** | *Flexithrix dorotheae DSM 6795* | WP_020529975.1 | pNP-a-L-Fucp | a-L-fucopyranosidase | 3.2.1.51 |
| **GH29** | *Zobellia galactanivorans DsijT* | CAZ94414.1 | pNP-a-L-Fucp | a-L-fucopyranosidase | 3.2.1.51 |
| **GH29** | *Zobellia galactanivorans DsijT* | CAZ94415.1 | pNP-a-L-Fucp | a-L-fucopyranosidase | 3.2.1.51 |
| **GH29** | *Saccharicrinis fermentans DSM 9555* | GAF01995.1 | pNP-a-L-Fucp | a-L-fucopyranosidase | 3.2.1.51 |
| **GH29** | *Niabella soli DSM 19437* | AHF15515.1 | pNP-a-L-Fucp | a-L-fucopyranosidase | 3.2.1.51 |
| **GH29** | *Niabella soli DSM 19437* | AHF15520.1 | pNP-a-L-Fucp | a-L-fucopyranosidase | 3.2.1.51 |
| **GH29** | *Niabella soli DSM 19437* | AHF15524.1 | pNP-a-L-Fucp | a-L-fucopyranosidase | 3.2.1.51 |
| **GH29** | *Cellulophaga baltica DSM 24729* | SDF23840.1 | Desulfated-oligo-porphyran | a-L-galactopyranosidase | 3.2.1. |
| **GH29** | *Cellulophaga baltica DSM 24729* | SDF23942.1 | Desulfated-oligo-porphyran | a-L-galactopyranosidase | 3.2.1. |
| **GH29*** | *Flavobacterium gilvum EM1308* | AOW10297.1 | aLFucp | a-L-fucopyranosidase | 3.2.1.51 |
| **GH29*** | *Flavobacterium gilvum EM1308* | AOW10301.1 | aLFucp | a-L-fucopyranosidase | 3.2.1.51 |
| **GH29*** | *Flavobacterium gilvum EM1308* | AOW10304.1 | aLFucp | a-L-fucopyranosidase | 3.2.1.51 |
| **GH29*** | *Flavobacterium gilvum EM1308* | AOW10314.1 | aLFucp | a-L-fucopyranosidase | 3.2.1.51 |
| **GH29*** | *Niabella ginsenosidivorans BS26* | ANH81215.1 | aLFucp | a-L-fucopyranosidase | 3.2.1.51 |
| **GH29*** | *Niabella ginsenosidivorans BS26* | ANH81213.1 | aLFucp | a-L-fucopyranosidase | 3.2.1.51 |
| **GH29\|CBM*** | *Flavobacterium gilvum EM1308* | AOW10315.1 | aLFucp, Oligos porphyran, Oligos agarose | a-L-fucopyranosidase, porphyranase, agarase | 3.2.1.51 |
| **GH30_2** | *Saccharicrinis fermentans DSM 9555* | GAF01993.1 | pNP-b-D-Xylp (fort) | b-D-xylopyranosidase | 3.2.1.37 |
| **GH31** | *Bacteroides ovatus ATCC 8483* | ALJ48502.1 | pNP-a-D-Xylp | a-D-xylopyranosidase | 3.2.1.177 |
| **GH31** | *Arenibacter echinorum DSM 23522* | RAJ11635.1 | pNP-a-D-Galp | a-D-galactopyranosidase | 3.2.1.22 |
| **GH31** | *Niabella soli DSM 19437* | AHF15526.1 | pNP-a-D-Galp | a-D-galactopyranosidase | 3.2.1.22 |
| **GH36** | *Bacteroides clarus YIT 12056* | EGF49700.1 | pNP-a-D-Galp | a-D-galactopyranosidase | 3.2.1.22 |
| **GH39** | *Saccharicrinis fermentans DSM 9555* | GAF01942.1 | pNP-b-D-Xylp | b-D-xylopyranosidase | 3.2.1.37 |
| **GH42** | *Flexithrix dorotheae DSM 6795* | WP_020529969.1 | pNP-b-D-Galp,pNP-a-L-Arap, pNP-a-L-Fucp | b-D-galactopyranosidase, a-L-arabinopyranosidase | 3.2.1.23 3.2.1.- |
| **GH43** | *Saccharicrinis fermentans DSM 9555* | GAF01996.1 | pNP-a-L-Fucp | a-L-fucopyranosidase | 3.2.1.51 |
| **GH50** | *Polaribacter sejongensis KCTC 23670* | AUC24144.1 | Oligo-agarose | Exo-b-agarase | 3.2.1 |
| **GH50** | *Wenyingzhuangia fucanilytica CZ1127* | ANW96656.1 | pNP-b-D-GlcA | b-D-glucuropyranosidase | 3.2.1.31 |
| **GH50** | *Saccharicrinis fermentans DSM 9555* | GAF03325.1 | pNP-b-D-GlcA | b-D-glucuropyranosidase | 3.2.1.31 |
| **GH50*** | *Pseudomonas aeruginosa VA-134* | ALP59072.1 | Laminaran, Curdlan, CM-Curdlan | endo-b-D-1,3-glucanase | 3.2.1.39 |
| **GH50*** | *Chthonomonas calidirosea T49* | CCW35650.1 | Laminaran, Curdlan, CM-Curdlan | endo-b-D-1,3-glucanase | 3.2.1.39 |
| **GH50*** | *Nostoc sp. HK-01 NIES-2109* | BBD60910.1 | pNP-b-D-Galp, Laminaran | b-D-galactopyranosidase | 3.2.1.23, 3.2.1.39 |
| **GH88** | *Bacteroides stercoris CC31F* | EPH20527.1 | Oligo-heparin, oligo-heparosan | D-4,5-unsaturated β-glucuronyl hydrolase | 3.2.1.- |
| **GH88** | *Saccharicrinis fermentans DSM 9555* | GAF01980.1 | Oligo-ascophyllan | D-4,5-unsaturated β-glucuronyl hydrolase | 3.2.1.- |
| **GH88** | *Bacteroides clarus YIT 12056* | EGF49697.1 | Oligo-heparin, oligo-heparosan | D-4,5-unsaturated β-glucuronyl hydrolase | 3.2.1.- |
| **GH92** | *Flagellimonas eckloniae DOKDO 007* | KQC29805.1 | pNP-a-D-Manp | a-D-mannopyranosidase | 3.2.1.24 |
| **GH95** | *Bacteroides stercoris CC31F* | EPH20507.1 | pNP-a-L-Fucp | a-L-fucopyranosidase | 3.2.1.51 |
| **GH95** | *Bacteroides clarus YIT 12056* | EGF49699.1 | pNP-a-L-Fucp | a-L-fucopyranosidase | 3.2.1.51 |
| **GH105** | *Niabella soli DSM 19437* | AHF15513.1 | Oligo-heparosan | D-4,5-unsaturated β-glucuronyl hydrolase | 3.2.1.- |
| **GH107** | *Wenyingzhuangia fucanilytica CZ1127* | ANW96098.1 | Fucan | endo-fucanase | 3.2.1.- |
| **GH117** | *Algoriphagus chordae DSM 19830* | PZX47569.1 | pNP-a-L-Fucp (weak) | a-L-fucopyranosidase | 3.2.1.51 |
| **GH117** | *Bacteroides cellulosilyticus WH2* | ALJ59764.1 | pNP-a-L-Fucp | a-L-fucopyranosidase | 3.2.1.51 |
| **GH117** | *Polaribacter reichenbachii KCTC 23969* | AUC19414.1 | Neo-oligo-agarose | a-L-galatopyranosidase | 3.2.1.159 |
| **GH117** | *Polaribacter reichenbachii KCTC 23969* | AUC19420.1 | Neo-oligo-agarose | a-L-galatopyranosidase | 3.2.1.159 |
| **GH117** | *Polaribacter reichenbachii KCTC 23969* | AUC19430.1 | Neo-oligo-agarose | a-L-galatopyranosidase | 3.2.1.159 |
| **GH117** | *Maribacter forsetii DSM 18668* | WP_209435193.1 | pNP-a-L-Fucp (Weak) | a-L-fucopyranosidase | 3.2.1.51 |
| **GH117** | *Cellulophaga baltica DSM 24729* | SDF24009.1 | Neo-oligo-agarose | a-L-galatopyranosidase | 3.2.1.159 |
| **GH127** | *Bacteroides ovatus CL02T12C04* | EIY64581.1 | Desulftated-neo-oligo-k-carrageenan | α-1,3-(3,6)-anhydro-D-galactosidase | 3.2.1.- |
| **GH150** | *Wenyingzhuangia fucanilytica CZ1127* | ANW96239.1 | l-carrageenan | l-carrageenase | 3.2.1.162 |

| | | | | | |
|---|---|---|---|---|---|
| **GH150** | *Wenyingzhuangia fucanilytica CZ1127* | ANW96242.1 | l-carrageenan | l-carrageenase | 3.2.1.162 |
| **GH167** | *Bacteroides ovatus CL02T12C04* | EIY64573.1 | Desulftated-neo-oligo-k-carrageenan | β-neocarrabiosidase | EC 3.2.1.- |
| **PL6** | *Niabella soli DSM 19437* | AHF15521.1 | Alginate, Poly-MG | alginate lyase | 4.2.2.- |
| **PL8_2** | *Bacteroides cellulosilyticus WH2* | ALJ59760.1 | Dermatan sulfate, Hyaluronic acid, Chondroitin sulfate | hyaluronate lyase, chondroitin sulfate lyase | 4.2.2.1, 4.2.2.20 |
| **PL8_2** | *Bacteroides cellulosilyticus WH2* | ALJ59767.1 | Dermatan sulfate, Hyaluronic acid, Chondroitin sulfate | hyaluronate lyase, chondroitin sulfate lyase | 4.2.2.1, 4.2.2.20 |
| **PL10** | *Cyclobacterium marinum DSM 745* | AEL26628.1 | Pectin Me 30%, Polygalacturonic acid (RGI) | | 4.2.2.- |
| **PL15** | *Bacteroides stercoris CC31F* | EPH20515.1 | Heparosan | heparosan lyase | 4.2.2.- |
| **PL15** | *Bacteroides stercoris CC31F* | EPH20517.1 | Heparosan | heparosan lyase | 4.2.2.- |
| **PL15** | *Bacteroides stercoris CC31F* | EPH20523.1 | Heparosan | heparosan lyase | 4.2.2.- |
| **PL15** | *Bacteroides clarus YIT 12056* | EGF49704.1 | Heparosan | heparosan lyase | 4.2.2.- |
| **PL15** | *Bacteroides clarus YIT 12056* | EGF49711.1 | Heparosan | heparosan lyase | 4.2.2.- |
| **PL15_2** | *Bacteroides stercoris CC31F* | EPH20506.1 | Heparin | heparin lyase | 4.2.2.7 |
| **PL15_2** | *Bacteroides stercoris CC31F* | EPH20511.1 | Heparosan | heparosan lyase | 4.2.2.- |
| **PL30** | *Bacteroides cellulosilyticus WH2* | ALJ59763.1 | Dermatan sulfate, Hyaluronic acid, Chondroitin sulfate | hyaluronate lyase, chondroitin sulfate lyase | 4.2.2.1, 4.2.2.20 |
| **PL33** | *Bacteroides stercoris CC31F* | EPH20528.1 | Heparosan | heparosan lyase | 4.2.2.- |
| **PL33_1** | *Bacteroides clarus YIT 12056* | EGF49696.1 | Heparosan | heparosan lyase | 4.2.2.- |
| **PL40** | *Saccharicrinis fermentans DSM 9555* | WP_211238157.1 | Saccorhysan | Saccorhysan | Saccorhysan |
| **PL43** | *Maribacter forsetii DSM 18668* | WP_157486574.1 | Saccorhysan | Saccorhysan | |
