## Supplementary Table 3 for "Screening for Polysaccharide Utilization Loci Targeting Marine Polysaccharides"

**Table S3:** List of additional targets selected to complete the coverage of the families GH50 and GH29 and to add new members to the newly created CAZy families (GHx446)

| | | Strain | Locus | Genbank |
|---|---|---|---|---|
| | Water | *Flavobacterium gilvum EM1308* | EM308_12710 | AOW10297.1 |
| | Water | *Flavobacterium gilvum EM1308* | EM308_12735 | AOW10301.1 |
| | Water | *Flavobacterium gilvum EM1308* | EM308_12750 | AOW10304.1 |
| | Water | *Flavobacterium gilvum EM1308* | EM308_12810 | AOW10314.1 |
| | Water | *Flavobacterium gilvum EM1308* | EM308_12815 | AOW10315.1 |
| | Compost | *Niabella ginsenosidivorans BS26* | A8C56_09675 | ANH81215.1 |
| | Compost | *Niabella ginsenosidivorans BS26* | A8C56_09660 | ANH81213.1 |
| | Water | *Flavobacterium gilvum EM1308* | EM308_12815 | AOW10315.1 |
| | Rumen | *Lachnospiraceae bacterium NLAE-zl-G231* | SAMN05216405_2716 | SFH34070.1 |
| | Human skin | *Pseudomonas aeruginosa VA-134* | ATC05_19450 | ALP59072.1 |
| | Soil | *Verrucomicrobium sp. GAS474* | SAMN05444156_1729 | SDU06021.1 |
| | Soil | *Chthonomonas calidirosea T49* | CCALI_01838 | CCW35650.1 |
| | Human oral | *Victivallales bacterium CCUG 44730* | C5Q97_17400 | AVM46385.1 |
| | Hot spring | *Thermosphaera aggregans DSM 11486* | Tagg_1112 | ADG91381.1 |
| | Marine | *Thermococcus guaymasensis DSM 11113* | X802_02960 | AJC71238.1 |
| | Sediment | *Sphaerochaeta pleomorpha str. Grapes* | SpiGrapes_1591 | AEV29398.1 |
| | Marine | *Phycisphaera mikurensis NBRC 102666* | PSMK_14930 | BAM03652.1 |
| | Freshwater | *Solidesulfovibrio magneticus RS-1* | DMR_13050 | BAH74796.1 |
| | Soil | *Nostoc sp. HK-01 NIES-2109* | NIES2109_37100 | BBD60910.1 |
| | Plant | *Bradyrhizobium diazoefficiens USDA 110* | bll4656 | BAC49921.1 |
| | Human gut | *Bifidobacterium longum BXY01* | GS08_09935 | WP_013141426.1 |
| | Human gut | *Bacteroides ovatus ATCC 8483* | Bovatus_03161 | ALJ47769.1 |
| | Saline sediment | *Sedimentisphaera salicampi ST-PulAB-D4* | STSP1_01769 | ARN57364.1 |
| | Soil | *Paenibacillus mucilaginosus K02* | B2K_10610 | AFH61168.1 |
| | Marine | *Maribacter polysiphoniae DSM 23514* | LX92DRAFT_00808 | MBD1262364.1 |
| | Plant associated | *Agrobacterium larrymoorei CFBP5473* | CFBP5473_20440 | QCJ00295.1 |
| | Plant associated | *Microbacterium hominis 01094* | IC744_13435 | QOC28385.1 |
| | Marine | *Rhodopirellula baltica SH 1* | RB13259 | CAD78028.1 |
| | Marine | *Zobellia galactanivorans DsijT* | ZOBELLIA_3623 | CAZ97761.1 |
| | Human gut | *Bacteroides xylanisolvens XB1A* | BXY_29870 | CBK68009.1 |
| | Human gut | *Bacteroides xylanisolvens XB1A* | BXY_44970 | CBK69371.1 |
| | Chicken gut | *Bacteroides gallinarum DSM 18171 = JCM 1* | C233DRAFT_01048 | WP_154656054.1 |
| | Chicken gut | *Bacteroides gallinarum DSM 18171 = JCM 1* | C233DRAFT_01049 | WP_128081889.1 |
| | Chicken gut | *Bacteroides gallinarum DSM 18171 = JCM 1* | C233DRAFT_01025 | WP_018666403.1 |
| | Human gut | *Bacteroides thetaiotaomicron VPI-5482* | BT0979 | AAO76086.1 |
| | Human gut | *Opitutaceae bacterium TAV5* | OPIT5_01565 | AHF89142.1 |
| | Periplaneta sp. hindgut | *Ereboglobus luteus Ho45* | CKA38_14220 | AWI10253.1 |
| | Periplaneta sp. hindgut | *Ereboglobus luteus Ho45* | CKA38_14295 | AWI10266.1 |
| | Plant associated | *Siphonobacter aquaeclarae DSM 21668* | Ga0070572_0687 | SDL30625.1 |
