## Supplementary Table 4 for "Screening for Polysaccharide Utilization Loci Targeting Marine Polysaccharides"

**Supplementary Table 4:** Attribution of the polysaccharide substrates of the PULs based on the demonstrated enzymes activities found in the PULs.

| PUL function | Locus | Genbank | CAZy - SulfAtlas | Substrate | Activity |
|---|---|---|---|---|---|
| **Undetermined** | ***SELECTED PUL 1 - Barnesiella viscericola DSM 18177*** **(Predicted PUL4)**<br>**GH16►GH50►◄ GH2 ◄ Sulf_1 ◄ GH20 ◄ Sulf_1 ◄ SusD ◄ SusC ◄ GH ◄ unk ◄ HTCS** | | | | **Undetermined** |
| | Barvi_0592 | AHF11965.1 | GH2 | pNP-b-D-Galp | b-D-galactopyranosidase |
| | Barvi_0594 | AHF11967.1 | GH20 | pNP-b-D-GlcNAc6S | b-D-N-Acetyl-6-sulfo-hexosaminidase |
| **Agarose** | ***SELECTED PUL 2 - Polaribacter sejongensis KCTC 23670*** **(Predicted PUL 34)**<br>**SusC►SusD►unk►◄ GH16 ◄ GH2 ◄ GH16 ◄ GH50 ◄ GH43 ◄ MFS ◄ GH117 ◄ EPI ◄ unk ◄ unk ◄ unk ◄ unk unk►◄ GH3 ◄ unk SusC►SusD►unk►** | | | | |
| | BTO15_18125 | AUC24144.1 | GH50 | Oligo-agarose | exo-b-agarase |
| | BTO15_18175 | AUC23902.1 | GH3 | pNP- b-D-Glcp, pNP-b-D-Xylp | b-D-glucopyranosidase, b-D-xylopyranosidase, |
| **Xylan** | ***SELECTED PUL 3 - Flavobacterium johnsoniae UW101*** **(Predicted PUL 12)**<br>**HTCS►SusC►SusD►GH10►unk►GH50►GH2►** | | | | |
| | Fjoh_2079 | ABQ05109.1 | GH10 | pNP-b-D-Cellobiose, Arabinoxylan | Arabinoxylanase |
| | Fjoh_2082 | ABQ05112.1 | GH2 | pNP-b-D-GlcA | b-D-glucuropyranosidase |
| **Undetermined** | ***SELECTED PUL 4 - Bacteroides ovatus ATCC 8483*** **(Pedicted PUL 39)**<br>**◄ GH50 ◄ GH50 ◄ GH2\|CBM57 ◄ unk ◄ SusD ◄ SusC ◄ HTCS** | | | | |
| | Bovatus_02309 | ALJ46942.1 | GH2\|CBM57 | pNP-b-D-GlcA | b-D-glucuropyranosidase |
| **Undetermined** | ***SELECTED PUL 5 - Wenyingzhuangia fucanilytica CZ1127*** **(Predicted PUL 14)**<br>**◄ Sulf_1 ◄ CE6 ◄ unk ◄ GH50 ◄ Sulf_1 ◄ HTCS ◄ GH29 ◄ GH29 ◄ GH92 ◄ SusD ◄ SusC unk►unk►GH78 ►** | | | | |
| | AXE80_10390 | ANW96656.1 | GH50 | pNP-b-D-GlcA | b-D-glucuropyranosidase |
| | AXE80_10405 | ANW96658.1 | GH29 | pNP-a-L-Fucp | a-L-fucopyranosidase |
| | AXE80_10410 | ANW96659.1 | GH29 | pNP-a-L-Fucp | a-L-fucopyranosidase |
| | AXE80_10435 | ANW96664.1 | unk (X425) | pNP-b-D-Galp | |
| **Undetermined** | ***SELECTED PUL 6 - Saccharicrinis fermentans DSM 9555*** **(Predicted PUL 37)**<br>**◄ GH2\|CBM57 ◄ GH136 ◄ CE6 ◄ unk ◄ SusD ◄ SusC ◄ SusD ◄ SusC ◄ Sulf_1 ◄ Sulf_1 ◄ GH50 ◄ Sulf_1 ◄ GH88 HTCS►◄ unk HTCS►GH43_24►Sulf_1\|GH16►Sulf_1►GH2►** | | | | |
| | CytfeDRAFT_4604 | GAF03325.1 | GH50 | pNP-b-D-GlcA | b-D-glucuropyranosidase |
| | CytfeDRAFT_4613 | WP_081736079.1 (G | GH2 | pNP-b-D-Galp | b-D-galactopyranosidase |
| **Undetermined** | ***SELECTED PUL 7 - Aquimarina agarivorans*** **(Cazymes cluster 1)**<br>**◄ unk ◄ unk ◄ Sulf_4 GH16►◄ GH50 ◄ unk** | | | | |
| **Undetermined** | ***SELECTED PUL 8 - Bacteroides cellulosilyticus WH2*** **(Predicted PUL 45)**<br>**◄ GH16\|Sulf_1 ◄ GH29 ◄ GH2 ◄ GH20 ◄ Sulf_1 ◄ SusD ◄ SusC HTCS►◄ GH159 ◄ unk ◄ GH146 ◄ GH50 ◄ GH159 ◄ unk ◄ SusD ◄ SusC ◄ SusD ◄ SusC HTCS►◄ Sulf_1 ◄ GH2** | | | | |
| | BcellWH2_02350 | ALJ59590.1 | GH2 | pNP-b-D-Galp | b-D-galactopyranosidase |

| | | | | | |
|---|---|---|---|---|---|
| | BcellWH2_02351 | ALJ59591.1 | GH20 | pNP-b-D-GlcNAc6S | b-D-N-Acetyl-6-sulfo-hexosaminidase |
| | BcellWH2_02368 | ALJ59608.1 | GH2 | b(1,3)-Galactan | b-D-1,3-galactopyranosidase |
| **Undetermined** | ***SELECTED PUL 9 - Bacteroides xylanisolvens NLAE-zl-H194* (Predited PUL 60)**<br>**◀ GH105 ◀ GH154 ◀ GH154\|Sulf_1 ◀ GH43_28 ◀ Sulf_1 ◀ GH88\|GH154 ◀ GH50 ◀ unk ◀ SusD ◀ SusC ◀ unk ◀ unk ◀ GH2 ◀ HTCS** | | | | |
| **Undetermined** | ***SELECTED PUL 10 - Bacteroides ovatus ATCC 8483* (Predicted PUL 81)**<br>**HTCS▶SusC▶SusD▶unk▶unk▶GH50▶unk▶GH▶GH31▶GH2\|CBM57▶CE6▶** | | | | |
| | Bovatus_03900 | ALJ48503.1 | GH2\|CBM | pNP-b-D-GlcA | b-D-glucuropyranosidase |
| | Bovatus_03899 | ALJ48502.1 | GH31_4 | pNP-a-D-Xylp | a-D-xylopyranosidase |
| **Undetermined** | **Bacteroides thetaiotaomicron 7330 (Predicted PUL 29)**<br>**◀ HTCS unk▶unk▶GH18▶SusC▶SusD▶GH▶Sulf_1▶GH20▶GH2▶GH29▶Sulf_1▶unk▶** | | | | |
| | Btheta7330_01433 | ALJ41001.1 | GH20 | pNP-b-D-GlcNAc6S | b-D-N-Acetyl-6-sulfo-hexosaminidase |
| | Btheta7330_01434 | ALJ41002.1 | GH2 | pNP-b-D-Galp | b-D-galactopyranosidase |
| | Btheta7330_01435 | ALJ41003.1 | GH29 | pNP-a-L-Fucp | a-L-fucopyranosidase |
| **Undetermined** | ***Parabacteroides gordonii DSM 23371* (Predicted PUL 51)**<br>**AraC▶◀ unk ◀ Sulf_1 ◀ GH29 ◀ Sulf_1 ◀ unk ◀ GH117\|GH117 ◀ SusD ◀ SusC ◀ Anti-σ ECF-σ▶** | | | | |
| | F592DRAFT_02163 | KKB57913.1 | GH20\|CBM | pNP-b-D-GlcNAc, pNP-b-D-GalNAc | b-D-N-Acetyl-hexosaminidase |
| **Undetermined** | **Algoriphagus chordae DSM 19830 (Predicted PUL 34)**<br>**◀ Sulf_1 ◀ Sulf_1 unk▶Sulf_1▶unk▶Sulf_1▶GH117▶unk▶GH117▶SusC▶SusD▶unk▶Sulf_1▶◀ GH95** | | | | |
| | LV85DRAFT_03918 | PZX47569.1 | GH117 | pNP-a-L-Fucp (weak) | a-L-fucopyranosidase |
| **Undetermined** | ***Bacteroides dorei DSM 17855* (Cazymes cluster 3)**<br>**GH29▶unk▶unk▶Sulf_1▶GH29▶Sulf_1▶GH20▶GH117▶** | | | | |
| | BACDOR_01585 | EEB26054.1 | **S1_46** | pNP-sulfate | Aryl-sulfatase |
| | BACDOR_01586 | EEB26055.1 | GH20 | pNP-b-D-GlcNAc6S, pNP-b-D-GlcNAc, pNP-b-D-GalNA | b-D-N-Acetyl-6-sulfo-hexosaminidase |
| **Undetermined** | ***Chitinophaga pinensis DSM 2588* (Predicted PUL 70)**<br>**SusC▶SusD▶◀ GH20 ◀ unk GH127▶Sulf_4▶unk▶** | | | | |
| | Cpin_4994 | ACU62427.1 | GH20 | pNP-b-D-GlcNAc, pNP-b-D-GalNAc | b-D-N-Acetyl-hexosaminidase |
| **Chondroitin sulfate, dermatan, hyaluronan** | ***Bacteroides cellulosilyticus WH2* (Predicted PUL 43)**<br>**HTCS▶Sulf_1▶SusC▶SusD▶PL8_2▶SusC▶SusD▶◀ PL30 GH117▶unk▶◀ Sulf_1 ◀ PL8_2 ◀ unk** | | | | |
| | BcellWH2_02521 | ALJ59760.1 | PL8_2 | Dermatan sulfate, Hyaluronic acid, Chondroitin sulfate | hyaluronate lyase, chondroitin sulfate lyase |
| | BcellWH2_02524 | ALJ59763.1 | PL30 | Dermatan sulfate, Hyaluronic acid, Chondroitin sulfate | hyaluronate lyase, chondroitin sulfate lyase |
| | BcellWH2_02525 | ALJ59764.1 | GH117 | pNP-a-L-Fucp | a-L-fucopyranosidase |
| | BcellWH2_02527 | ALJ59766.1 | **S1_9** | pNP-sulfate | Aryl-sulfatase |
| | BcellWH2_02528 | ALJ59767.1 | PL8_2 | Dermatan sulfate, Hyaluronic acid, Chondroitin sulfate | hyaluronate lyase, chondroitin sulfate lyase |
| **Undetermined** | ***Arenibacter echinorum DSM 23522* (Cazymes cluster 4)**<br>**unk▶unk▶GH▶GH33▶GH▶unk▶unk▶◀ unk ◀ unk ◀ unk ◀ MFS ◀ unk ◀ GH29\|GH117 ◀ Sulf_1 ◀ GH29 ◀ unk ◀ unk ◀ GH31 ◀ unk ◀ Sulf_1 ◀ GH ◀ GH29** | | | | |

| | | | | | |
|---|---|---|---|---|---|
| | LV92DRAFT_02563 | RAJ11632.1 | GH29 | pNP-a-L-Fucp | a-L-fucopyranosidase |
| | LV92DRAFT_02566 | RAJ11635.1 | GH31_14 | pNP-a-D-Galp | a-D-galactopyranosidase |
| **Agarose, porphyran** | ***Polaribacter reichenbachii KCTC 23969* (Cazymes cluster 5)**<br>**◀ GntR ◀ Sulf_1 ◀ Sulf_1 ◀ GH117 ◀ GH16 ◀ GH117 ◀ GH2 ◀ GH2 ◀ unk ◀ unk unk▶◀ GH117 ◀ Sulf_1 ◀ Sulf_1 ◀ GH28 ◀ Sulf_1 ◀ Sulf_1 ◀ GH2 ◀ Sulf_1 ◀ GH16 ◀ GH117 ◀ GH29 ◀ GH117 ◀ GH2 ◀ GH2 unk▶◀ GH28 ◀ unk ◀ Sulf_1 ◀ GH29 HTCS▶** | | | | |
| | BTO17_12250 | AUC19414.1 | GH117 | Neo-oligo-agarose | a-L-galatopyranosidase |
| | BTO17_12280 | AUC19420.1 | GH117 | Neo-oligo-agarose | a-L-galatopyranosidase |
| | BTO17_12310 | AUC20527.1 | GH2 | pNP-b-D-Galp | b-D-galactopyranosidase |
| | BTO17_12320 | AUC19427.1 | GH16_11 | Porphyran | Porphyranase |
| | BTO17_12335 | AUC19430.1 | GH117 | Neo-oligo-agarose | a-L-galatopyranosidase |
| | BTO17_12355 | AUC19434.1 | GH28 | pNP-a-D-GlcA | a-D-glucuronidase |
| **Undetermined** | ***Algibacter lectus JCM 19300* (Predicted PUL 23)**<br>**◀ SusD ◀ unk ◀ SusC Sulf_1▶unk▶◀ GH117 ◀ GH95 ◀ unk ◀ GH117 ◀ GH117 ◀ GH2 ◀ GH117 ◀ unk ◀ GH97 ◀ Sulf_1 ◀ CBM47 ◀ Sulf_1 ◀ Sulf_1 SusC▶SusD▶SusC▶SusD▶unk▶** | | | | |
| | Ga0062136_116110 | GAL64206.1 | GH2 | pNP-a-D-GlcA | b-D-glucuropyranosidase |
| **Polygalacturonan** | ***Bacteroides ovatus NLAE-zl-C57* (Predicted PUL 19)**<br>**◀ GH2 ◀ PL11_1 GH28▶◀ unk ◀ unk ◀ GH117 ◀ unk ◀ HTCS SusC▶SusD▶unk▶unk▶unk▶◀ PL11_1 PL9_1▶unk▶** | | | | |
| | Ga0104390_100624 | SDH45426.1 | GH28 | Polygalacturonic acid (RG I), pNP-a-D-GlcA | a-D-1,4-galacturonase |
| **Undetermined** | ***Sphingobacterium thalpophilum DSM 11723* (Predicted PUL 13)**<br>**CE12▶unk▶SusC▶SusD▶unk▶unk▶SusC▶SusD▶GH51▶GH115▶GH43_18▶unk▶GH2▶unk▶◀ GH28 GH28▶PL11_1▶CE12▶CE12▶Sulf_1▶GH105 ▶GH2▶unk▶GH106▶SusC▶SusD▶GH117▶GH28▶unk▶unk▶SusC▶SusD▶unk▶PL1_2▶CE8▶PL1_2▶SusC▶SusD▶GH105▶GntR▶** | | | | |
| **Undetermined** | ***Maribacter forsetii DSM 18668* (Predicted PUL 13)**<br>**◀ GH95 ◀ Sulf_1 ◀ PL40 ◀ unk ◀ GH117 ◀ PL33_2 ◀ unk ◀ GH29 ◀ GH ◀ Pept_SC ◀ PL40 ◀ Sulf_1 ◀ unk ◀ MFS ◀ GH88 SusC▶SusD▶◀ HTCS** | | | | |
| | P177DRAFT_02789 | WP_209435193.1 | GH117 | pNP-a-L-Fucp (Weak) | a-L-fucopyranosidase |
| **Undetermined** | ***Saccharicrinis fermentans DSM 9555* (Cazymes cluster 6)**<br>**◀ GH ◀ PL40 ◀ unk ◀ unk ◀ unk ◀ GH39 ◀ unk ◀ CBM6 ◀ GH117 ◀ unk ◀ Sulf_1 ◀ GH2 ◀ unk ◀ Sulf_1 ◀ unk ◀ unk ◀ GH97 ◀ Sulf_1 HTCS▶◀ GH29** | | | | |
| | CytfeDRAFT_1523 | GAF01942.1 | GH39 | pNP-b-D-Xylp | b-D-xylopyranosidase |
| **Undetermined** | ***Flammeovirga sp. MY04* (Predicted PUL 37)**<br>**◀ GH117 Sulf_1▶Sulf_1▶Sulf_1▶GH2▶Sulf_1▶Sulf_1▶GH136\|CBM16\|CBM16▶◀ GH95 ◀ Sulf_1 ◀ GH16 ◀ GH2 ◀ GH128 ◀ SusD ◀ SusC Sulf_1▶HTCS▶GH136▶unk▶◀ GH136 ◀ GH136 ◀ Sulf_1 ◀ GH2 ◀ GH2 ◀ GH136 ◀ Sulf_1 ◀ SusD ◀ SusC** | | | | |
| | MY04_5120 | ANQ52455.1 | GH2 | pNP-b-D-Galp | b-D-galactopyranosidase |
| **Undetermined** | ***Parabacteroides johnsonii CL02T12C29* (Predicted PUL 23)**<br>**ECF-σ▶Anti-σ▶SusC▶SusC▶SusD▶GH16▶unk▶unk▶GH97▶GH106▶GH117\|GH43_24▶Sulf_1▶unk▶unk▶unk▶CE▶GH105▶** | | | | |

| | | | | | |
|---|---|---|---|---|---|
| **Carrageenan** | ***Tamlana sp. UJ94* (Predicted PUL 13)**<br>**SusC▶SusD▶Sulf_1▶GH127▶unk▶MFS▶unk▶Sulf_1▶GH16▶unk▶unk▶GH167▶GH16▶** | | | | |
| | C1A40_08425 | AUS05489.1 | GH16_13 | a-carrageenan | a-carrageenase |
| | C1A40_08445 | AUS05492.1 | GH16_17 | k-carrageenan | k-carrageenase |
| **Carrageenan** | ***Wenyingzhuangia fucanilytica CZ1127* (Predicted PUL 9)**<br>**◀ GH167 ◀ GH167 ◀ GH110 ◀ unk ◀ unk ◀ Sulf_1 ◀ unk ◀ GH150 ◀ unk ◀ unk ◀ GH150 ◀ unk ◀ unk ◀ SusD ◀ SusC ◀ HTCS ◀ GH127 ◀ GH2** | | | | |
| | AXE80_08085 | ANW96239.1 | GH150 | l-carrageenan | l-carrageenase |
| | AXE80_08100 | ANW96242.1 | GH150 | l-carrageenan | l-carrageenase |
| | AXE80_08135 | ANW96249.1 | GH2 | pNP-b-D-Galp | b-D-galactopyranosidase |
| **Carrageenan** | ***Bacteroides ovatus CL02T12C04* (Predicted PUL 25)**<br>**◀ GntR ◀ Sulf_1 unk▶ ◀ unk ◀ unk ◀ unk ◀ unk ◀ unk ◀ GH16 ◀ unk ◀ unk ◀ unk ◀ SusD ◀ SusC ◀ GH16 ◀ unk ◀ GH2 ◀ GH167 ◀ GH2 ◀ Sulf_1 ◀ unk ◀ unk ◀ unk ◀ Sulf_1 ◀ Sulf_1 ◀ GH127 ◀ unk ◀ unk** | | | | |
| | HMPREF1069_0208 | EIY64557.1 | **S1_30** | Exo-4S-i-carrageenan sulfatase | |
| | HMPREF1069_0209 | EIY64570.1 | GH16_17 | k-carrageenan, i-carrageenan | k-carrageenase, i-carrageenase |
| | HMPREF1069_0210 | EIY64572.1 | GH2 | pNP-b-D-Galp | b-D-galactopyranosidase |
| | HMPREF1069_0210 | EIY64573.1 | GH167 | Desulftated-neo-oligo-k-carrageenan | β-neocarrabiosidase |
| | HMPREF1069_0210 | EIY64574.1 | GH2 | pNP-b-D-Galp | b-D-galactopyranosidase |
| | HMPREF1069_0210 | EIY64575.1 | **S1_30** | neo-oligo-i-carrageenan | Exo-4S-i-carrageenan sulfatase |
| | HMPREF1069_0210 | EIY64579.1 | **S1_16** | neo-oligo-k-carrageenan | Exo-4S-k-carrageenan sulfatase |
| | HMPREF1069_0210 | EIY64580.1 | **S1_81** | neo-oligo-a-carrageenan | Exo-2S-a-carrageenan sulfatase |
| | HMPREF1069_0211 | EIY64581.1 | GH127 | Desulftated-neo-oligo-k-carrageenan | α-1,3-(3,6)-anhydro-D-galactosidase |
| **Polygalacturonan** | ***Cyclobacterium marinum DSM 745* (Predicted PUL 24)**<br>**unk▶PL10_1▶PL1_2▶GH127▶unk▶ ◀ unk Sulf_1▶ ◀ unk ◀ unk ◀ Sulf_1 ◀ SusD ◀ SusC** | | | | |
| | Cycma_2892 | AEL26628.1 | PL10_1 | Pectin Me 30%, Polygalacturonic acid (RGI) | |
| **Undetermined** | ***Nonlabens sp. Hel1_33_55* (Predicted PUL 1)**<br>**HTCS▶SusC▶SusD▶GH3▶Sulf_1▶Sulf_1▶Sulf_1▶unk▶GH29▶Sulf_1▶unk▶Sulf_1▶GH10▶unk▶Sulf_1▶PL▶Sulf_1▶Sulf_1▶unk▶unk ▶** | | | | |
| | SAMN05192588_064 | SCX99618.1 | GH3 | pNP-b-D-Glcp, pNP-b-D-cellobiosides | b-D-glucopyranosidase |
| **Heparin, heparosan** | ***Bacteroides stercoris CC31F* (Predicted PUL 28)**<br>**Sulf_1▶PL15_2▶GH95▶ ◀ unk SusC▶SusD▶PL15_2▶unk▶unk▶unk▶PL15▶HTCS▶PL15▶SusC▶SusD▶unk▶ ◀ unk unk▶PL15▶ ◀ unk GH29▶unk▶GH88▶PL33_1▶** | | | | |
| | HMPREF1181_0172 | EPH20505.1 | **S1_9** | Exo-2S-Oligo-heparin sulfatase | △4,5-hexuronate-2-O-sulfate 2-O-sulfohydrolase |
| | HMPREF1181_0172 | EPH20506.1 | PL15_2 | Heparin | heparin lyase |
| | HMPREF1181_0172 | EPH20507.1 | GH95 | pNP-a-L-Fucp | a-L-fucopyranosidase |

| | | | | | |
|---|---|---|---|---|---|
| | HMPREF1181_01727 | EPH20511.1 | PL15_2 | Heparosan | heparosan lyase |
| | HMPREF1181_01730 | EPH20514.1 | unk | Heparin | heparin lyase |
| | HMPREF1181_01731 | EPH20515.1 | PL15 | Heparosan | heparosan lyase |
| | HMPREF1181_01733 | EPH20517.1 | PL15 | Heparosan | heparosan lyase |
| | HMPREF1181_01739 | EPH20523.1 | PL15 | Heparosan | heparosan lyase |
| | HMPREF1181_01741 | EPH20525.1 | GH29 | pNP-a-L-Fucp | a-L-fucopyranosidase |
| | HMPREF1181_01743 | EPH20527.1 | GH88 | Oligo-heparin, oligo-heparosan | D-4,5-unsaturated β-glucuronyl hydrolase |
| | HMPREF1181_01744 | EPH20528.1 | PL33_1 | Heparosan | heparosan lyase |
| **Undetermined** | ***Echinicola pacifica DSM 19836* (Predicted PUL 33)**<br>**◄ SusD ◄ SusC ◄ Sulf_1 ◄ unk ◄ Sulf_1 ◄ Sulf_1 ◄ GH29 ◄ Sulf_1 ◄ unk ◄ Sulf_1 ◄ unk ◄ GH29 ◄ unk ◄ GH ◄ Sulf_1 ◄ Sulf_1 ◄ Sulf_1 ◄ Sulf_1 ◄ GH29 ◄ GH95 ◄ GH ◄ SusD ◄ SusC** | | | | |
| | B050DRAFT_01789 | WP_229802408.1 | GH29 | pNP-a-L-Fucp | a-L-fucopyranosidase |
| | B050DRAFT_01801 | WP_044202327.1 | GH29 | pNP-a-L-Fucp | a-L-fucopyranosidase |
| **Fucan** | ***Wenyingzhuangia fucanilytica CZ1127* (Predicted PUL 8)**<br>**◄ GH ◄ GH107 ◄ GH107 ◄ unk ◄ unk ◄ unk ◄ unk ◄ Sulf_1 ◄ Sulf_1 ◄ unk ◄ GH29 ◄ GH95 ◄ unk ◄ unk ◄ GH ◄ GH ◄ GH29 ◄ Sulf_1 ◄ SusD ◄ SusC ◄ Sulf_1 ◄ unk ◄ GH29 ◄ Sulf_1 ◄ GH107 ◄ GH107** | | | | |
| | AXE80_07310 | ANW96098.1 | GH107 | Fucan | endo-fucanase |
| | AXE80_07380 | ANW96108.1 | GH29 | pNP-a-L-Fucp | a-L-fucopyranosidase |
| | AXE80_07410 | ANW96113.1 | GH29 | pNP-a-L-Fucp | a-L-fucopyranosidase |
| **Undetermined** | ***Flagellimonas eckloniae DOKDO 007* (Predicted PUL 6)**<br>**◄ GntR ◄ PL7 ◄ unk ◄ unk ◄ SusD ◄ SusC ◄ unk ◄ PL17_2 ◄ PL6_1\|PL6_1 unk► ◄ GH92 ◄ unk ◄ CBM9 ◄ GH20 ◄ GH92 ◄ GH20 ◄ GH29 ◄ Sulf_1 ◄ GH92 ◄ GH3 ◄ Sulf_1 ◄ GH95 ◄ Sulf_1 ◄ Sulf_1 ◄ GH29 ◄ Sulf_1 ◄ Sulf_1 ◄ GH130 ◄ unk ◄ GH92 ◄ SusD ◄ SusC** | | | | |
| | AAY42_07780 | KQC29797.1 | GH20 | pNP-b-D-GlcNAc, pNP-b-D-GalNAc | b-D-N-Acetyl-hexosaminidase |
| | AAY42_07790 | KQC31661.1 | GH20 | pNP-b-D-GlcNAc, pNP-b-D-GalNAc | b-D-N-Acetyl-hexosaminidase |
| | AAY42_07795 | KQC29799.1 | GH29 | pNP-a-L-Fucp | a-L-fucopyranosidase |
| | AAY42_07860 | KQC29805.1 | GH92 | pNP-a-D-Manp | a-D-mannopyranosidase |
| **Undetermined** | ***Flexithrix dorotheae DSM 6795* (Predicted PUL 40)**<br>**unk►unk►unk►Sulf_4► ◄ unk ◄ GH28 ◄ Sulf_1 ◄ Sulf_1 ◄ Sulf_1 ◄ unk ◄ Sulf_1 ◄ GH ◄ unk ◄ unk ◄ GH42 ◄ unk ◄ SusD ◄ SusC HTCS► ◄ GH29 ◄ PL29 ◄ GH88 ◄ unk ◄ unk ◄ GH** | | | | |
| | A3EMDRAFT_05982 | WP_020529969.1 | GH42 | pNP-b-D-Galp,pNP-a-L-Arap, pNP-a-L-Fucp | b-D-galactopyranosidase, a-L-arabinopyranosidase |
| | A3EMDRAFT_05987 | WP_020529975.1 | GH29 | pNP-a-L-Fucp | a-L-fucopyranosidase |
| | ***Zobellia galactanivorans DsijT* (Predicted PUL 9)** | | | | |

| | | | | | |
|---|---|---|---|---|---|
| Undetermined | ◀ GH ◀ unk ◀ Sulf_1 ◀ PL\|CBM16 ◀ GH29\|CBM47 ◀ GH29\|CBM47 ◀ unk ◀ GH29 ◀ GH29 ◀ GH95 ◀ GH3 ◀ GH29 ◀ GH29\|CBM51 ◀ GH3 HTCS▶ ◀ unk ◀ SusD ◀ SusC ◀ SusD ◀ SusC ◀ unk ◀ GH97 ◀ SusD ◀ SusC ◀ Anti-σ ◀ ECF-σ | | | | |
| | ZOBELLIA_334 | CAZ94407.1 | GH_New2 | pNP-a-D-Galp | a-D-galactosidase |
| | ZOBELLIA_341 | CAZ94414.1 | GH29 | pNP-a-L-Fucp | a-L-fucopyranosidase |
| | ZOBELLIA_342 | CAZ94415.1 | GH29 | pNP-a-L-Fucp | a-L-fucopyranosidase |
| Saccorhysan | ***Saccharicrinis fermentans DSM 9555* (Predicted PUL 14)**<br>**◀ PL40 ◀ PL40 ◀ GH43 ◀ GH29 ◀ unk ◀ GH30_2 ◀ unk ◀ GH97 ◀ SusD ◀ SusC ◀ unk ◀ unk ◀ SusD ◀ SusC ◀ unk ◀ HTCS ◀ GH88 ◀ unk ◀ Sulf_1 ◀ CBM6\|CBM6\|CBM6\|CBM6 ◀ PL ◀ SusD ◀ SusC** | | | | |
| | CytfeDRAFT_1478 | WP_211238157.1 (G | PL40 | Saccorhysan | Saccorhysan lyase |
| | CytfeDRAFT_1479 | GAF01996.1 | GH43 | pNP-a-L-Fucp | a-L-fucopyranosidase |
| | CytfeDRAFT_1480 | GAF01995.1 | GH29 | pNP-a-L-Fucp | a-L-fucopyranosidase |
| | CytfeDRAFT_1482 | GAF01993.1 | GH30_2 | pNP-b-D-Xylp | b-D-xylopyranosidase |
| | CytfeDRAFT_1493 | GAF01980.1 | GH88 | Oligo-ascophyllan | D-4,5-unsaturated β-glucuronyl hydrolase |
| Alginate | ***Niabella soli DSM 19437* (Cazymes cluster 5)**<br>**◀ GH28 ◀ GH29 ◀ PL6_2 ◀ unk ◀ GH88 ◀ GH29 ◀ PL38 ◀ GH31 ◀ GH95 ◀ GH2 ◀ unk ◀ SusD ◀ SusC GH28 ▶** | | | | |
| | Niaso_1999 | AHF15512.1 | PL_New1 | Alginate, Polymannuronic acid | alginate lyase |
| | Niaso_2000 | AHF15513.1 | GH105 | Oligo-heparosan | D-4,5-unsaturated β-glucuronyl hydrolase |
| | Niaso_2002 | AHF15515.1 | GH29 | pNP-a-L-Fucp | a-L-fucopyranosidase |
| | Niaso_2007 | AHF15520.1 | GH29 | pNP-a-L-Fucp | a-L-fucopyranosidase |
| | Niaso_2008 | AHF15521.1 | PL6_2 | Alginate, Poly-MG | alginate lyase |
| | Niaso_2011 | AHF15524.1 | GH29 | pNP-a-L-Fucp | a-L-fucopyranosidase |
| | Niaso_2013 | AHF15526.1 | GH31 | pNP-a-D-Galp | a-D-galactopyranosidase |
| | Niaso_2015 | AHF15528.1 | GH2 | pNP-b-D-GlcA | b-D-galactopyranosidase, a-L-arabinopyranosidase |
| Heparosan | ***Bacteroides clarus YIT 12056* (Predicted PUL 41)**<br>**◀ PL33_1 ◀ GH88 ◀ unk ◀ GH95 ◀ GH36 ◀ GH29 ◀ SusD ◀ SusC ◀ PL15 ◀ GH95 ◀ HTCS ◀ HTCS ◀ GH88 ◀ Sulf_1 ◀ unk ◀ PL15** | | | | |
| | HMPREF9445_0306[illegible] | EGF49696.1 | PL33_1 | Heparosan | heparosan lyase |
| | HMPREF9445_0306[illegible] | EGF49697.1 | GH88 | Oligo-heparin, oligo-heparosan | D-4,5-unsaturated β-glucuronyl hydrolase |
| | HMPREF9445_0307[illegible] | EGF49699.1 | GH95 | pNP-a-L-Fucp | a-L-fucopyranosidase |
| | HMPREF9445_0307[illegible] | EGF49700.1 | GH36 | pNP-a-D-Galp | a-D-galactopyranosidase |
| | HMPREF9445_0307[illegible] | EGF49704.1 | PL15 | Heparosan | heparosan lyase |
| | HMPREF9445_0308[illegible] | EGF49711.1 | PL15 | Heparosan | heparosan lyase |
| | ***Cellulophaga baltica DSM 24729* (Predicted PUL 22)** | | | | |

| | | | | | |
|---|---|---|---|---|---|
| **Agarose, porphyran** | **unk ▶ unk ▶ unk ▶ unk ▶ GH29 ▶ SusC ▶ SusD ▶ unk ▶ GH16 ▶ Sulf_1 ▶ unk ▶ GH29 ▶ unk ▶ GH16 ▶ Sulf_1 ▶ GH29 ▶ GH29 ▶ ◀ unk ◀ Sulf_1 ◀ GH117 ◀ GH28 ◀ GH16 ◀ Sulf_1 ◀ GH28 ◀ Sulf_1 Sulf_1 ▶ ◀ GH2 ◀ GH16 ◀ Sulf_1 ◀ Sulf_1 ◀ Sulf_1 ◀ GH2 ◀ Sulf_1 ◀ Sulf_1 ◀ GH2 ◀ Sulf_1 ◀ HTCS** | | | | |
| | Ga0070226_10974 | SDF23778.1 | GH16 | Porphyran, agarose | b-agarase |
| | Ga0070226_10976 | SDF23816.1 | GH_new1 | Porphyran, agarose | b-agarase |
| | Ga0070226_10977 | SDF23840.1 | GH29 | Desulfated-oligo-porphyran | a-L-galactopyranosidase |
| | Ga0070226_10982 | SDF23942.1 | GH29 | Desulfated-oligo-porphyran | a-L-galactopyranosidase |
| | Ga0070226_10985 | SDF24009.1 | GH117 | Neo-oligo-agarose | a-L-galatopyranosidase |
| | Ga0070226_10987 | SDF24056.1 | GH16 | Porphyran | porphyranase |
| | Ga0070226_10992 | SDF24160.1 | GH2 | pNP-b-D-Galp | b-D-galactopyranosidase |
| | Ga0070226_109100 | SDF24334.1 | GH2 | Porphyran, agarose | b-agarase (ou exo-b-neo-agarobiosidase) |
| **Porphyran** | ***Reichenbachiella agariperforans DSM 26134* (Predicted PUL 21)** **SusC ▶ SusD ▶ SusC ▶ SusD ▶ unk ▶ unk ▶ GH29 ▶ Sulf_1 ▶ Sulf_1 ▶ PL ▶ GH28 ▶ GH28 ▶ Sulf_1 ▶ GH2 ▶ GH29 ▶ GH29 ▶ ◀ Sulf_1 unk ▶ ◀ GH2 ◀ GH16 ◀ Sulf_1 ◀ Sulf_1** | | | | |
| | Ga0070003_10798 | SHK67940.1 | GH2 | pNP-b-D-Galp | b-D-galactopyranosidase, b-D-fucopyranosidase |
| | Ga0070003_10799 | SHK67965.1 | GH16 | Porphyran, oligo-porphyran | porphyranase |
