## Supplementary material for "Screening for Polysaccharide Utilization Loci Targeting Marine Polysaccharides": Captions Supplementary figures

### Captions for supplementary figures

**Figure S1:** Phylogenetic tree calculated from members of the GH29 family located in the PUL. The composition of the PUL, including other CAZymes and sulfatases, as well as the corresponding organism, are indicated on each branch. Sulfatases are coloured orange, GH50 and GH117 studied here are coloured green, and some other CAZymes (e.g., GH16) are coloured in blue, to highlight the conserved composition within clades.

**Figure S2:** Phylogenetic tree calculated from members of the GH50 family located in the PUL. The composition of the PUL, including other CAZymes and sulfatases, as well as the corresponding organism, are indicated on each branch. Sulfatases are coloured orange, GH29 and GH117 studied here are coloured green, and some other CAZymes (e.g., GH16) are coloured in blue, to highlight the conserved composition within clades.

**Figure S3:** Phylogenetic tree calculated from members of the GH117 family located in the PUL. The composition of the PUL, including other CAZymes and sulfatases, as well as the corresponding organism, are indicated on each branch. Sulfatases are coloured orange, GH29 and GH50 studied here are coloured green, and some other CAZymes (e.g., GH16) are coloured in blue, to highlight the conserved composition within clades.

**Table S1**: List of the selected PULs. The selected proteins listed glycoside hydrolases (GH), polysaccharides lyases (PL), sulfatases (S) and unclassified proteins (unk). * Not studied

**Table S2**: Activity screening of the enzymes grouped in 31 established CAZy families and 5 SulfAtlas sub-families. * Not in PUL.
